# Characterizing the Solution Space of Migration Histories of Metastatic Cancers with MACH2

**DOI:** 10.1101/2024.11.19.624301

**Authors:** Mrinmoy S. Roddur, Vikram Ramavarapu, Abigail Bunkum, Ariana Huebner, Roman Mineyev, Nicholas McGranahan, Simone Zaccaria, Mohammed El-Kebir

## Abstract

Understanding the migration history of cancer cells is essential for advancing metastasis research and developing therapies. Existing migration history inference methods often rely on parsimony criteria such as minimizing migrations, comigrations, and seeding locations. Importantly, existing methods either yield a single optimal solution or are probabilistic algorithms without guarantees on optimality nor comprehensiveness of the returned solution space. As such, current methods are unable to capture the full extent of uncertainty inherent to the data. To address these limitations, we introduce MACH2, a method that systematically enumerates all plausible migration histories by exactly solving the Parsimonious Migration History with Tree Refinement (PMH-TR). In addition to the migration and comigration criteria, MACH2 employs a novel parsimony criterion that minimizes the number of clones unobserved in their inferred location of origin. MACH2 allows one to specify both the order and the set of criteria to include during optimization, allowing users to adapt the model to specific analysis needs. MACH2 also includes a summary graph to identify high-confidence migrations. Finally, we introduce MACH2-viz, an interactive webtool for visualizing and exploring MACH2 solution spaces. Using simulated tumors with known ground truth, we show that MACH2, especially the version that prioritizes the new unobserved clone criterion, outperforms existing methods. On real data, MACH2 detects extensive uncertainty in non-small cell lung, ovarian, and prostate cancers, and infers migration histories consistent with orthogonal experimental data.

## 1 Introduction

Metastasis, the spread of cancer cells from the primary tumor to distant anatomical locations, is the main cause of cancer-related deaths [1, 2]. Reconstructing the precise pattern of spread of metastatic cancers helps uncover metastatic pathways, identify treatment targets, and develop precision medicine [3–11]. Since cancer cells in different locations evolve independently, one way to trace the migration history of a tumor is by analyzing the evolutionary relationships of its cells across metastatic sites. To that end, researchers run computational methods [12–18] to identify *clones* — groups of cells with the same set of mutations — from sequencing data of primary and metastatic biopsies as well as their evolutionary history in the form of a *clonal tree*, where each node represents a clone labeled by the anatomical locations in which it was observed (Fig. 1a, b). Recent migration history inference methods further analyze this clonal tree to determine each clone’s location of origin and identify migrations indicated by parent-child pairs with distinct origins via one or more parsimony criteria [19–21]. The key principle underlying these methods is that metastatic spread between locations is a rare event, and therefore inferring migration histories with as few metastasis events is preferable over more complex histories with many such events.

**Figure 1:**
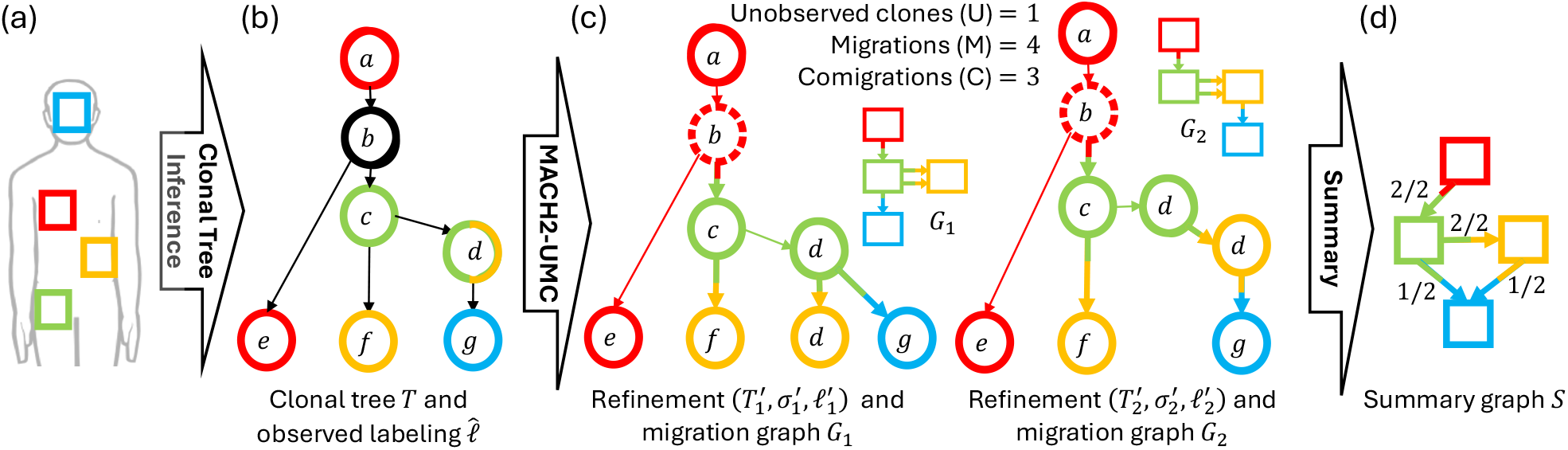
Overview of MACH2. (a) From sequencing data of matched primary and metastatic biopsies, (b) we infer a clonal tree *T* with observed location labeling 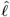 (colors), describing the evolutionary history of the tumor and the clonal composition of each biopsy, respectively. Here, clone ‘b’ was not observed in any location, whereas clone ‘d’ was observed in the green and yellow locations. (c) MACH2 in UMC mode enumerates all optimal Parsimonious Migration History with Tree Refinement (PMH-TR) solutions composed of refinements of 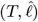, lexicographically minimizing the number of unobserved clones (U, indicated by dashed circles), then the number of migrations (M, indicated by bichromatic edges), and finally, the number of comigrations (C). Here, there are two distinct optimal solutions with different patterns of migration, as shown by their migration graphs (rectangular nodes). (d) The corresponding summary graph of all migration graphs.

The *migration criterion*, which minimizes the number of migrations was first proposed by Slatkin and Madisson [22] and later, used by McPherson et al. [23] to reconstruct migration histories of metastatic ovarian cancers. Extending this work, the MACHINA algorithm [19] proposed to solve the Parsimonious Migration History with Tree Refinement (PMH-TR) problem (Fig. 1c), which included two additional criteria: (i) the *comigration criterion*, minimizing the number of comigrations, or simultaneous migrations of one or more clones from one location to another [19, 24–33], and (ii) the *seeding location criterion*, minimizing the number of locations that seed other metastases. The PMH-TR problem also addresses uncertainty reflected by *polytomies*, i. e. nodes with more than two children. While each cell division results in two daughter cells, the input clonal tree might be non-binary and contain polytomies typically due to data limitations such as limited sequencing depth. A subset of polytomies can be refined by introducing additional nodes and edges guided by minimization of the parsimony criteria (Fig. S1). In recent work, Roddur et al. [34] modified the comigration criterion to ensure that comigrations maintain temporal consistency with respect to the output clonal tree.

One key limitation of MACHINA is that it produces a single optimal solution to the PMH-TR problem, overlooking alternative explanations for metastatic spread that are equally supported by the data. Importantly, not taking into account the full solution space, and focusing on a single solution, underestimates the extent of uncertainty and may lead to incorrect conclusions about metastatic spread. A second limitation of MACHINA is that it adheres to a fixed criteria ordering: first minimizing migrations, then comigrations, and finally seeding locations. A recent method, Metient [20] overcomes these two issues by returning multiple solutions minimizing the three discussed criteria in any of six possible permutations. Metient ranks the identified solutions by learning criterion-specific weights across a patient cohort using prior information in the form of the branch lengths/genetic distances of migration edges and, optionally, organotropism information (prior probabilities of directional seeding between a pair of locations). The stochastic gradient descent approach employed by Metient offers no guarantees regarding optimality of the identified solutions nor guarantees about the completeness of the reconstructed solution space. This combined with the requirement for a homogeneous cohort of patients to properly calibrate the criterion-specific weights, potential unreliability in genetic distances, and the seeding location criterion not being realistic for many cancer types, means that Metient, similarly to MACHINA, may overlook alternative parsimonious solutions supported by the data.

Addressing the limitations of prior methods, we introduce MACH2 (Migration Analysis of Clonal Histories 2), a new method that guarantees to enumerate all optimal solutions to the PMH-TR problem. Analogous to the unsampled lineage criterion used in phylodynamics [35–38], MACH2 introduces the *unobserved clone criterion*, minimizing the number of clones assigned to locations in which they were not observed (Fig. S1). Rather than adhering to a specific fixed ordering, MACH2 supports the specification of arbitrary criteria orderings of unobserved clones, migrations, and comigrations. In addition, MACH2 ensures the inferred migrations to be temporally consistent [34]. To efficiently enumerate the complete PMH-TR optimal solution space, MACH2 employs a new compact representation of refinements as edge-weighted multigraphs (Fig. S2). MACH2 represents the enumerated solution space in the form a summary graph, allowing one to identify high-confidence migrations (Fig. 1d). Alongside MACH2, we introduce MACH2-viz, an interactive tool for visualizing MACH2’s solution space. On the theoretical side, we provide new complexity results for the decision and counting versions of the PMH-TR problem. Running MACH2 on simulated data with different criteria orderings, along with MACHINA and Metient, we found minimizing the number unobserved clones first, followed by the number of migrations and comigrations yielded the best results, and made this ordering the default recommendation for MACH2. Moreover, we found MACH2 to identify extensive uncertainty in real data of metastatic non-small cell lung [39], ovarian [23] and prostate [40] cancers, with the input data supporting many alternative migration histories. In many cases, MACH2’s solutions are more consistent with unused, orthogonal data compared to Metient and MACHINA.

## 2 Problem Statement

Our input is a clonal tree inferred by existing methods [12–18] from single-cell and/or bulk DNA sequencing data of matched primary and metastasis samples. Mathematically, a *clonal tree* is a tree *T* rooted at *r*(*T*) with node set *V* (*T*), leaf set *L*(*T*) and edge set *E*(*T*). The nodes *V* (*T*) correspond to clones and each edge (*u, v*) ∈ *E*(*T*) is directed such that *u* is closer to the root *r*(*T*) than *v*. We denote the parent and the set of children of any node *v* by *π*_*T*_ (*v*) and *δ*_*T*_ (*v*), respectively. We write *u* ⪯_*T*_ *v* if node *u* is ancestral to *v*, i. e. there is a directed path from *u* to *v*. Conversely, we write *u*⊥ _*T*_ *v* if *u* and *v* occur on distinct lineages, i. e. neither *u* ⪯_*T*_ *v* nor *v*⪯ _*T*_ *u*. We note that while clonal tree inference methods [12–18] assign mutations to clones, this information will not be used in the problem formulation and the resulting method MACH2. In addition to a clonal tree *T*, we are also given an *observed labeling* 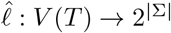that maps each node *u* ∈ *V* (*T*) to a subset 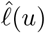 of zero or more locations Σ in which the clone was observed (Fig. 1b).

By definition, a leaf clone does not have any descendants, and for it to be inferred, it must be observed in at least one location. This leads to the following definition of an observed labeling.

### Definition 1.

*An* observed labeling *is a function* 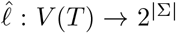 *that maps the clones V* (*T*) *to subsets of locations* Σ *such that each leaf is observed in at least one location, i*. *e*. 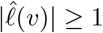 *for all leaves v* ∈ *L*(*T*).

There are three sources of uncertainty regarding a tumor’s migration history. First, if 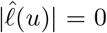 then node *u* was not observed in any location at the time of sequencing. Second, if 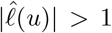 then node *u* was present in multiple locations and the precise migration order among these locations is unknown. Third, tree *T* may include *polytomies*, i. e. nodes with more than two children. To precisely specify the migration history, we need to resolve these sources of uncertainty (Fig. S1). To that end, we seek to refine 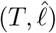 into (*T*^′^, *σ*^*′*^, *ℓ*^*′*^). Tree *T*^′^ is the refined clonal tree obtained from *T* as indicated by the refined-node to original-node mapping *σ*^*′*^ : *V* (*T*^′^) *→ V* (*T*). The function *ℓ*^*′*^ : *V* (*T*^′^) *→* Σ is a location labeling, assigning each node of *T*^′^ exactly one location. We define refinements as follows.

### Definition 2.

*The tuple* (*T*^′^, *σ*^*′*^ : *V* (*T*^′^) *→ V* (*T*), *ℓ*^*′*^ : *V* (*T*^′^) *→* Σ) *forms a* refinement *of* 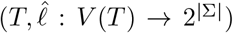 *provided (i) for each original node v* ∈ *V* (*T*), *refined nodes* {*v*^*′*^ ∈ *V* (*T*^′^) | *σ*^*′*^(*v*^*′*^) = *v*} *induce a connected subtree in T*^′^, *(ii) for each original edge* (*u, v*) ∈ *E*(*T*), *there exists exactly one refined edge* (*u*^*′*^, *v*^*′*^) ∈ *E*(*T*^′^) *such that σ*^*′*^(*u*^*′*^) = *u and σ*^*′*^(*v*^*′*^) = *v, (iii) for any two distinct, refined nodes u*^*′*^, *v*^*′*^ ∈ *V* (*T*^′^) *where σ*^*′*^(*u*^*′*^) = *σ*^*′*^(*v*^*′*^), *it holds that ℓ*^*′*^(*u*^*′*^) ≠ *ℓ*^*′*^(*v*^*′*^) *and (iv) for each original node v* ∈*V* (*T*) *observed in location* 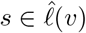, *there exists a refined node v*^*′*^ *in T*^′^ *such that σ*^*′*^(*v*^*′*^) = *v and ℓ*^*′*^(*v*^*′*^) = *s*.

The mapping function *σ*^*′*^ maps each refined node *u*^*′*^ of *T*^′^ to its corresponding original node *σ*(*u*^*′*^) in *T*, with conditions (i) and (ii) preserving original evolutionary relationships in the refined clonal tree *T*^′^, condition (iii) preventing multiple, superfluous edges and migrations of the same original clone to the same location and condition (iv) enforcing the labeling *ℓ*^*′*^ of *T*^′^ to agree with the observed labeling 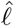of *T* . In contrast with species phylogenetics, where refinements are typically defined for leaf-labeled trees (with leaves corresponding to taxa) our definition of refinement applies to node-labeled trees (where nodes correspond to clones) and includes additional constraints based on the input and output location labelings. In particular, the refined tree *T*^′^ need not have the same set of leaves as the original tree *T* . We refer to Fig. 1c for two example refinements. While there exist many refinements for an input clonal tree *T* and observed labeling 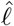, we prioritize output refinements (*T*^′^, *σ*^*′*^, *ℓ*^*′*^) using three different criteria.

First, we count the number *U* (*T*^′^, *σ*^*′*^, *ℓ*^*′*^) of unobserved clones, which are refined nodes that are assigned locations they were not observed in. When a clone is assigned a location where it is not observed, one makes an inference not directly supported by the data. Thus, minimizing this criterion improves data fit (Fig. S1). We note that this criterion is analogous to the unsampled lineage criterion in phylodynamics [35–38].

### Definition 3.

*A refined node v*^*′*^ ∈ *V* (*T*^′^) *is an* unobserved clone *if* 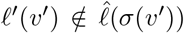, *and* 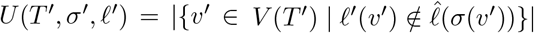 *is the number of unobserved clones*.

Second, we count the number *M* (*T*^′^, *σ*^*′*^, *ℓ*^*′*^) of migrations, defined as follows.

### Definition 4.

*A refined edge* (*u*^*′*^, *v*^*′*^) ∈ *E*(*T*^′^) *is a* migration *if ℓ*^*′*^(*u*^*′*^) ≠ *ℓ*^*′*^(*v*^*′*^), *and M* (*T*^′^, *σ*^*′*^, *ℓ*^*′*^) = |{(*u*^*′*^, *v*^*′*^) ∈ *E*(*T*^′^) | *ℓ*^*′*^(*u*^*′*^) ≠ *ℓ*^*′*^(*v*^*′*^)}| *is the number of migrations*.

Third, we count the number *C*(*T*^′^, *σ*^*′*^, *ℓ*^*′*^) of comigrations, which are defined as follows.

### Definition 5.

*A subset D of migrations is a* comigration *provided ℓ*^*′*^(*u*^*′*^) = *ℓ*^*′*^(*u*^″^) *and ℓ*^*′*^(*v*^*′*^) = *ℓ*^*′*^(*v*^″^) *for any two migrations* (*u*^*′*^, *v*^*′*^), (*u*^″^, *v*^″^) ∈ *D*.

While our clonal trees *T*^′^ lack exact timestamps, its edges *E*(*T*^′^) allow one to temporally order comigrations.

### Definition 6.

*Comigration D*_*p*_ precedes *comigration D*_*q*_, *denoted as D*_*p*_ ≺_comig_ *D*_*q*_, *if there exist migrations* (*u*_*p*_, *v*_*p*_) ∈ *D*_*p*_ *and* (*u*_*q*_, *v*_*q*_) ∈ *D*_*q*_ *inducing a directed path from u*_*p*_ *to v*_*q*_ *in T*^′^, *i*. *e. v*_*p*_ ⪯_*T*_ *′ u*_*q*_.

Following [34], we ensure temporal consistency and define the number *C*(*T*^′^, *σ*^*′*^, *ℓ*^*′*^) of comigrations as follows.

### Definition 7.

*A partition 𝒟 of all migrations of a refinement* (*T*^′^, *σ*^*′*^, *ℓ*^*′*^) *forms* consistent comigrations *provided (i) each part D* ∈ 𝒟 *is a comigration and (ii)* (𝒟, ≺_comig_) *is a partially ordered set*.

### Definition 8.

*The number C*(*T*^′^, *σ*^*′*^, *ℓ*^*′*^) *is the size of the smallest partition of consistent comigrations*.

While the exact computation of *C*(*T*^′^, *σ*^*′*^, *ℓ*^*′*^) is NP-hard, it can be efficiently determined for practical problem instances of moderate sizes using integer linear programming (ILP) [34]. The difference between the number of migrations and comigrations is that the former counts the total number of times an individual clone migrates whereas the latter counts the number of migration events composed of one or more clones. We refer to Fig. 1 and Fig. S1 for examples of scoring refinements using the three criteria discussed above. As discussed previously for migrations and comigrations [41] and illustrated in Fig. S1, there exist tradeoffs between the criteria, i. e. optimizing one criterion often comes at the expense of other criteria. We propose to model the desired tradeoff by taking as input an ordered sub-sequence ℱ = (*f*_1_, …, *f*_*k*_) of the aforementioned three criteria. We say that a solution (*T*^′^, *σ*^*′*^, *ℓ*^*′*^) *lexicographically dominates* solution (*T* ^″^, *σ*^″^, *ℓ*^″^) if there exists a criterion *j*∈ [*k*] for which (*T* ^″^, *σ*^″^, *ℓ*^″^) received a higher cost than (*T*^′^, *σ*^*′*^, *ℓ*^*′*^), i. e. *f*_*j*_(*T* ^″^, *σ*^″^, *ℓ*^″^) *> f*_*j*_(*T*^′^, *σ*^*′*^, *ℓ*^*′*^), while having the same costs in all previous criteria 1≤ *i < j*≤ *k*, i. e. *f*_*i*_(*T*^′^, *σ*^*′*^, *ℓ*^*′*^) = *f*_*i*_(*T* ^″^, *σ*^″^, *ℓ*^″^) for all *i*∈ {1, …, *j*− 1} . Then, the set of *lexicographically optimal refinements* consists of refinements for which no lexicographically dominating refinement exists. In this work, we focus on the following problem.

### Problem 1

(Parsimonious Migration History with Tree Refinement (PMH-TR)). *Given a clonal tree T*, *set* Σ *of locations, observed labeling* 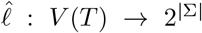 *and criteria ordering* ℱ, *enumerate the space* PMH-TR 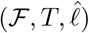 *of all refinements* (*T*^′^, *σ*^*′*^, *ℓ*^*′*^) *of* 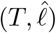 *that are lexicographically optimal with respect to* ℱ.

In this work, we allow the criteria ordering ℱ to correspond to any of the 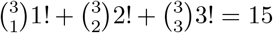 ordered sub-sequences of the three criteria defined above (unobserved clones denoted as ‘U’, migrations as ‘M’, and comigrations as ‘C’). For example, we denote with ℱ = UMC the criteria ordering where first unobserved clones are prioritized, followed by migrations and finally comigrations. While an expert user can specify any of the 15 possible criteria orderings, we set ℱ = UMC as the default recommended criteria ordering based on an extensive simulation study (Section 5. 1). The original MACHINA [19] algorithm returns a single optimal solution for the F ∈ {MC, CM} cases.

## 3 Combinatorial Characterization

Here, we discuss properties of the optimal solution space of PMH-TR. Due to space constraints proofs have been moved to Appendix B. 3. Existing methods, including MACHINA [19] and Metient [20], aim to refine leaf-labeled trees rather than fully-labeled trees. As such, these methods employ a preprocessing step to “pull down” leaves, i. e. attach new leaves to some of the existing nodes of the input clonal tree *T*, assigning each leaf a unique location. This procedure results in a new clonal tree 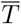 with leaf labeling 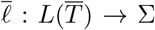 such that there exists a bijection with the pairs (*v, s*) of nodes *v* observed in location 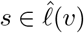 in the original clonal tree *T* . To that end, for each observed node-location pair (*v, s*) (such that *v* is an internal node or a leaf observed in more than one location, i. e. either *v* ∈ *I*(*T*) or *v* ∈ *L*(*T*) and |*ℓ*(*v*)| *>* 1), we add a new leaf *v*^*′*^ and edge (*v, v*^*′*^) labeled by 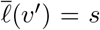. In addition, for each leaf *v* of *T* observed in exactly one location (i. e. 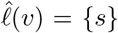), we label *v* with 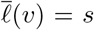 in 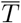. We denote this procedure as PullDownLeaves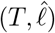, yielding the pair 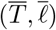. Although only the leaves have observed locations (in fact, in exactly one location), the resulting pair 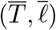 itself is an instance of PMH-TR. We have the following important proposition, showing the redundancy of this preprocessing step.

### Proposition 1.

*Let* 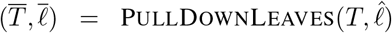 *and* ℱ ∈ {MC, CM}. *Then, the solution space* PMH-TR 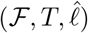 *is a subset of the solution space* FromPulledDownRefinement(PMH-TR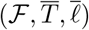).

### Corollary 2.

*Let* 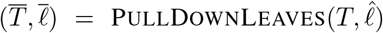 *and* ℱ ∈ {MC, CM}. *Then, all the optimal solutions* (*T*^′^, *σ*^*′*^, *ℓ*^*′*^) *and* 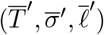 *to* PMH-TR *instances* 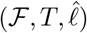 *and* 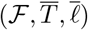, *respectively, have the same number of migrations and comigrations, i*. *e*. 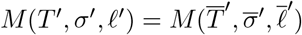 *and* 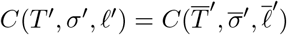.

We prove this by defining an injective function ToPulledDownRefinement (Algorithm S2) to map each optimal refinement (*T*^′^, *σ*^*′*^, *ℓ*^*′*^) of 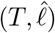 to a refinement 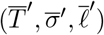 of 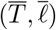, and a surjective function FromPulled-DownRefinement (Algorithm S3) to map each optimal refinement 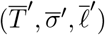 of 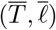 to a refinement (*T*^′^, *σ*^*′*^, *ℓ*^*′*^) of 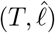. These two functions are defined more precisely in Appendix 3 and are illustrated in Fig. S2. Fig. S11 gives an example of solutions in PMH-TR 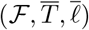 that are not present in PMH-TR 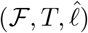. To support efficient exhaustive enumeration of the PMH-TR solution space, we present an equivalent representation of refinements as constrained, directed multigraphs defined as follows.

### Definition 9.

*A directed multigraph R whose nodes V* (*R*) *correspond to locations* Σ *and whose edges are labeled by e* : *E*(*R*) *→ V* (*T*) ∪ *E*(*T*) *is a* refinement graph *for a clonal tree T and observed labeling* 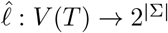 *provided*

i. *for each node u* ∈ *V* (*T*), *the subgraph R*_*u*_ *of R is a directed tree without any multi-edges — where R*_*u*_ *is composed of: (i-a) all nodes s* ∈ *V* (*R*) *with an incoming edge* (*s*^*′*^, *s*) ∈ *E*(*R*) *labeled by either e*((*s*^*′*^, *s*)) = *u or e*((*s*^*′*^, *s*)) = (*π*_*T*_ (*u*), *u*) *or an outgoing edge* (*s, t*) ∈ *E*(*R*) *labeled by either e*((*s, t*)) = *u or e*((*s, t*)) = (*u, v*) *for any v* ∈ *δ*_*T*_ (*u*), *and (i-b) all edges* (*s, t*) ∈ *E*(*R*) *labeled by e*((*s, t*)) = *u;*
ii. *for each edge* (*u, v*) ∈ *E*(*T*), *there is exactly one edge* (*s, t*) ∈ *E*(*R*) *labeled by e*((*s, t*)) = (*u, v*);
iii. *for each node* 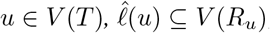;
iv. *for each node u* ∈ *V* (*T*)\{*r*(*T*)}, *R*_*u*_ *is rooted at t if there exists an edge* (*s, t*) *labeled by e*((*s, t*)) = (*π*_*T*_ (*u*), *u*).

Importantly, there exists a bijection between refinement graphs (*R, e*) and refinements (*T*^′^, *σ*^*′*^, *ℓ*^*′*^) (Fig. S2).

### Theorem 3.

*There exists a bijection between refinements* (*T*^′^, *σ*^*′*^, *ℓ*^*′*^) *and pairs* (*R, e*) *of edge-labeled refinement graphs for a given tree T with observed labeling* 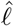.

We prove Theorem 3 by showing that the function ConstructTree 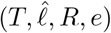 (Algorithm S4) returns a refinement (*T*^′^, *σ*^*′*^, *ℓ*^*′*^) if and only if ConstructGraph 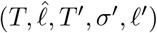 (Algorithm S5) returns the refinement graph (*R, e*). The following corollary directly follows from procedures ConstructTree and ConstructGraph and implies that for each edge (*u, v*) in *T*^′^ there exists exactly one edge (*ℓ*^*′*^(*u*), *ℓ*^*′*^(*v*)) in (*R, e*).

### Corollary 4.

*For any refinement* (*T*^′^, *σ*^*′*^, *ℓ*^*′*^) *of instance* 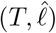, *there is a bijection between the edges in T*^′^ *and the edges in the corresponding refinement graph R*.

We list the properties of the optimal solution space in relation to different criteria ordering in Appendix B. 2. In addition, we prove the decision and counting versions of PMH-TR to be NP-complete and #P-complete for several criteria orderings, including the default ordering ℱ = UMC (Appendix B. 4 and summarized in Fig. S2c).

## 4 Method

In this section, we introduce MACH2 (<u>M</u>igration <u>A</u>nalysis of <u>C</u>lonal <u>H</u>istories 2), an exact algorithm to solve the PMH-TR problem (Section 4. 1) given a clonal tree with observed labeling. To interpret the set of refinements inferred by MACH2, we define a migration graph and summary graph, and introduce MACH2-viz in Section 4. 2.

### 4.1 Integer Linear Program for PMH-TR

Since PMH-TR is both NP-hard and #P-hard (Appendix B. 4), and therefore is unlikely solvable in polynomial time, MACH2 employs an integer linear program (ILP) based on the compact, equivalent representation of solutions with refinement graphs *R* and edge labelings *e* (Section 3). Our ILP is carefully constructed to maintain a bijection between its optimal solutions and the PMH-TR solution space. The ILP has 𝒪 (|Σ| ^2^ |*V* (*T*) |) variables (with the same asymptotic number of integer variables) and 𝒪 (|Σ| ^3^ |*V* (*T*) |) constraints. We provide a complete description of the ILP in Appendix C. MACH2 provides an implementation of the ILP using Gurobi, and is publicly available at https://github.com/elkebir-group/MACH2 under the BSD 3-clause license.

### 4. 2 Interpretation of MACH2 Solution Spaces

To interpret the refinements (*T*^′^, *σ*^*′*^, *ℓ*^*′*^) and analyze the corresponding migration histories in Section 5, we introduce several concepts and a visualization tool. First, following El-Kebir et al. [19], we depict the migration history associated with a refinement (*T*^′^, *σ*^*′*^, *ℓ*^*′*^) by a *migration graph*, defined as follows.

#### Definition 10

(El-Kebir et al. [19]). *A directed multigraph G is a* migration graph *for a PMH-TR solution* (*T*^′^, *σ*^*′*^, *ℓ*^*′*^) *provided (i) the node set V* (*G*) *corresponds to the input locations* Σ *and (ii) for each edge* (*s, t*) *in the multi-set E*(*G*) *of edges there is a unique migration* (*u*^*′*^, *v*^*′*^) *such that *ℓ**^*′*^(*u*^*′*^) = *s and *ℓ**^*′*^(*v*^*′*^) = *t*.

Note that a migration graph *G*^*′*^ for a refinement (*T*^′^, *σ*^*′*^, *ℓ*^*′*^) is obtained by removing self-loops from corresponding refinement graph *R* introduced in Section 3. We show examples of migration graphs in Fig. 1c. To summarize the solution space of PMH-TR instance 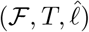. we introduce the summary graph, defined as follows.

#### Definition 11.

*An edge-weighted, directed graph S is a* summary graph *of a solution space PMH-TR*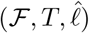*provided (i) the node set V* (*S*) *corresponds to the input locations* Σ *and (ii) S has an edge* (*s, t*) *with weight w*(*s, t*) *>* 0 *if and only if there exist w*(*s, t*) *>* 0 *solutions* (*T*^′^, *σ*^*′*^, *ℓ*^*′*^) ∈ PMH-TR 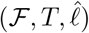 *with a migration* (*u*^*′*^, *v*^*′*^) *from location *ℓ**^*′*^(*u*^*′*^) = *s to location *ℓ**^*′*^(*v*^*′*^) = *t*.

Often we divide the weights by the total number of solutions to get the probability of a migration happening between a pair of locations for PMH-TR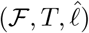, Summary graphs help in quantifying the likelihood of migrations between specific locations and also facilitate focusing on a subset of solutions with specific migrations. We show an example of a summary graph in Fig. 1d. As discussed, both MACHINA and Metient use the seeding location criterion, which minimizes the number of seeding locations defined as follows.

#### Definition 12

(El-Kebir et al. [19]). *A location s* ∈Σ *is a* seeding location *if there is a migration* (*u*^*′*^, *v*^*′*^) *i*. *e. *ℓ**^*′*^(*u*^*′*^) ≠ *ℓ*^*′*^(*v*^*′*^) *such that *ℓ**^*′*^(*u*^*′*^) = *s*.

Although MACH2 does not directly incorporate the seeding location criterion into its problem formulation as it is not realistic for several cancer types, it retains the criterion as an optional filtering step to prioritize solutions if desired. Finally, we present MACH2-viz, a visual tool for exploring PMH-TR solution spaces (Fig. S3). The tool is accessible from MACH2 itself, or through the deployed web application via https://elkebir-group.github.io/mach2-viz/. MACH2-viz is written in JavaScript using React and Cytoscape libraries. The interface allowing users to open the visualizer directly from MACH2 is written in Python, using the Flask library. We refer to Appendix D for more details.

## 5 Results

### 5. 1 Simulations

To benchmark MACH2, we compared it to Metient [20] and MACHINA [19] on 40 previously-published simulation instances [19] with |Σ| ∈ {5, …, 7} locations (referred to as ‘m5’ in [19]). These instances were generated using an agent-based model accounting for tumor evolution, selection and migration. Importantly, the simulation instances contained clones that were not observed in any location (ranging from 5. 88% to 26. 32% of total clones), due to complete clonal expansions as well as limited-resolution sequencing. Each simulation instance consists of a clonal tree *T* with location labeling 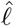, which were provided as input to each method. In addition, we provided Metient with branch lengths corresponding to the number of mutations introduced along each edge. As in the original publication [20], we ran Metient in ‘calibrate’ mode on two cohorts, one cohort containing only primary-to-metastasis spread, and another also containing metastasis-to-metastasis spread. In this simulation study, we assess the number of solutions, whether the enumerated solution space included the ground-truth migration history, and the accuracy of the inferred migration histories and inferred migrating clones. For brevity, we will only report solutions obtained with MACH2 with criteria ordering F = UMC (denoted MACH2-UMC), which outperformed all other criteria orderings (as discussed later).

As an example, we show a simulation instance 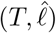 with |*V* (*T*) |= 19 clones and |Σ |= 7 locations in Fig. 2a, and its corresponding ground-truth migration graph *G*^*^ in Fig. 2b. By design, MACHINA inferred only a single solution, which did not match the ground-truth migration graph (Fig. 2c). MACH2-UMC and Metient inferred 8 and 17 solutions, respectively, with the MACH2-UMC summary graph shown in Fig. 2d. While the MACH2-UMC solutions included the ground truth, the Metient solutions did not. To more precisely quantify how well the inferred migration graphs match the ground truth, we use the previously introduced *migration graph F*_1_ *score* [19], which equals the harmonic mean of precision and recall of migration graph edges (see Appendix E). The migration graph *G* reported by MACHINA missed 8 out of 9 migration edges of *G*^*^ and included 7 migration edges not present in *G*^*^, leading to a migration graph *F*_1_ score of 2*/*17 ≈0. 118. On the other hand, the 17 Metient solutions achieved migration graph *F*_1_ scores ranging from 0. 211 to 0. 889 and a median of 0. 211. Finally, the 8 MACH2-UMC solutions achieved migration graph *F*_1_ scores ranging from 0. 778 to 1 and a median of 0. 778. Fig. 2e shows the distributions of the migration graph *F*_1_ scores for this one simulation instance.

**Figure 2:**
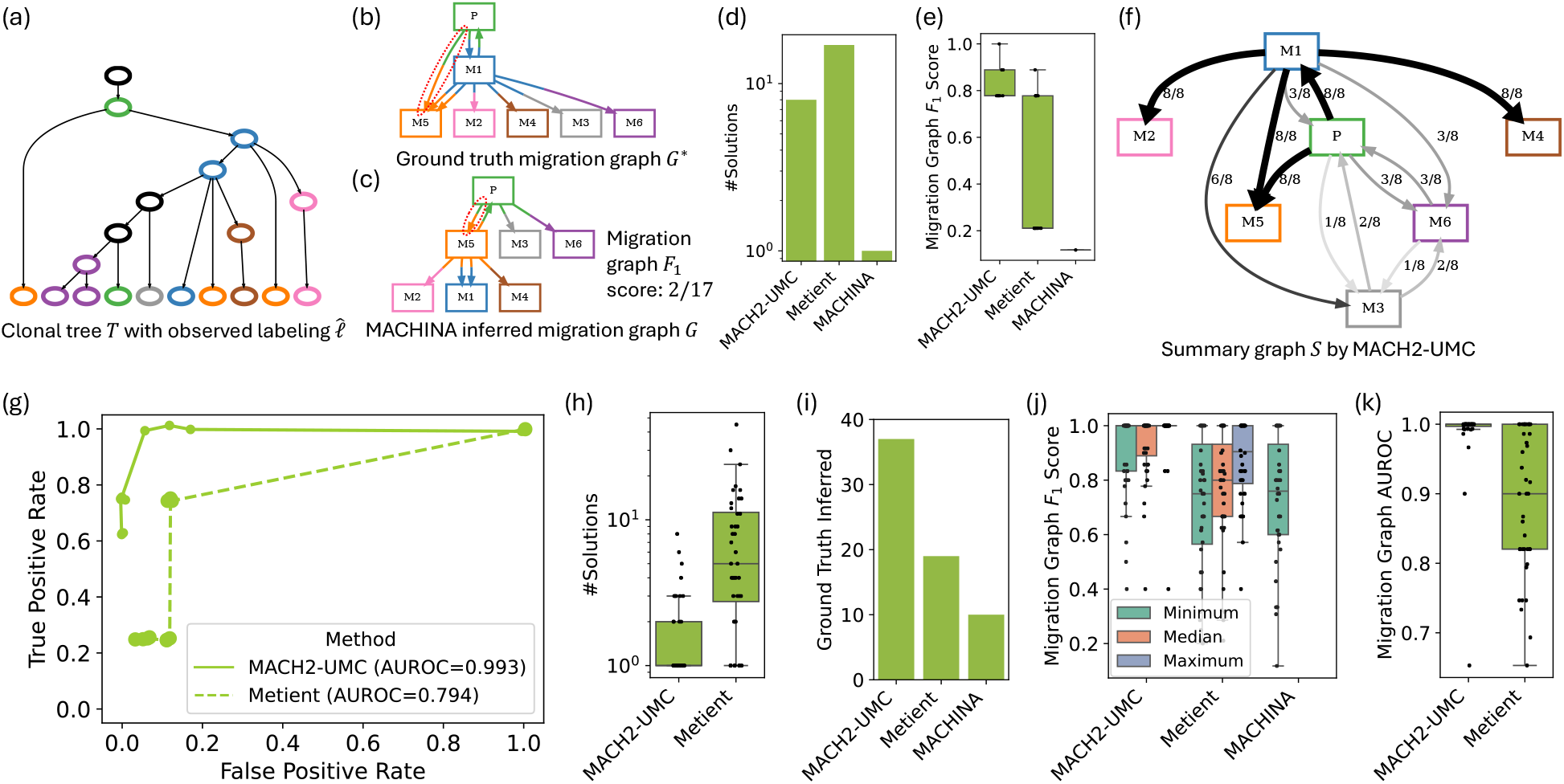
Comparison between MACH2-UMC, Metient [20] and MACHINA [19] on simulated data. (a) Example simulation instance 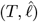 and (b) ground-truth migration graph *G*^*^. (c) The migration graph reported by MACHINA, achieving a migration graph *F*_1_ score of 2*/*17 ≈0. 118 (with dashed ellipses showing matching edges). (d) The number of identified solutions of the illustrated simulation instance. (e) The summary graph of the 8 solutions identified by MACH2-UMC (with bold edges indicating migrations present in all solutions). (f) Migration graph *F*_1_ score of the illustrated simulation instance. (g) ROC curves obtained from summary graphs of the shown simulation instance. (h) MACH2 and Metient inferred multiple solutions for majority of the 40 simulation instances. (i) Number of times each method included the ground-truth solution. (j) Minimum, median, and maximum migration graph *F*_1_ score across all instances (with a single instance shown in Fig. S4b). (k) Distribution of AUROC values across all instances (with a single instance shown in (f)).

To deal with the uncertainty of multiple optimal solutions in real data, one could restrict downstream analyses to only those migrations between locations that have strong support among the enumerated solutions. In other words, for a specified threshold *τ* ∈ [0, 1], one could retain only those summary graph edges that are present in at least a fraction *τ* of the solution space. For the illustrated simulation instance, setting *τ* = 1 restricts analysis to the 5 migrations present in all 8 solutions identified by MACH2-UMC (Fig. 2f). By varying *τ* we can assess the trade-off between sensitivity (or true positive rate) and specificity (or false positive rate) of each method in a Receiver Operating Characteristic (ROC) plot (Fig. 2g). For this instance, we find that MACH2-UMC achieved the largest Area Under the ROC (AUROC) values of 0. 993, outperforming Metient (AUROC: 0. 794).

When considering all 40 simulation instances, all methods, except MACHINA, inferred multiple solutions for the majority of simulation instances (MACH2-UMC ranging from 1 to 8 solutions and Metient from 1 to 45 solutions; Fig. 2h). MACH2-UMC inferred the ground truth 37 out of 40 times (Fig. 2i). On the other hand, Metient included the ground-truth solution only 19 times and MACHINA only 10 times. Furthermore, MACH2-UMC outperformed all other methods for the migration graph *F*_1_ score, achieving median values of 1 for the minimum, median and maximum migration graph *F*_1_ score (Fig. 2j). Metient performed worse than MACH2-UMC, achieving median values of 0. 75 for the minimum, 0. 8 for the median and 0. 89 for the maximum of the computed migration graph *F*_1_ scores across the identified solution spaces. MACHINA achieved median values of 0. 75 for the migration graph *F*_1_ score. Finally, when considering the AUROC values obtained by thresholding the summary graphs of the inferred solution spaces, we find MACH2-UMC performs best, achieving the largest total AUROC value of 39. 46 (summed across all 40 simulation instances) whereas Metient achieved a total AUROC value of 35. 57. We show the AUROC distributions in Fig. 2k.

To assess the impact of the choice of criteria ordering ℱ, we ran MACH2 with 13 different criteria orderings. We excluded MACH2-U and MACH2-C, as they were too unconstrained, leading to too many and often nonsensical solutions and excessive running times. We found that for all criteria ordering, MACH2 inferred multiple solutions for the majority of simulation instances (Fig. S4a). MACH2-UM included the ground-truth solution in its solution space 39 out of 40 times, followed by MACH2-UMC, which returned a subset of MACH2-UM solutions, inferring the ground truth 37 out of 40 times (Fig. S4b). Although both MACH2-UM and MACH2-UMC achieved median values of 1 for the minimum, median and maximum migration graph *F*_1_ score, MACH2-UMC had the highest mean across all the criteria orderings for the minimum (0. 902) and median (0. 929) migration graph *F*_1_ score (Fig. S4c). For AUROC values, both MACH2-UMC and MACH2-UM achieved a median value of 1 (Fig. S4d).

In summary, we found MACH2-UMC performed best with respect to other criteria orderings as well as MACHINA and Metient, demonstrating the importance of the new unobserved clones (U) criterion. In addition, we showed that ignoring alternative optimal solutions might lead to wrong migration history inferences. Finally, we demonstrated the utility of summary graphs to summarize large solution spaces and identify high-confidence migrations.

### 5. 2 Application of MACH2 to Real Data

#### 5. 2. 1 MACH2 Accurately Characterizes Uncertainty in Metastatic Spread

We applied MACH2 in UMC mode, Metient [20], and MACHINA [19] to three real datasets, including (i) one non-small-cell lung cancer patient (CRUK0063) from the TRACERx study [39], (ii) ten androgen-deprived metastatic prostate cancer patients [40], and (iii) seven high-grade serous ovarian cancer patients [23]. Each method was provided the same input clonal tree and observed labeling. In addition, Metient was provided the genetic distance of each edge as input and access to other instances of the same cohort to calibrate its parameters.

For the majority of the real data instances, MACH2-UMC and Metient inferred multiple solutions, with 52 solutions for MACH2-UMC and 5 for Metient in the lung cancer instance, between 1 and 11 solutions for MACH2-UMC and between 1 and 8 for Metient across ten prostate cancer instances, and, between 1 and 44 solutions for MACH2-UMC and between 1 and 8 for Metient in seven ovarian cancer instances (Fig. 3a and Table S1). As shown in Fig. S6a, we found the number of MACH2-UMC inferred solutions to be slightly correlated with an increasing number of input clones (Pearson coefficient *r* = 0. 33; t-test p-value *p* = 0. 01) and locations (*r* = 0. 28; *p* = 0. 04). Since MACH2-UMC prioritizes minimizing the number of unobserved clones, it inferred solutions with the fewest unobserved clones across all 18 real instances compared to Metient and MACHINA (Fig. 3b and Table S1). On average, the median number of unobserved clones for MACH2 was 49. 75% lower than Metient’s and 37. 5% lower than MACHINA’s. Nevertheless, the median number of migrations and comigrations were similar across methods, with the number of migrations in MACH2 being 1. 45% lower than Metient’s and 5% higher than MACHINA’s, while the number of comigrations were 5. 17% and 10. 93% higher in MACH2 than in Metient and in MACHINA, respectively (Fig. 3c-d and Table S1). In terms of running time, MACH2-UMC outperformed both Metient and MACHINA, with a median running time of 0. 44 seconds, compared to 12. 88 seconds for Metient and 2. 09 seconds for MACHINA (Fig. S6b). MACH2 achieves this speed-up due to the compact graph-based representation of refinements as discussed in Section 3. We report the running time and peak memory usage in relation with the number of nodes, locations, and solutions in Fig. S7, showing increases in both quantities with increasing number of nodes, locations, and solutions.

**Figure 3:**
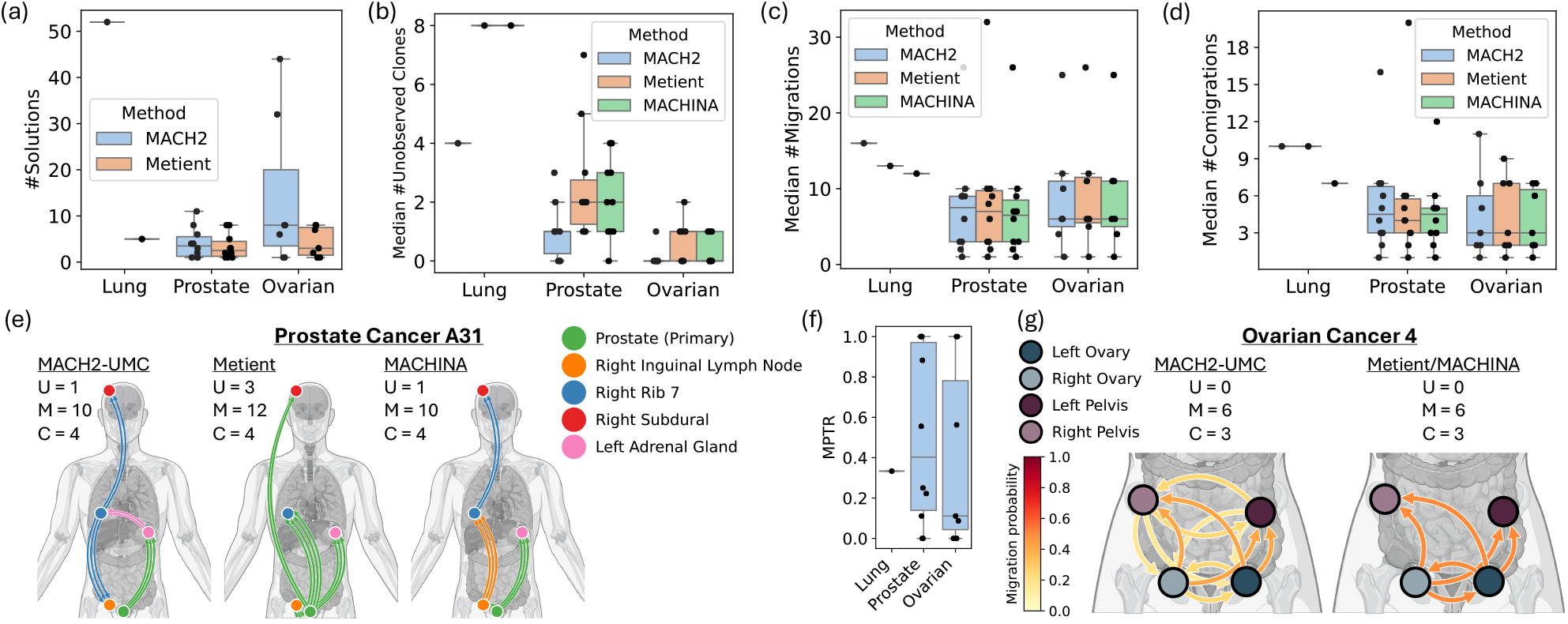
Non-uniqueness of optimal migration histories identified by MACH2. (a) MACH2 and Metient identify multiple solutions for metastatic non-small-cell lung [39], prostate [40] and ovarian [23] cancers. (b) MACH2 in UMC mode minimizes the number unobserved clones, while keeping the number of (c) migrations and (d) comigrations comparable to Metient and MACHINA. (e) For Prostate Cancer A31 [40], both MACH2 and Metient inferred solutions with migration from the primary location (prostate) to the right inguinal lymph node, which is consistent with prior studies [42, 43] unlike the solution returned by MACHINA, where ‘right rib 7’ seeded the right inguinal lymph node. (f) Migration pattern totality ratio. (g) The solution space of Ovarian Cancer 4 inferred by MACH2 had an MPTR of 0, indicating the complete lack of certainty in the pattern of metastatic spread (see Fig. S10). Due to the seeding location objective, Metient solutions inaccurately indicate only migrations from a primary ovary location to a metastatic site are plausible. U: unobserved clones, M: migrations, and C: comigrations.

Importantly, methods that generate multiple solutions are better at capturing alternative migration patterns, which may be overlooked by methods returning a single solution like MACHINA. Fig. 3e shows such an example, where for Prostate Cancer A31 [40] MACHINA yielded a single solution where the ‘right rib 7’ seeded the right inguinal lymph node, which is in close proximity to the prostate. In contrast, both MACH2 and Metient yielded solutions with the prostate seeding the right inguinal lymph node. This finding better aligns with prior studies suggesting that proximal lymph nodes near the prostate gland are often the initial location of metastasis, from which migrating clones further spread to distant sites [42, 43]. While MACH2 is guaranteed to infer temporally consistent comigrations [34], MACHINA and Metient may infer temporally inconsistent solutions (cf. Prostate Cancer A22 for Metient in Fig. S8). Beyond including more biologically-justifiable migration histories, MACH2’s ability to generate the complete solution space allows it to fully characterize the uncertainty inherent to migration history inference. This is a key ability that will allow one to avoid drawing incorrect conclusions due to not fully taking the inherent, data-specific uncertainty in migration history inference into account. In such cases, summary graphs can be used to identify possible migrations along with probabilities. In Fig. S9a, the summary graph for Prostate Cancer A24 contained 2 migrations that were shared across all 8 solutions (prostate to right axillary lymph node, right diaphragm to right rib). Therefore, it can be said with greater certainty that these migrations likely occurred. Conversely, edges in the summary graph with probabilities below 1 indicate uncertainty about the corresponding migration. For example, the probability of xiphoid to be seeded from right axillary lymph node, right rib, or right diaphragm was 4*/*8 = 0. 5, 2*/*8 = 0. 25, and 2*/*8 = 0. 25, respectively. We propose to quantify this uncertainty using the *migration pattern totality ratio* (MPTR), defined as the ratio of the intersection to the union of migration graph edges among all solutions. As such, an MPTR of 0 indicates no shared migrations common to all solutions whereas an MPTR of 1 indicates that each migration is present in all solutions (details in Section E. 3).

In our real data experiments, we found MPTR to decrease with increasing number of solutions (Fig. 3b and Table S1). Strikingly, we found instances where MPTR was 0, indicating the absence of shared migrations across all the solutions (Fig. 3f and Table S1). One such extreme case is Ovarian Cancer 4 [23] where MACH2-UMC predicted 32 different solutions, corresponding to all possible tree-like seeding patterns among the left ovary, right ovary, left pelvic area, and right pelvic area (Fig. S10). The corresponding summary graph in Fig. 3g predicts the probability of migration between any two pairs of locations to be nonzero, but less than one. Note that although Metient, like MACH2, returns multiple solutions, it often tries to reduce the number of seeding locations, i. e. locations from where a clone migrated to another location, and as a result, may miss plausible migration histories in many cases. For Ovarian Cancer 4, both Metient and MACHINA inferred only two solutions, one for considering either ovary as the primary location, where migration only happened from the primary location to the other sites. Thus, Metient and MACHINA overlook the possibility of metastasis-to-metastasis spread, which is frequent in ovarian cancer due to the peritoneal cavity lacking distinct physical barriers. Nevertheless, one may wish to consider the seeding clone criterion for certain cancer types. This is supported by MACH2 as an additional filtering step post inference. In Fig. S9b, for Prostate Cancer A24, prioritizing solutions with the fewest seeding locations yielded two solutions with four shared migrations. While MACH2-UMC assigned a 6*/*8 = 0. 75 probability to the migration from the right axillary lymph node to the right diaphragm, minimizing seeding locations increased this probability to 1. On the other hand, 3 migrations from xiphoid and right rib did not appear in any of the solutions after prioritization.

#### 5. 2. 2 MACH2 Inferred Solutions Are Consistent with Orthogonal Experimental Data

We found that solutions inferred by MACH2 in UMC mode that MACHINA and Metient did not infer better match orthogonal experimental data. We present two such examples here. First, for non-small-cell Lung Cancer CRUK0063 [39], samples were collected at three time points: (i) on day 0, a primary lung sample (P) and a metastatic lymph node sample (LN); (ii) on day 467, after relapse, a paravertebral sample (R); and (iii) on day 857, post-mortem, four lung samples (L1-L4) and one from the paravertebral area (B). We used the clonal tree *T* and observed labeling 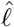 reported in the original paper, which were inferred by CONIPHER [12] and contained |*V* (*T*) |= 46 clones (Fig. 4a). Running MACH2-UMC, which does not consider sample collection time, produced 52 solutions with 4 unobserved clones, 16 migrations, and 10 comigrations (Fig. 4b). Strikingly, for all the 52 solutions, the initial seeding of the relapse sample R (collected at day 467) was from the primary tumor P, involving a migration from Clone 20 (labeled by P) to Clone 10 (labeled by R). All other seeding events involving the samples collected post mortem (on day 857) corresponded to migrations of clones descendant from Clone 10.

**Figure 4:**
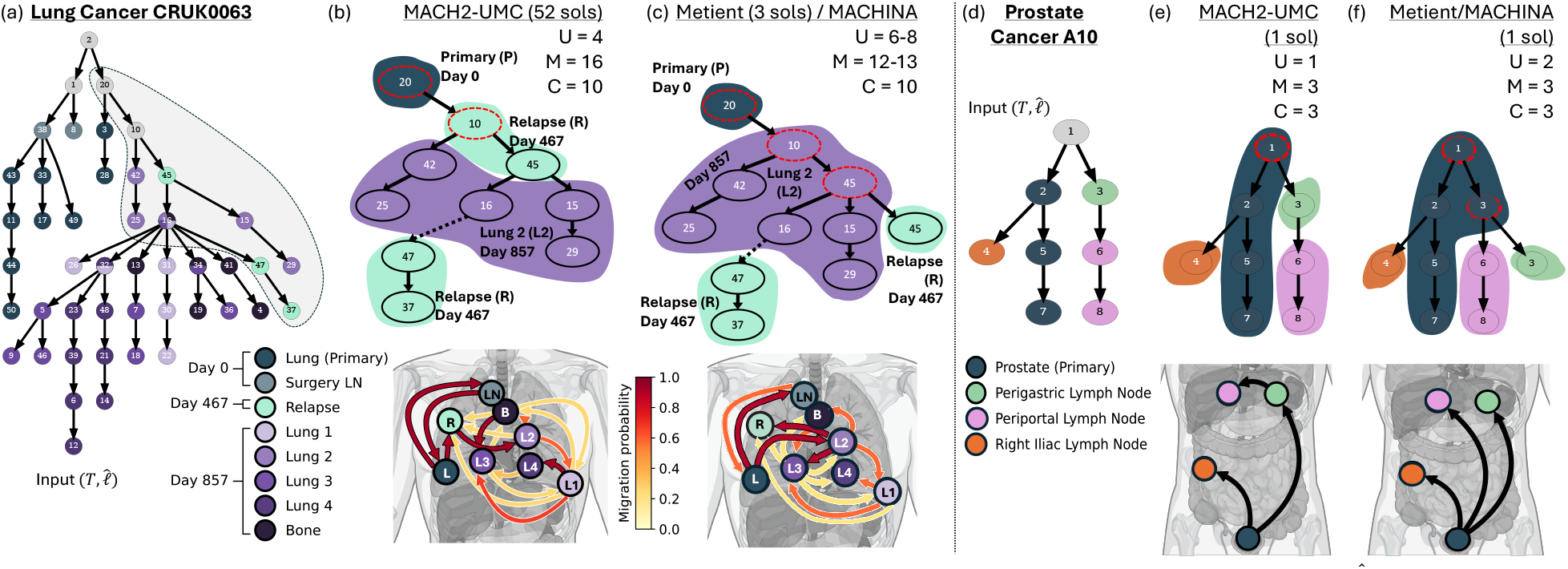
MACH2 solutions match orthogonal experimental data. (a) Clonal tree *T* with observed labeling 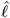 (colors) for Lung Cancer CRUK0063 [39]. (b) Summary graph and subtree present in all 52 MACH2-UMC solutions with colors and contours indicating locations, red dashed nodes indicating unobserved clones, and dashed edge indicating a path in the actual refinements. In all MACH2 solutions, Relapse (R) was seeded initially from the Primary Lung (L), agreeing with sample collection times. (c) Summary graph and subtree present in all 3 Metient solutions. In these solutions, Relapse (R) was seeded from post-mortem sample Lung 2 (L2), disagreeing with sample collection times. Additionally, unlike MACH2 solutions, Clone 45 is unobserved (red dashed) in L2 in the Metient solutions. (d) Clonal tree *T* with observed labeling 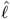 (colors) for Prostate Cancer A10. (e) In the unique MACH2-UMC solution, Clone 3 is observed only in the periportal lymph node, leading to seeding of the periportal lymph node from the perigastric lymph node. (f) Metient and MACHINA additionally assign Clone 3 to the prostate as an unobserved clone (red dashed), resulting in metastasis from the primary tumor only. U: unobserved clones, M: migrations, and C: comigrations.

On the other hand, the three solutions inferred by Metient, which included the MACHINA solution, had more unobserved clones (ranging from 6 to 8) but fewer migrations (ranging from 12 to 13) and the same number of 10 comigrations (Fig. 4c). Importantly, in these solutions, P (collected at day 0) seeded L2 (collected post mortem at day 857), which in turn, seeded R (collected at day 467). This is an unrealistic explanation, as it would imply the existence of undetected tumor in the L2 region at or prior to the collection time of the relapse (R) sample. Closer inspection reveals that the three Metient/MACHINA solutions designate L2 as the origin of Clones 10 and 45, leading to a single migration into L2 at the cost of assigning Clone 45 an unobserved location. On the other hand, as MACH2-UMC prioritizes the unobserved clones criterion, it accepts more migrations into L2 (3 migrations) since it leads to fewer unobserved clones, including Clone 45 which is labeled solely by its single observed location R.

Second, for Prostate Cancer A10 [40] with |*V* (*T*) | = 8 clones sequenced in |Σ| = 4 locations (shown in Fig. 4d with the mapping to original clone labels provided in Table S2), MACH2-UMC inferred a solution space composed of a single refinement with 1 unobserved clone, 3 migrations, and 3 comigrations (Fig. 4e). In contrast, both Metient and MACHINA yielded the exact same refinement with 2 unobserved clones and the same number of migrations and comigrations as the MACH2 refinement (Fig. 4f). MACH2-UMC had one fewer unobserved clone as it assigned Clone 3 to the sole location where it was observed, the perigastric lymph node, suggesting a migration from the perigastric lymph node to the periportal lymph node. Since Metient and MACHINA do not attempt to minimize the number of unobserved clones but instead seek to minimize the number of seeding locations (in addition to migrations and comigrations), these methods suggest that Clone 3 originated in the prostate and then migrated to the periportal lymph node. Consequently, all metastases were seeded from the primary tumor at the expense of an additional unobserved clone. Importantly, MACH2’s location assignment of Clone 3 matched experimental findings [40]. While the input clonal tree was constructed from whole-genome sequencing data (at a coverage of 55×), additional targeted deep resequencing (with a coverage of 471×) failed to detect Clone 3 in the prostate. This combined with the physical proximity of the perigastric and the periportal lymph nodes and the fact that the two periportal lymph node clones (Clones 6 and 8) are descendants of Clone 3 suggests that the MACH2 migration history is more plausible than the history inferred by both Metient and MACHINA.

## 6 Discussion

In this study, we introduced MACH2, a new framework for reconstructing metastatic cancer migration histories. Unlike previous methods, MACH2 additionally minimizes the number of unobserved clones, accepts flexible criteria ordering as input, and returns all optimal solutions instead of a single or limited subset of solutions. Moreover, our definition of tree refinement eliminates the need to add leaves or near-duplicate solutions (Fig. S11). To support efficient solution enumeration, we have proposed a compact representation of refinements and demonstrated a bijection between refinements and their compact forms, alongside proving complexity for various criteria orderings. Experiments on simulated data showed MACH2 in UMC mode as the best performer, while on real data, MACH2-UMC revealed biologically meaningful migration histories. There are several future directions. First, we plan to further experiments to determine under which conditions specific criteria orderings are preferred. Second, we plan to support organotropism data, as used in Metient. Third, we plan to account for mutational signature shifts in metastatic samples [44, 45] to further reduce uncertainty in migration history inference. Fourth, providing Metient with MACH2 solutions, which are guaranteed to be complete and optimal, might improve Metient’s performance. Finally, we plan to apply MACH2 to large cohorts of metastatic cancers to gain deeper insights into the migration trends specific to each primary cancer type as well as study whether repeated migration trajectories can be used to reduce uncertainty.

## A Additional Figures and Tables

**Figure S1:**
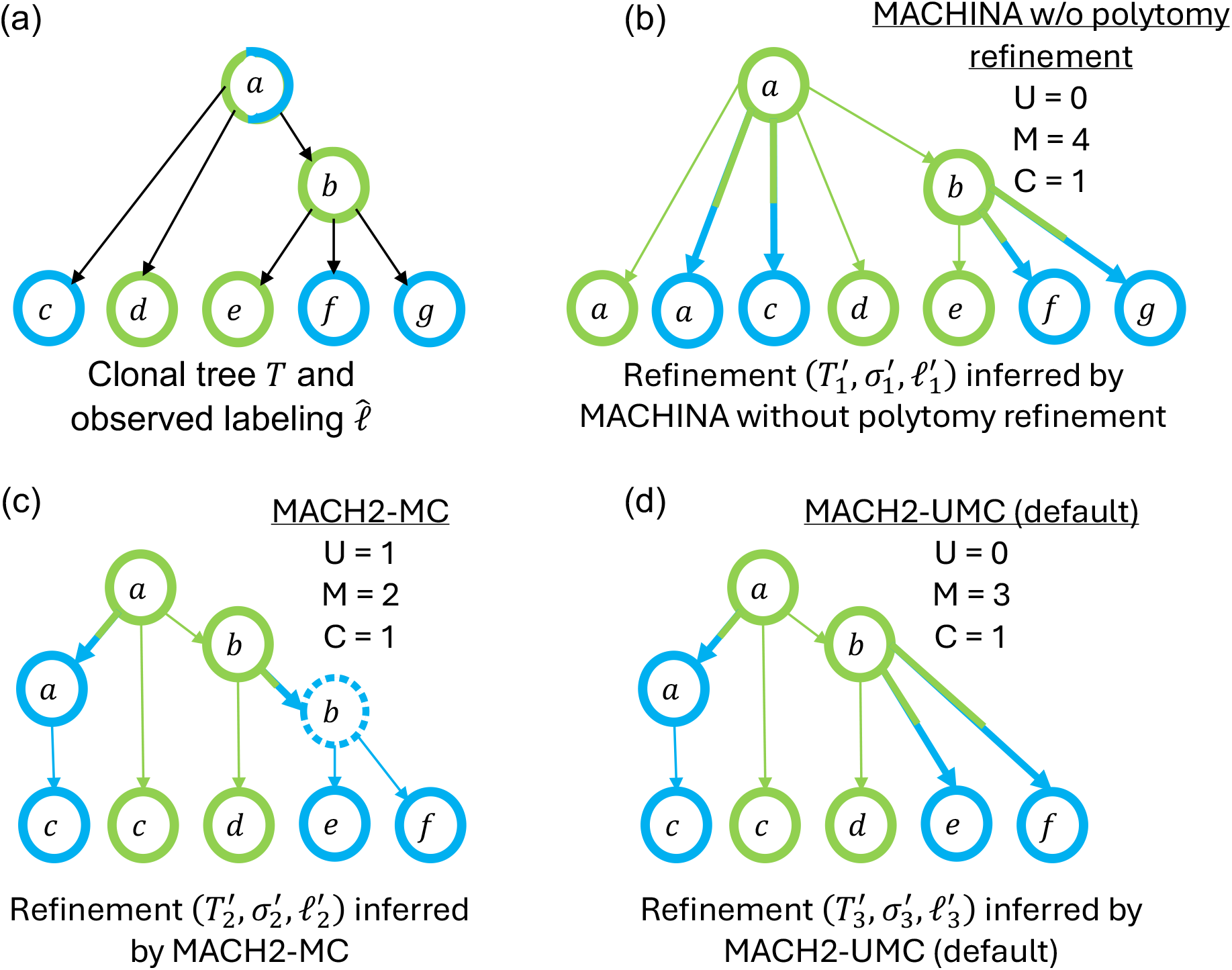
Minimizing the number of unobserved clones led to more realistic scenarios supported by the data. (a) Input clonal tree *T* with observed labeling 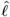 (colors). (b) MACHINA without polytomy refinement returns a solution 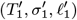 with 4 migrations and without any unobserved clones. Instead of refining node *a*, which is observed in two locations, MACHINA without polytomy refinement pulls down two leaves from *a* and labels them with the locations where *a* is observed. (c) MACH2-MC (also MACHINA with polytomy refinement) aggressively minimizes the number of migrations by refining nodes *a* and *b*, resulting in solution 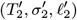 with 2 migrations. Since *b* is present only in the green location, refining *b* produces an unobserved clone (dashed) labeled with the blue location, which is not supported by the input data. (d) MACH2-UMC (default) returns solution 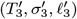, where only the node *a* observed in two locations is refined. Thus, compared to 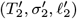, the solution 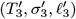 achieves zero unobserved clones at the expense of only one additional migration.

## B Additional Proofs from Combinatorial Characterization and Complexity

### B. 1 Preliminaries

We denote the set of unobserved clones, migrations and comigrations by 𝒰 (*T*^′^, *σ*^*′*^, *ℓ*^*′*^), ℳ (*T*^′^, *σ*^*′*^, *ℓ*^*′*^) and 𝒞 (*T*^′^, *σ*^*′*^, *ℓ*^*′*^), respectively, to distinguish them from their respective cardinalities *U* (*T*^′^, *σ*^*′*^, *ℓ*^*′*^), *M* (*T*^′^, *σ*^*′*^, *ℓ*^*′*^) and *C*(*T*^′^, *σ*^*′*^, *ℓ*^*′*^).

Let 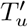 indicate the subgraph of *T*^′^ induced by nodes *u*^*′*^ ∈ *v*(*T*^′^) such that *σ*^*′*^(*u*^*′*^) = *u* (Definition 2). By condition (iii) of Problem 1 it holds that for each refinement (*T*^′^, *ℓ*^*′*^, *σ*^*′*^), two distinct nodes *u*^*′*^ ≠ *u*^″^ of *T*^′^ that correspond to the same node *u* = *σ*^*′*^(*u*^*′*^) = *σ*^*′*^(*u*^″^) of *T* must be labeled by distinct locations, i. e. *ℓ*^*′*^(*u*^*′*^) ≠ *ℓ*^*′*^(*u*^″^). The notation 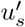 makes use of this fact to refer to the node *u*^*′*^ of *T* such that *σ*^*′*^(*u*^*′*^) = *u* and *ℓ*^*′*^(*u*^*′*^) = *s* (if such node exists). We denote the subtree of *T* rooted at *u* by *T* [*u*].

**Figure S2:**
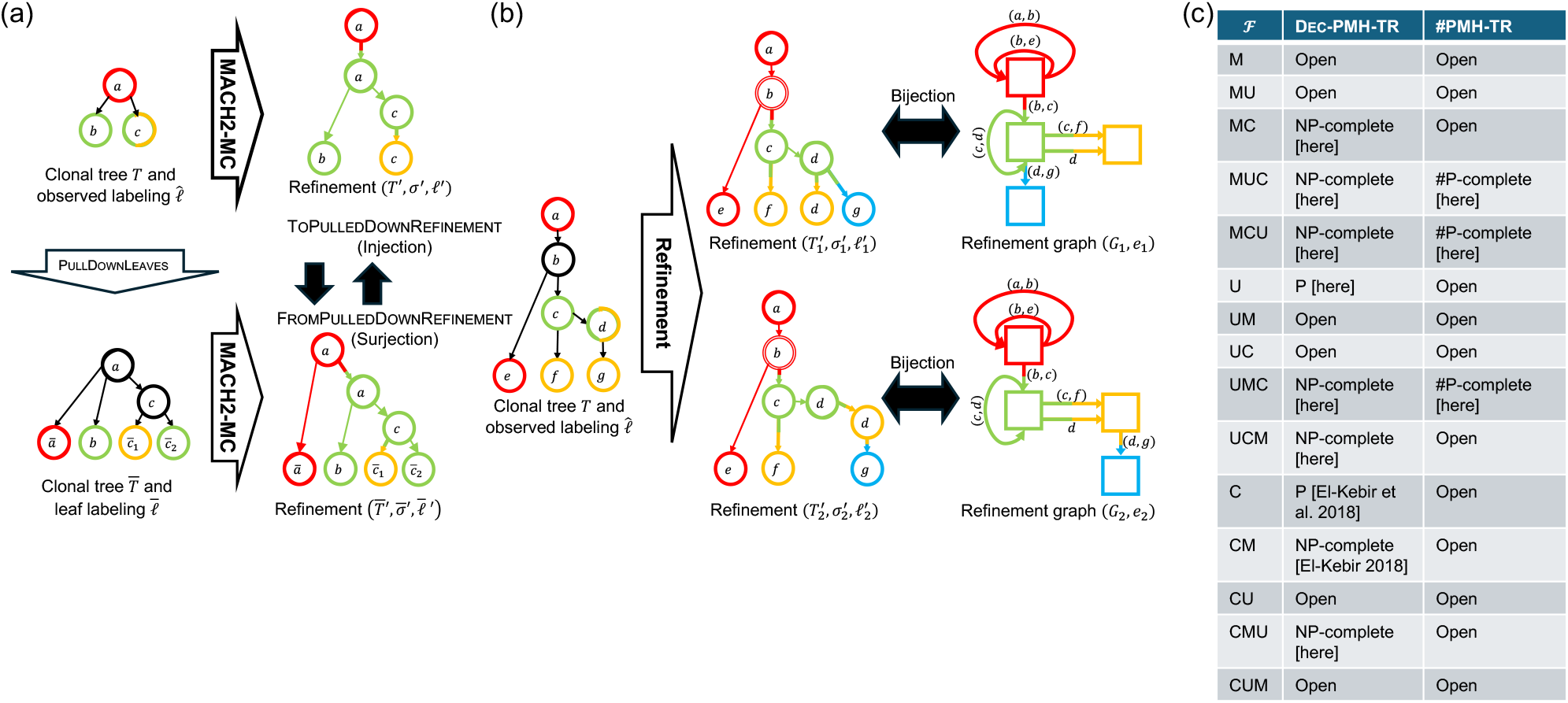
Combinatorial characterization of PMH-TR. (a) Pulling down leaves, as done by MACHINA [19] and Metient [20] maintains optimality of the original solution space when ℱ ∈ *{* MC, CM} . (b) MACH2 relies on a compact and equivalent characterization of solutions as refinement graphs. We prove there exists a bijection between refinements (*T*^′^, *σ*^*′*^, *ℓ*^*′*^) and edge-labeled refinement graphs (*R, e*) of a given tree *T* with observed labeling 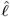. Here, node labels indicate mapping functions 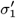 and 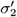, and colored nodes are used to illustrate the observed labeling in 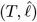, and the location labeling in 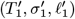 and 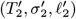. Nodes colored black in 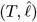 depict that the node was not observed in any location, and double-edged nodes in 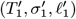 and 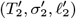 represent unobserved clones. (c) Prior and new complexity results established in this paper.

**Table S1:**
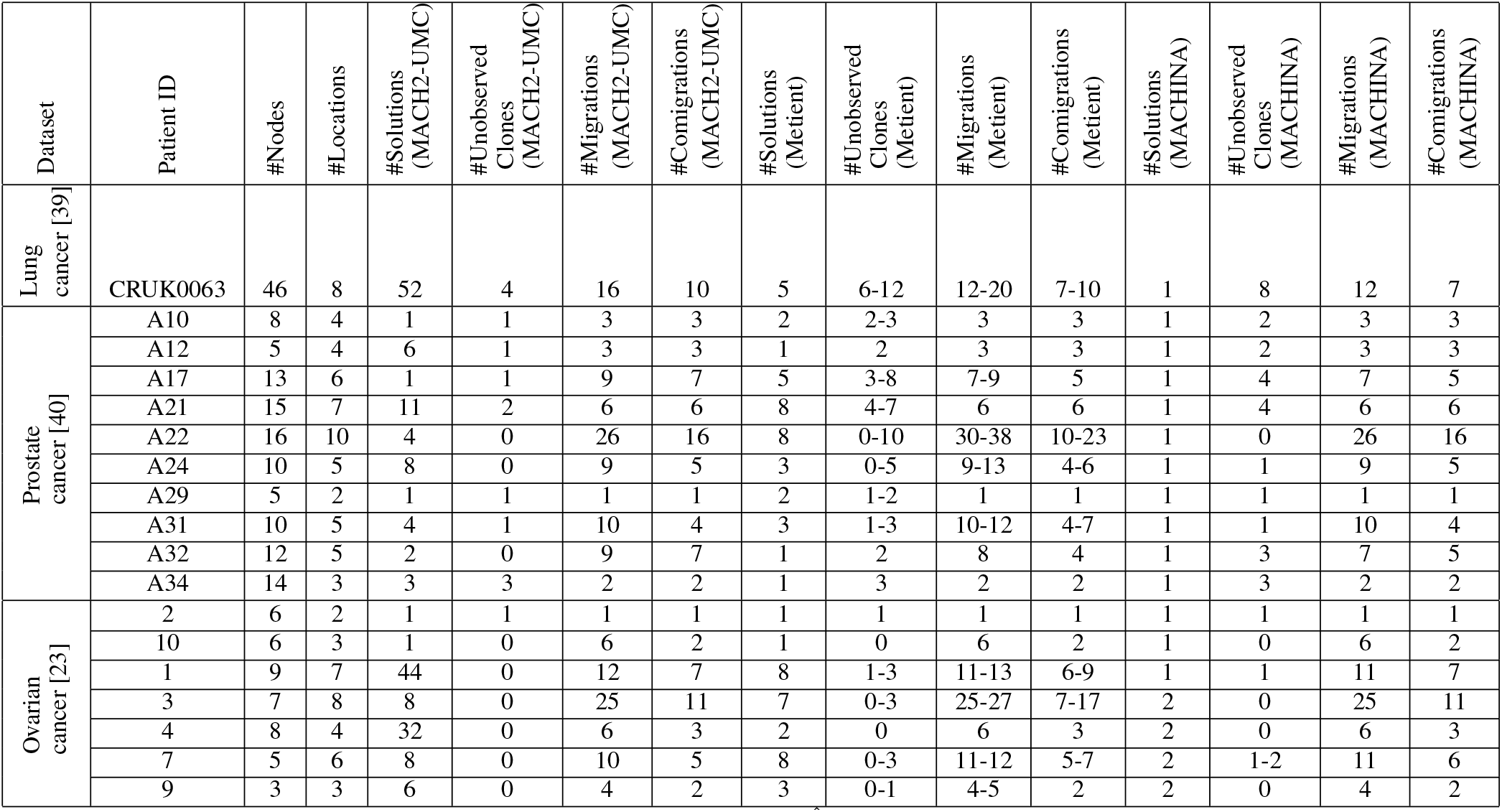
Detailed real data results. For each real data instance 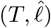, we list the number |*V* (*T*) | of nodes, the number |Σ| of locations, the number of solutions (*T*^′^, *σ*^*′*^, *ℓ*^*′*^), the number U(*T*^′^, *σ*^*′*^, *ℓ*^*′*^) of unobserved clones, the number *M* (*T*^′^, *σ*^*′*^, *ℓ*^*′*^) of migrations and the number *C*(*T*^′^, *σ*^*′*^, *ℓ*^*′*^) of comigrations, identified by MACH2-UMC, Metient [20], and MACHINA [19].

**Figure S3:**
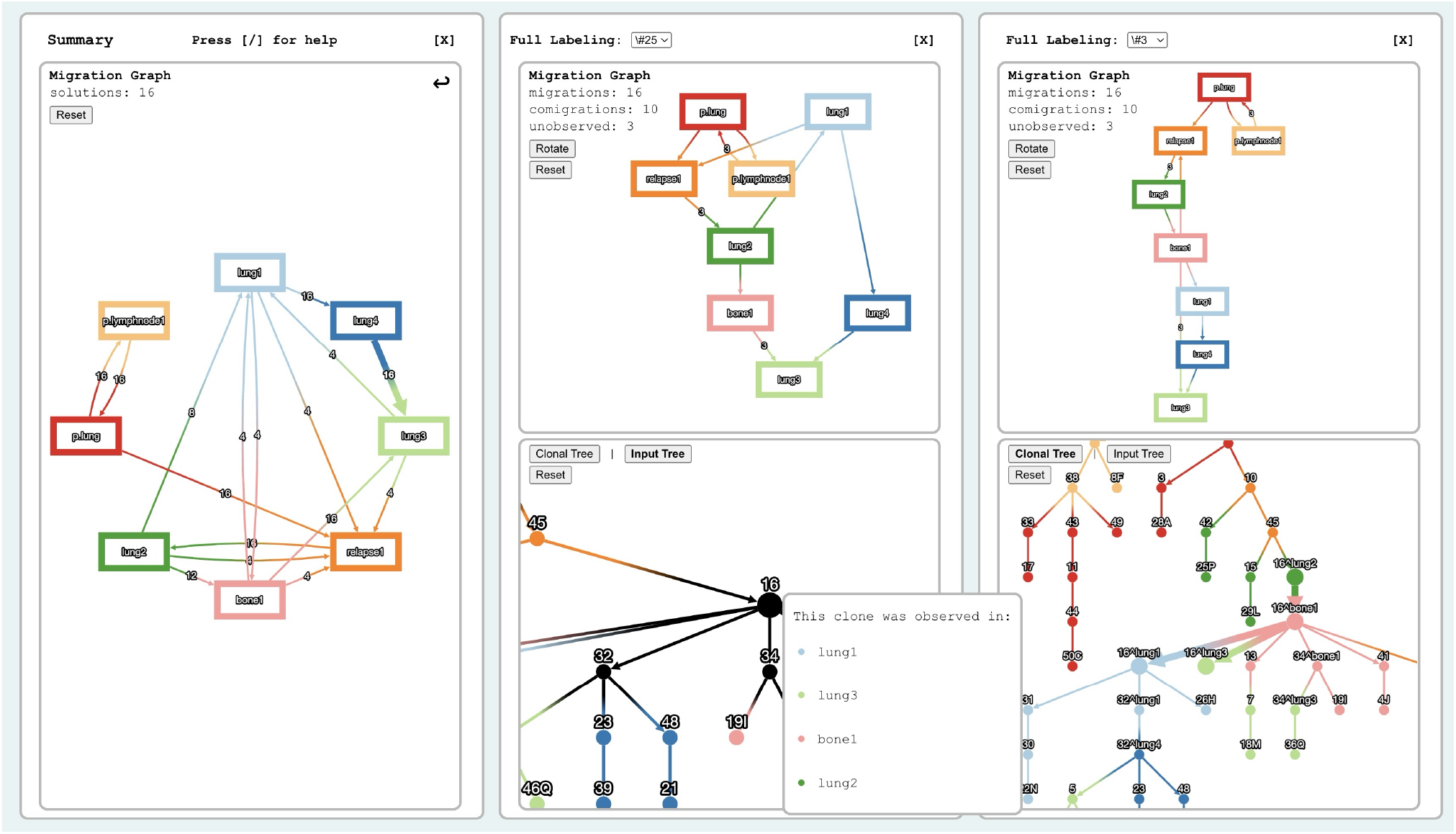
Three panel summary view in `mach2-viz` of the TracerX lung cancer study. The left panel shows the summary graph, computed through the union of edges across the solution space. There is an enforcement of the migration between `lung4` and `lung3`. The right two panels shows a two solutions with a different migration graph each. The middle panel shows an input clonal tree with multiple observed locations for an unlabeled node being hovered over. The corresponding polytomy resolution is highlighted in the rightmost panel’s output clonal tree.

**Table S2:** Mapping of clone names for Prostate Cancer A10.

| Clone name in this manuscript | Clone name in Gundem et al. [40] |
| --- | --- |
| 1 | grey |
| 2 | dark pink |
| 3 | gold |
| 4 | dark brown |
| 5 | pink |
| 6 | dark purple |
| 7 | blue |
| 8 | orange |

### B. 2 Additional Properties of the Solution Space

We have the following lower bound on the number U(*T*^′^, *σ*^*′*^, *ℓ*^*′*^) of unobserved clones.

#### Proposition 5.

*The number* U(*T*^′^, *σ*^*′*^, *ℓ*^*′*^) *of unobserved clones is at least the number* 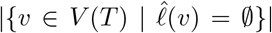 *of clones not observed anywhere*.

The following previously-proven proposition establishes a lower bound on the number *M* (*T*^′^, *σ*^*′*^, *ℓ*^*′*^) of migrations in PMH-TR solution (*T*^′^, *σ*^*′*^, *ℓ*^*′*^).

#### Proposition 6.

(El-Kebir et al. [19]). *For any refinement* (*T*^′^, *σ*^*′*^, *ℓ*^*′*^) *of* 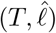, *the number M* (*T*^′^, *σ*^*′*^, *ℓ*^*′*^) *of migrations is at least* |Σ| − 1, *where* 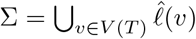 *is the set observed locations*.

Here, we provide the necessary and sufficient conditions to meeting this lower bound.

#### Proposition 7.

*It holds that* 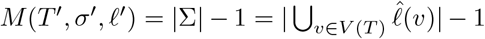 *if and only if (i) *ℓ**^*′*^(*v*^*′*^) ∈ Σ *for each node v*^*′*^ *of T*^′^ *and (ii) for any two nodes w*^*′*^, *w*^″^ *of T*^′^ *where *ℓ**^*′*^(*w*^*′*^) = *ℓ*^*′*^(*w*^″^), *each node in the undirected path from w*^*′*^ *to w*^″^ *is labeled with *ℓ**^*′*^(*w*^*′*^) = *ℓ*^*′*^(*w*^″^).

**Figure S4:**
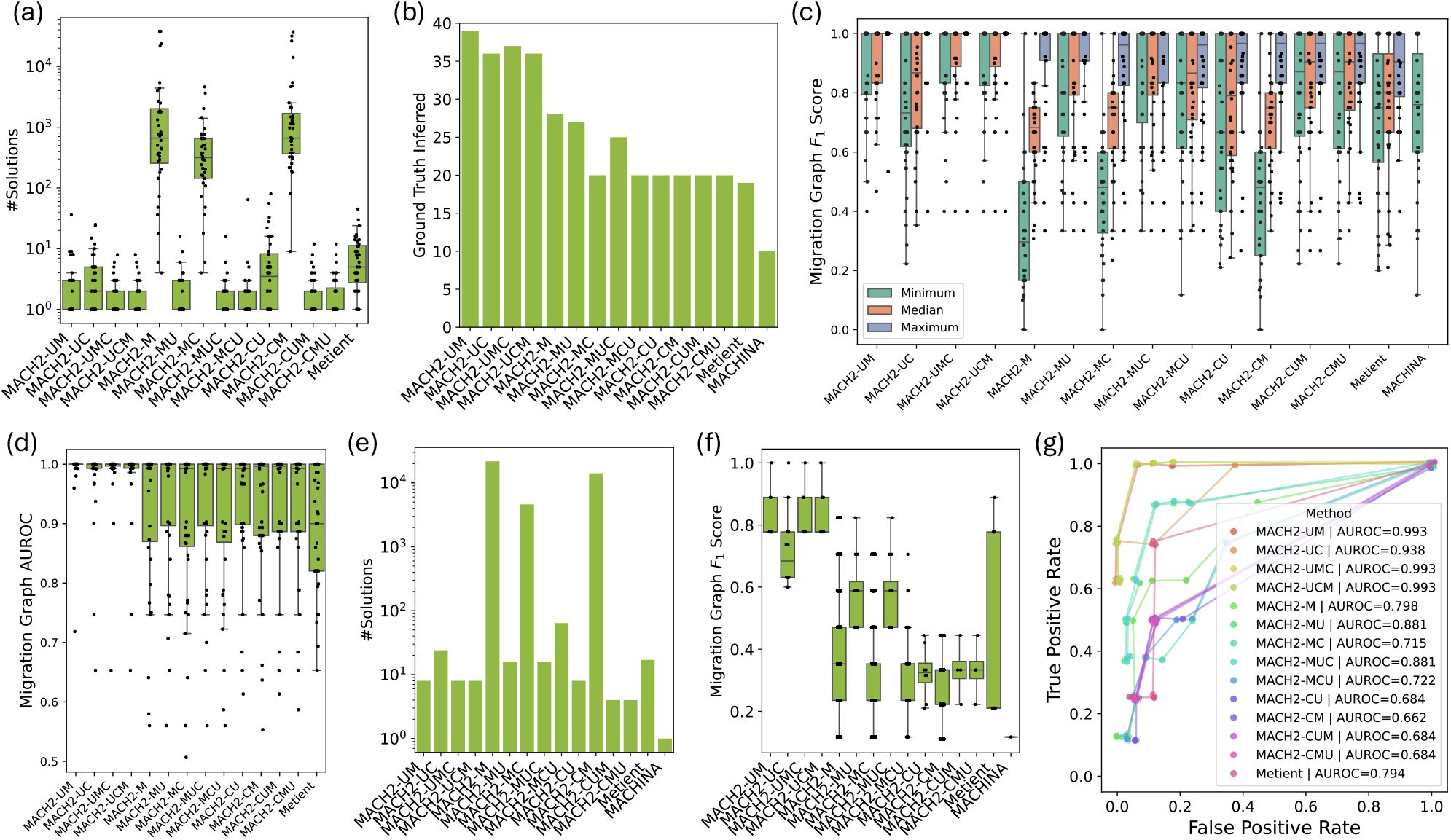
Comparison between MACH2 with 13 criteria orderings, Metient [20] and MACHINA [19] on simulated data. (a) MACH2 with 13 criteria orderings and Metient inferred multiple solutions for the majority of the instances. (b) Number of times each method included the ground-truth solution. (c) Minimum, median, and maximum migration graph *F*_1_ score across all instances. (d) Distribution of AUROC values across all instances. (e) The number of identified solutions of the solution spaces inferred by different methods for the instance discussed in Section 5. 1 and shown in Fig. 2a. (f) ROC curves obtained from summary graphs of the discussed simulation instance. (g) Migration graph *F*_1_ score of the discussed simulation instance.

The next proposition gives an upper bound for the number *C*(*T*^′^, *σ*^*′*^, *ℓ*^*′*^) of comigrations.

#### Proposition 8.

*It holds that C*(*T*^′^, *σ*^*′*^, *ℓ*^*′*^) ≤ |*E*(*R*)| ≤ |*E*(*T*)| + |*V* (*T*)|(|Σ| − 1).

Finally, we have the following propositions that apply to specific criteria orderings ℱ, bounding the number 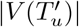 of refined nodes *u*^*′*^ ∈ *V* (*T*^′^) that map to original node *u*, i. e. *σ*^*′*^(*u*^*′*^) = *u*.

#### Proposition 9.

*Let* (*T*^′^, *σ*^*′*^, *ℓ*^*′*^) *be a refinement that minimizes the number of unobserved clones first. Then, for any node u* ∈ *V* (*T*) *observed in less than two locations, i*. *e*. 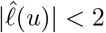, *it holds that* 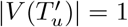.

#### Proposition 10.

*Let* (*T*^′^, *σ*^*′*^, *ℓ*^*′*^) *be a refinement that minimizes the number of migrations first. Then, there exists an optimal refinement* (*T*^′^, *σ*^*′*^, *ℓ*^*′*^) *such that for any node u* ∈ *V* (*T*) *without polytomies and observed in less than two locations, i*. *e*. |*δ*_*T*_ (*u*)| *<* 2 *and* 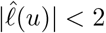, *it holds that* 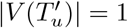.

### B. 3 Combinatorial Characterization

For completeness, we give the pseudocode for the procedure PullDownLeaves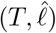 in Algorithm S1.

In this section, we prove the following proposition.

**(Main text) Proposition 1**. *Let* 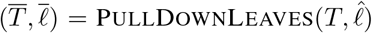 *and* ℱ ∈ {MC, CM}. *Then, the solution space* PMH-TR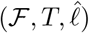 *is a subset of the solution space* FromPulledDownRefinement(PMH-TR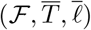).

Let 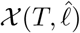 be the set composed of all observed node-location pairs (*v, s*) where *v* ∈ *V* (*T*) and 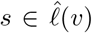. By definition, PullDownLeaves induces a bijection 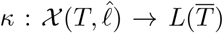 between the observed node-location pairs 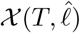 of 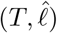 and the location-labeled leaves 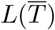 of 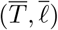. More specifically, 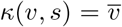 if 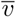 is a pulled-down leaf, i. e. 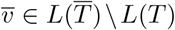 with incoming edge 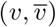 in 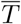 and label 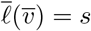;and *κ*(*v, s*) = *v* if *v* is not a pulled-down leaf of *T* observed in unique location *s*, i. e. *v* ∈ *L*(*T*) and 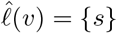. We refer to Fig. S2a for an example.

**Figure S5:**
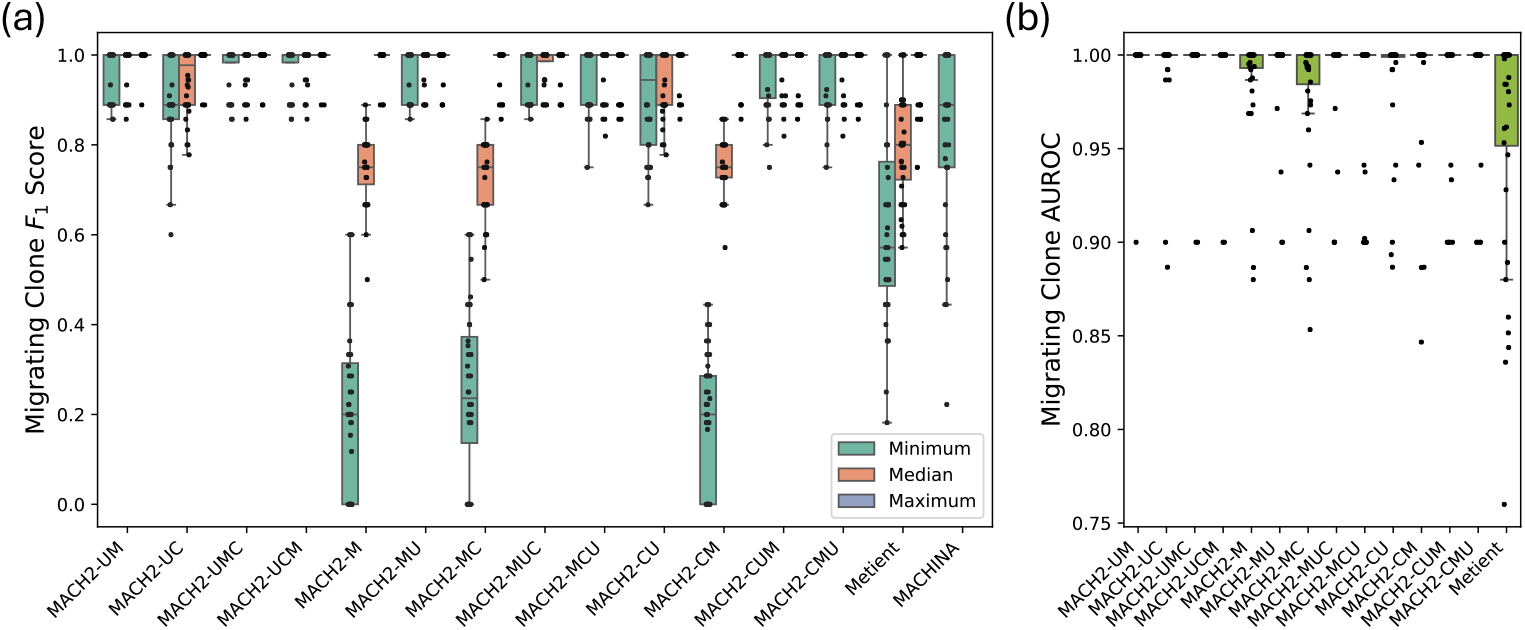
(a) Minimum, median, and maximum migrating clone *F*_1_ score (Section E), and (b) migrating clone area under receiver operating characteristics curve (AUROC) for MACH2 with 13 criteria orderings, Metient, and MACHINA.

**Figure S6:**
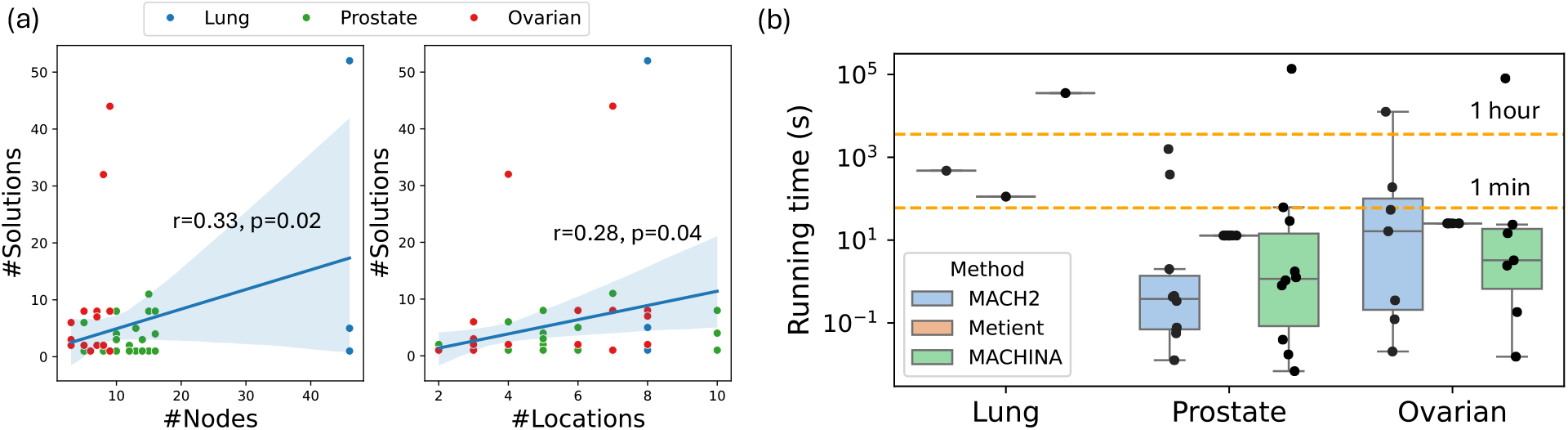
(a) The number of solutions for MACH2-UMC increases with the number of nodes and the number of locations. (b) MACH2 in UMC mode is slightly faster than MACHINA and MACH2. Note that Metient in calibrate mode processes an entire cohort at once. Thus we reported the mean running time per instance, calculated by dividing the total cohort time by the number of instances. All experiments were run on a server with Intel Xeon Gold 5120 dual CPUs with 14 cores each at 2. 20 GHz and 512 GB RAM.

**Table S3:** Mapping of clone names for Prostate Cancer A22.

| Clone name in this manuscript | Clone name in Gundem et al. [40] |
| --- | --- |
| 1 | grey |
| 2 | red |
| 3 | dark brown |
| 4 | mid blue |
| 5 | light green |
| 6 | dark purple |
| 7 | dark blue |
| 8 | orange |
| 9 | mid green |
| 10 | dark green |
| 11 | pink |
| 12 | light purple |
| 13 | gold |
| 14 | light blue |
| 15 | yellow |
| 16 | light brown |

**Figure S7:**
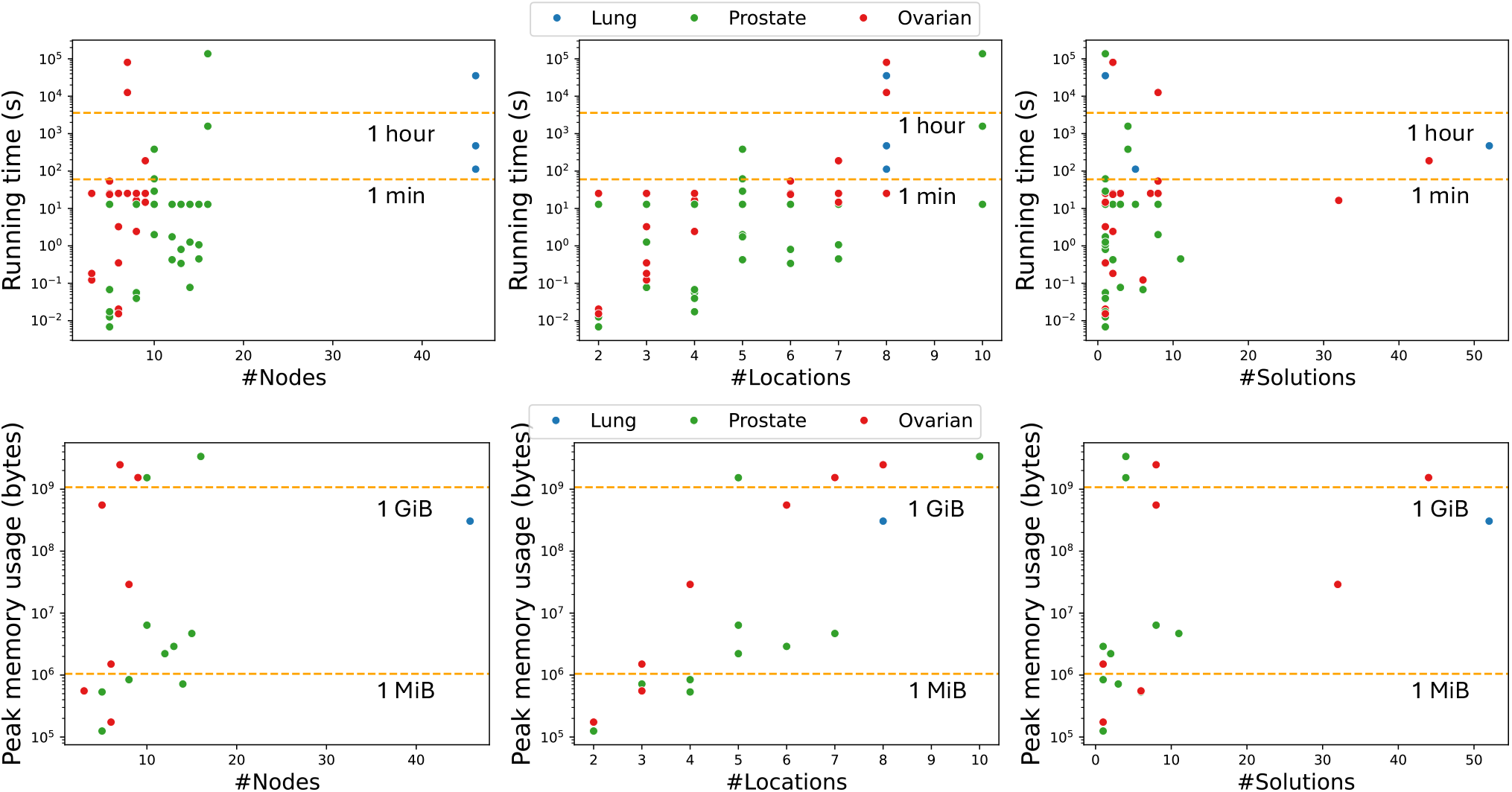
Running time (in seconds) and peak memory usage (in bytes) of MACH2-UUMC as functions of the number of nodes, locations, and solutions. All experiments were run on a server with Intel Xeon Gold 5120 dual CPUs with 14 cores each at 2. 20 GHz and 512 GB RAM.

#### Algorithm S1

Construction of 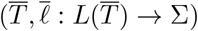 by pulling down leaves from 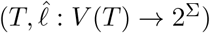

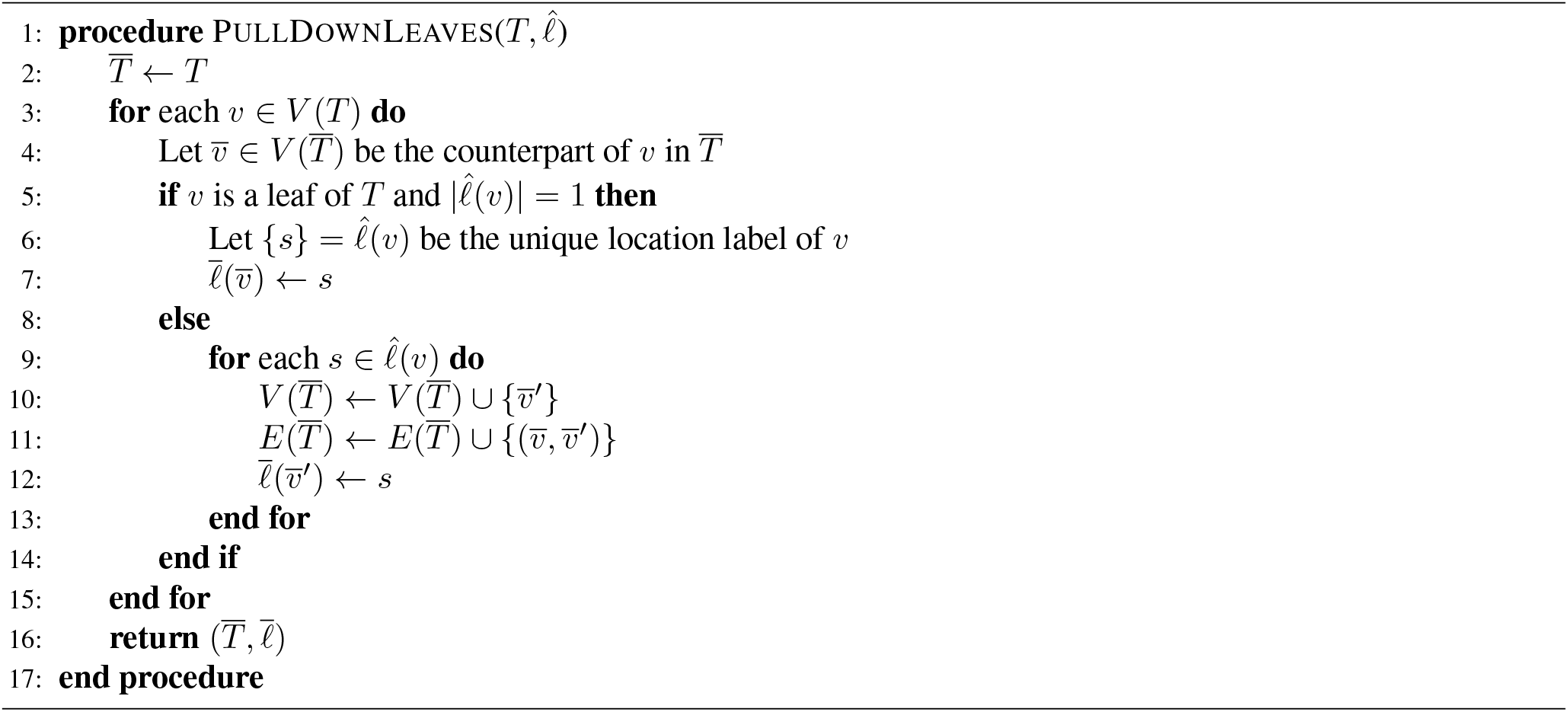

**Figure S8:**
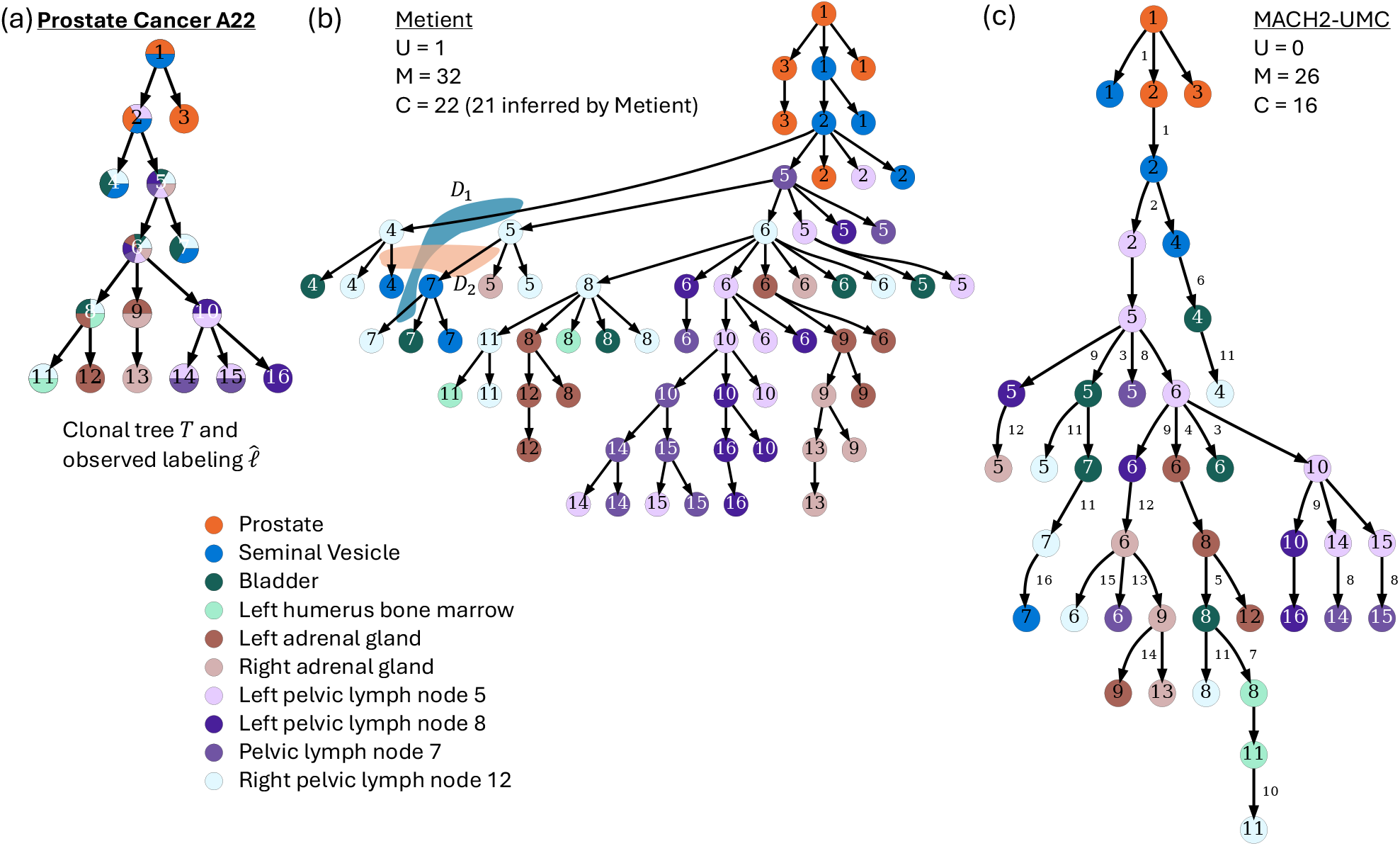
MACH2 accurately infers the number of temporally consistent comigrations. (a) Clonal tree *T* with observed labeling 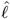 for Prostate Cancer A22. (b) Metient returns a solution with 1 unobserved clone (U), 32 migrations (M) and 22 comigrations (C). But Metient reports the tree to have 21 comigrations. Metient groups the migrations indicated by the contour *D*_1_ in one comigration and the migrations indicated by the contour *D*_2_ in another comigration. However, this creates temporal inconsistency because none of the migrations precede one another. In other words, if migrations from the comigration *D*_1_ happen simultaneously, then migration from the comigration *D*_2_ cannot happen simultaneously, and vice versa. (c) MACH2 returns a solution with 0 unobserved clone (U), 26 migrations (M) and 16 temporally consistent comigrations (C), where migrations with the same label belong to the same comigration.

**Figure S9:**
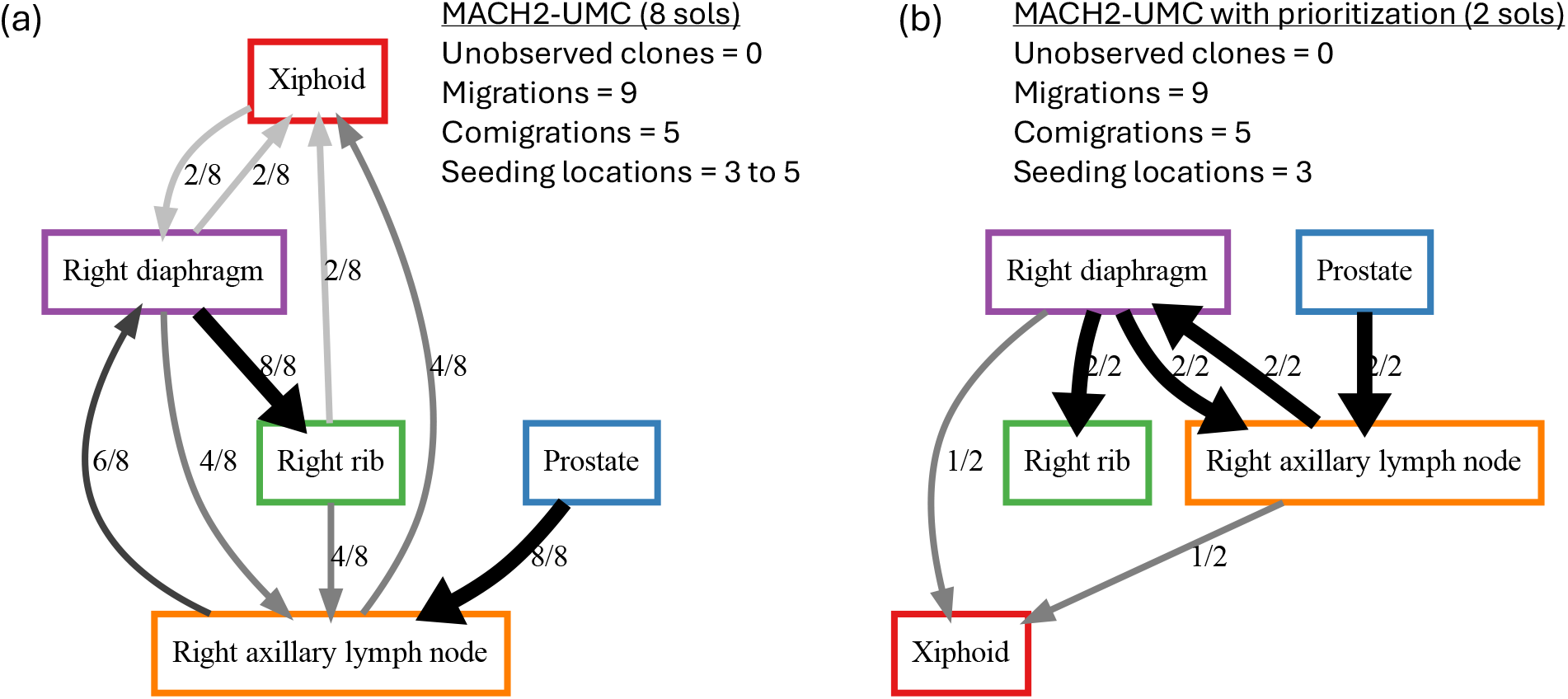
(a) Summary graph for Prostate Cancer A24 with 2 migrations with a probability of 1 (boldface). (b) Summary graph for the same patient after prioritizing solutions that minimizes the number of seeding locations, with 4 migrations with a probability of 1 (boldface).

**Figure S10:**
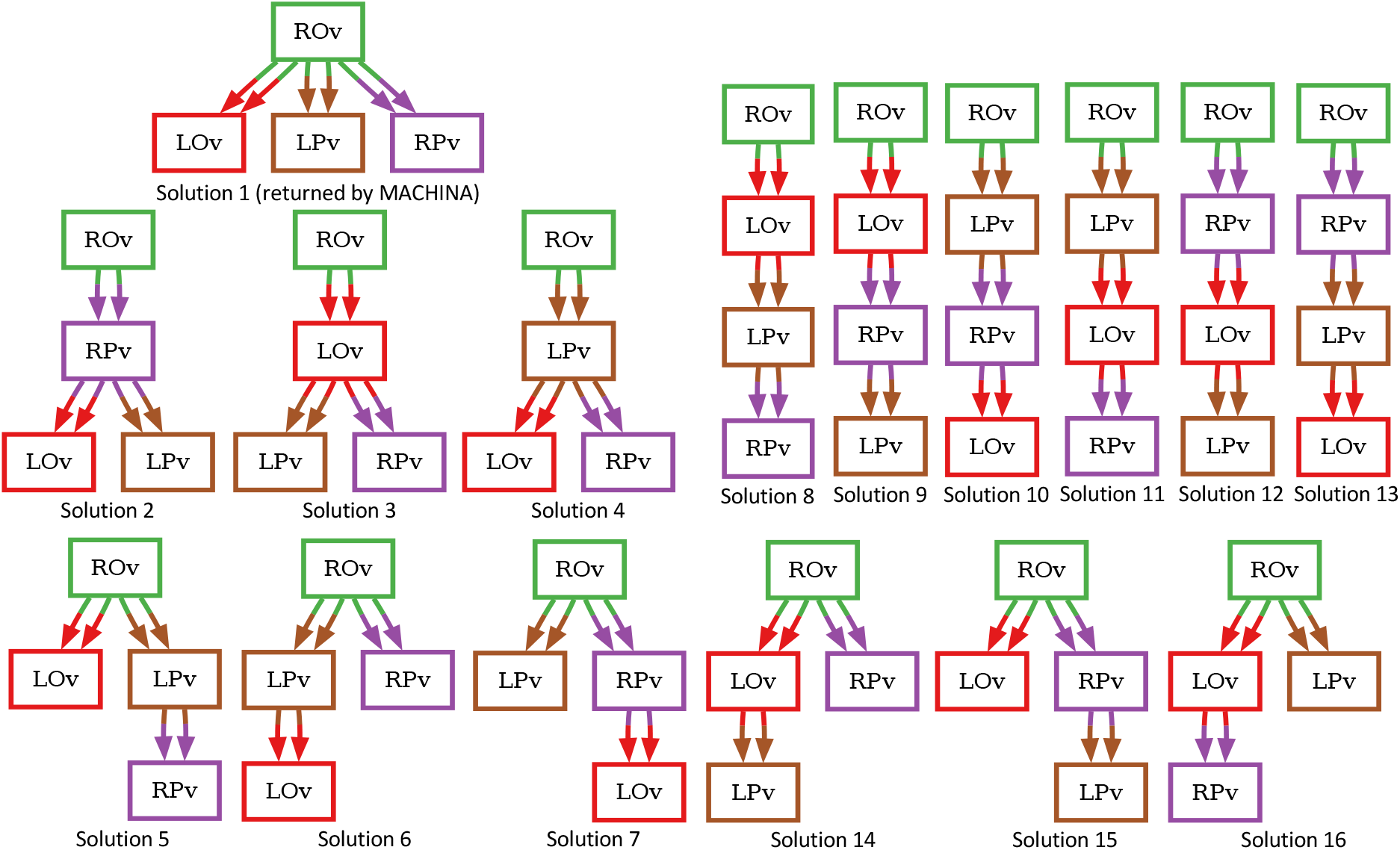
16 solutions inferred by MACH2-UMC for Ovarian Cancer 4 [23] when the right ovary (ROv) is considered as the primary location. The migration graph of each solution is presented, where migration graph is the refinement graph without self-loops. Ignoring multi-edges, each migration graph admits a unique tree topology with the root labeled by the right ovary (ROv) and each of the non-roots labeled either by the left ovary (LOv), the left pelvic area (LPv), or the right pelvic area (RPv). There other 16 MACH2-UMC solutions when the left ovary is considered as the primary location can be obtained by flipping the labels of the left ovary (LOv) and the right ovary for each solution.

**Figure S11:**
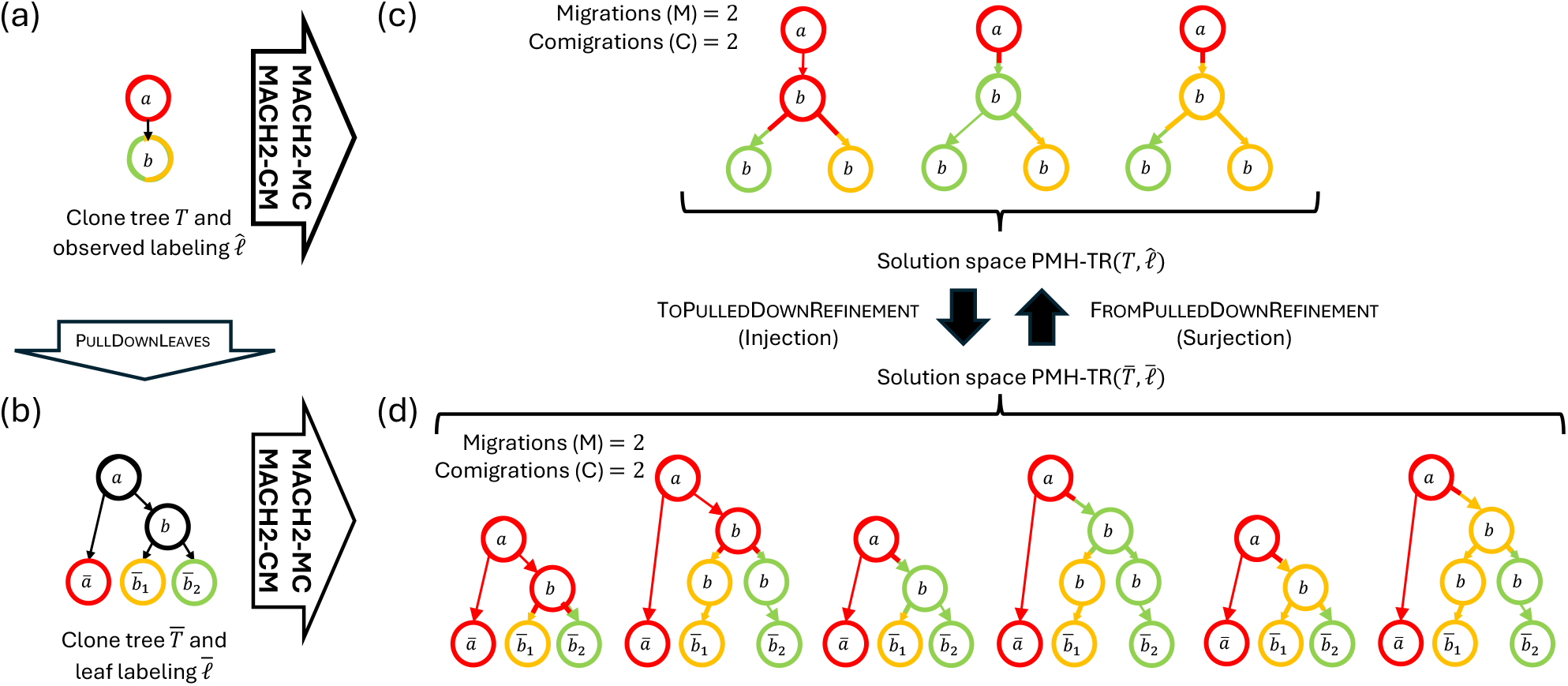
Example showing that PMH-TR 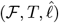 may be a proper subset of FromPulledDownRefinement(PMH-TR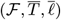). (a) A PMH-TR input instance 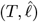 with two clones observed in three locations. (b) The corresponding input instance 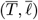 obtained by pulling down leaves from 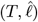. (c) The solution space PMH-TR 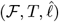 has three solutions with M = 2 migrations and C = 2 comigrations. (d) The solution space PMH-TR 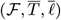 has three additional solutions with M = 2 migrations and C = 2 comigrations.

To prove Proposition 1, we first show that for any optimal solution (*T*^′^, *σ*^*′*^, *ℓ*^*′*^) to PMH-TR instance 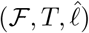, there exists a unique feasible solution (*T ′, σ*^*′*^, **ℓ*′*) to PMH-TR instance 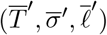 with the same number of migrations and comigrations. To that end, we present a function ToPulledDownRefinement to build a solution 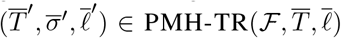 from 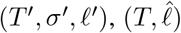 and 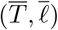, with the same number of migrations and comigrations, i. e. 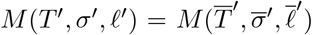 and 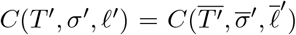. ToPulledDownRefinement does this by iteratively considering all observed node-location pairs 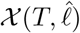 and modifying (*T*^′^, *σ*^*′*^, *ℓ*^*′*^). For each observed node-location pair (*v, s*) 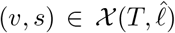, there is exactly one refined node *v*^*′*^ of *T*^′^ labeled by location *ℓ*^*′*^(*v*^*′*^) = *s* that corresponds to the original node *σ*(*v*^*′*^) = *v* — this follows from condition (iv) of Definition 2. Moreover, 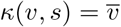 is the pulled down leaf from *v* labeled with 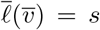 or node *v* itself if *v* is observed in one location. If *v*^*′*^ is an internal node of *T*^′^, we (i) add a new node 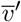, (ii) add the edge 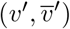, (iii) set the original node of new leaf 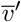 to be 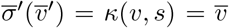, and (iv) label with 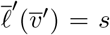. On the other hand, if *v*^*′*^ is a leaf, we update the original node of *v*^*′*^ to be 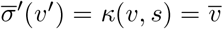. For completeness, we give the pseudocode of the procedure ToPulledDownRefinement in Algorithm S2.

We denote the space of *optimal solutions* of the original PMH-TR instance by PMH-TR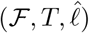. Let 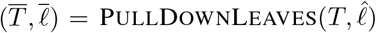 be the pulled down PMH-TR instance obtained from 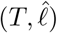. We denote the space of *feasible solutions* of the pulled-down instance by PMH-TR^*^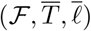In our first lemma, we claim that for ℱ ∈ {MC, MC} the function ToPulledDownRefinement with domain PMH-TR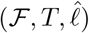 is an injective function for the range of feasible solutions PMH-TR^*^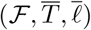with the same number of migrations and comigrations as solutions in PMH-TR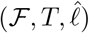.

#### Lemma 11.

*For* ℱ ∈ {MC, MC}, *the function* ToPulledDownRefinement *with domain* PMH-TR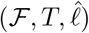 *is an injective function for the range of feasible solutions* PMH-TR^*^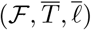*with the same number of migrations and comigrations as solutions in* PMH-TR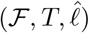.

*Proof*. For any refinement 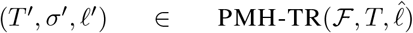, the function call 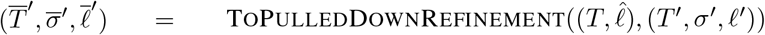 attaches edges to the existing nodes of *T*^′^, sets the original node and the location labels of the new nodes, and ensures 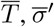 and 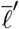 are a valid tree, an original node mapping and a location labeling, respectively. Moreover, since ToPulledDownRefinement only adds non-migrations to (*T*^′^, *σ*^*′*^, *ℓ*^*′*^) to obtain 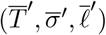, they have identical sets of migrations and comigrations, i. e. 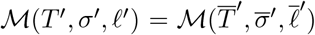 and 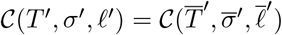, and thus identical numbers of migrations and comigrations. By Problem 1, 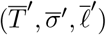 is a feasible solution to PMH-TR instance 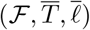 provided 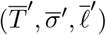 is a refinement of 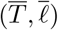. To that end, we show that 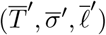 meets all the four conditions of refinement from Definition 2.

#### Algorithm S2

Construction of refinement 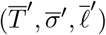 of 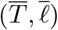 from given refinement (*T*^′^, *σ*^*′*^, *ℓ*^*′*^) of 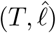

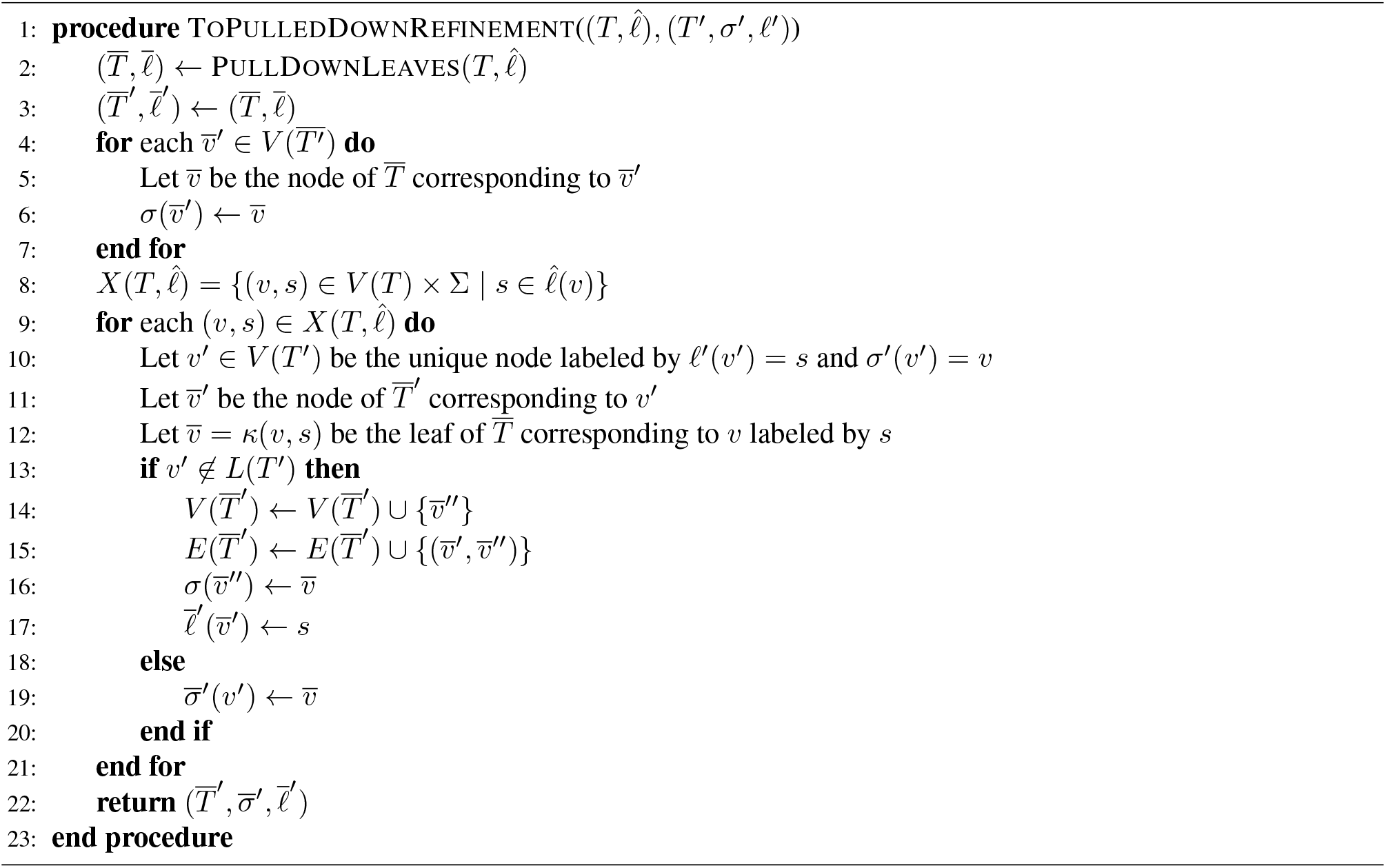

1. Condition (i) states that for any original node 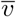 in 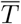, the refined nodes 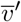 in 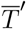 where 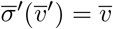 induce a connected subtree 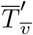 in 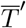. There are two cases.
  - 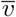 is in *T* . Note that 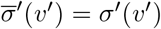 for any refined node *v*^*′*^ in *T*^′^ that is unomodified by ToPulledDown-Refinement. Since *T*^′^ is a refinement of *T*, the refined nodes *v*^*′*^ in *T*^′^ where 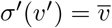 induce a connected subtree 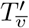 in *T*^′^. Moreover, as ToPulledDownRefinement adds or modifies leaves in *T*^′^ only, a leaf in *T*^′^ is also a leaf in 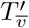, and removing leaves retains the tree structure, the nodes *v*^*′*^ in 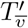 not modified by ToP-ulledDownRefinement induce a subtree both in *T*^′^ and 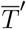. Therefore, to prove that 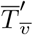 is an induced subtree, it suffices to show that each leaf 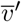 in 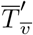 added or modified by ToPulledDownRefinement either is connected with an unmodified node from 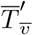, or 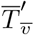 is a single-node subtree. Now two cases can arise.
    - 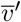 is in *T*^′^. In this case, 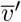 is the unique leaf with original node and location label being 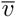 and 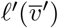, and for node-location pair 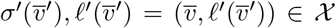, ToPulledDownRefinement modifies the corresponding original node of 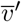 to be 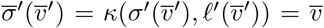. By definition, *κ*(*v, s*) = *v* for any original node *v* and location *s* if *v* is in *T* . Therefore, 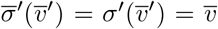. Thus, 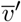 is part of subtree 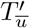 and so, either 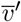 has an incoming edge from an unmodified node of 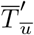 or 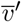 is the only node in 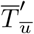.
    - 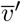 is not in *T*^′^. In this case, *v*^*′*^ is the unique leaf with original node and location label being 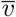 and *ℓ*^*′*^(*v*^*′*^), and for node-location pair 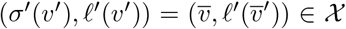, ToPulledDownRefinement adds the edge 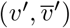. Thus, 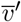 has an incoming edge from an unmodified node *v*^*′*^ of 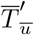. Therefore, 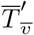 is a subtree.
  - 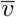 is a pulled-down leaf, i. e. 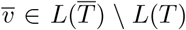. In this case, there is a unique node-location pair (*v*, ∈*s*) χ such that 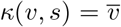 by definition. Consequently, ToPulledDownRefinement sets the original node of exactly one refined leaf 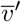 in 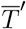 to 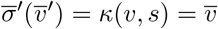. Thus, 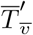 is a single-node tree.
2. Condition (ii) states that for each edge 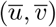 in 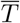, there is exactly one edge 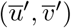 such that 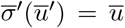 and 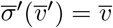. There are two cases.
  - 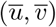 is in *T* . Since *T*^′^ is a refinement of *T*, there is exactly one edge (*u*^*′*^, *v*^*′*^) in *T*^′^ such that 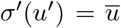 and 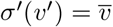. As ToPulledDownRefinement adds or modifies leaves only, and *u*^*′*^ is an internal node, 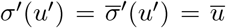. Moreover, since 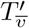 is a connected subtree, *v*^*′*^ must be the root of 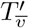. Now, if *v*^*′*^ is also an internal node, then 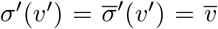. On the other hand, if *v*^*′*^ is a leaf, then 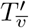 is a single-node tree, which implies 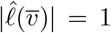. Let 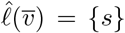. Then by PullDownLeaves, 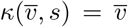, and so the corresponding original node in 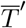 is 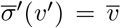. Note that since 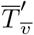 is a connected subtree, for any edge 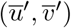 in *T*^′^ where 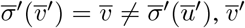 must be the root of 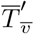. Since there can be only one root 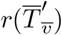, there is exactly one such edge in 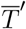.
  - 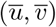 is not in *T* . In this case, 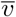 is a pulled-down leaf, and by PullDownLeaves, 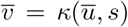, where 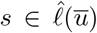. Let *u*^*′*^ be the unique node in *T*^′^ where 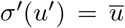 and *ℓ*^*′*^(*u*^*′*^) = *s*. If *u*^*′*^ is an internal node, ToPulledDownRefinement adds exactly one edge 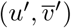 where 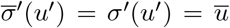 and 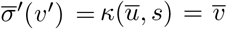. On the other hand, if *u*^*′*^ is a leaf, ToPulledDownRefinement modifies the original node of *u*^*′*^ to be 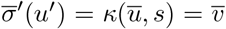. Moreover, since 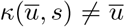, by PullDownLeaves, either 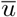 is internal in *T* or 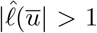. In both cases, at least one internal node *u*^″^ in *T*^′^ with 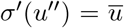 must be the parent of *u*^*′*^, as 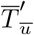 forms a connected subtree. As ToPulledDownRefinement does not modify internal nodes, 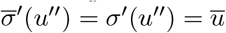.
3. Condition (iii) stating that there are no two nodes in 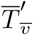 for any 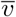 with the same label is satisfied, as (i) if 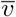 is in *T*, then 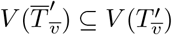 and there are no two nodes in 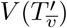, and (ii) if 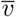 is not in *T*, then 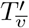 is a single-node tree.
4. Condition (iv) is satisfied because for each leaf 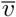 in 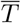 observed in 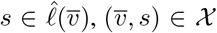, and ToPulledDown-Refinement adds a corresponding leaf 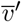 in 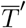 labeled 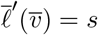.

Therefore, 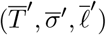 is a feasible solution to PMH-TR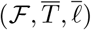.

Finally, we must show that 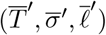 is mapped to uniquely by (*T*^′^, *σ*^*′*^, *ℓ*^*′*^). For a contradiction, assume ToP-ulledDownRefinement returns 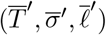 for two distinct refinements (*T*^′^, *σ*^*′*^, *ℓ*^*′*^) and (*T* ^″^, *σ*^″^, *ℓ*^″^) of 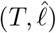. Now, since deleting the nodes location labeled by pulled-down leaves and the corresponding incoming edges added in 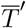 returns *T*^′^ and *T* ^″^, they are isomorphic. Also, since ToPulledDownRefinement does not change the location of the nodes simultaneously present in *T*^′^ or *T* ^″^, and 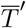, it holds that *ℓ*^*′*^ and *ℓ*^″^ are the same. As the construction also does not change any internal node, it holds that *σ*^*′*^(*v*^*′*^) = *σ*^″^(*v*^*′*^) for any internal node *v*^*′*^ in *T*^′^ and *T* ^″^. For a contradiction, assume *σ*^*′*^(*v*^*′*^) ≠ *σ*^″^(*v*^*′*^) for a leaf *v*^*′*^. Without loss of generality, three cases can happen.

- Both (*σ*^*′*^(*v*^*′*^), *ℓ*^*′*^(*v*^*′*^)) and (*σ*^″^(*v*^*′*^), *ℓ*^″^(*v*^*′*^)) are in 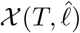. In this case, the construction sets the original node of *v*^*′*^ in 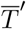 to be *κ*(*σ*^*′*^(*v*^*′*^), *ℓ*^*′*^(*v*^*′*^)) = *κ*(*σ*^″^(*v*^*′*^), *ℓ*^″^(*v*^*′*^)). But this is impossible if *σ*^*′*^(*v*^*′*^) ≠ *σ*^″^(*v*^*′*^).
- (*σ*^*′*^(*v*^*′*^), *ℓ*^*′*^(*v*^*′*^)) is in 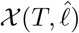 but (*σ*^″^(*v*^*′*^), *ℓ*^″^(*v*^*′*^)) is not. In this case, the construction sets the original node of *v*^*′*^ in 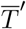 to be *κ*(*σ*^*′*^(*v*^*′*^), *ℓ*^*′*^(*v*^*′*^)) = *σ*^″^(*v*^*′*^). But that implies either (i) *σ*^*′*^(*v*^*′*^) = *σ*^*′*^(*v*^″^), which violates the premise, or (ii) *σ*^*′*^(*v*^*′*^) is the parent of *σ*^″^(*v*^*′*^), which is impossible as both are leaves.
- None of (*σ*^*′*^(*v*^*′*^), *ℓ*^*′*^(*v*^*′*^)) and (*σ*^″^(*v*^*′*^), *ℓ*^″^(*v*^*′*^)) are in χ. In this case, the construction sets the original node of *v*^*′*^ in 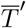 to be *σ*^*′*^(*v*^*′*^) = *σ*^″^(*v*^*′*^), which violates the premise.

Thus, *σ*^*′*^ = *σ*^″^, and therefore (*T*^′^, *σ*^*′*^, *ℓ*^*′*^) = (*T* ^″^, *σ*^″^, *ℓ*^″^).

To conclude the proof of Proposition 1, we present a function FromPulledDownRefinement to construct a feasible solution (*T*^′^, *σ*^*′*^, *ℓ*^*′*^) to PMH-TR instance 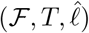 from 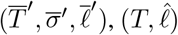, and 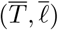, with the same number of migrations and comigrations, i. e. 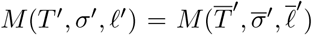 and 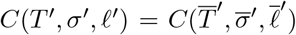. For each node 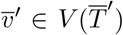 where (i) the original node 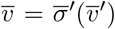 is a pulled-down leaf, i. e. 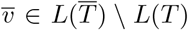, and (ii) the unique incoming edge 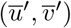 to 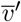 is a non-migration, i. e. 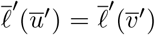, the function FromPulledDownRefinement constructs the clonal tree *T*^′^ by deleting node 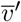 from 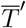 and attaching its children 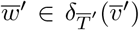 to its parent 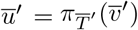. Afterwards, FromPulledDownRefinement constructs *σ*^*′*^ such that for any node 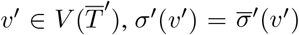 if 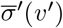 is in *T*, or 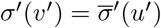 where *u*^*′*^ is the parent of *v*^*′*^ in 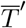 if the original node 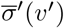 is a pulled-down leaf, i. e. 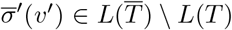. Last, FromPulledDownRefinement defines *ℓ*^*′*^(*v*^*′*^) to be 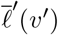 for all the nodes *v*^*′*^ in *T*^′^.

#### Algorithm S3

Construction of refinement (*T*^′^, *σ*^*′*^, *ℓ*^*′*^) of 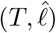 from given refinement 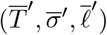 of 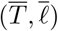

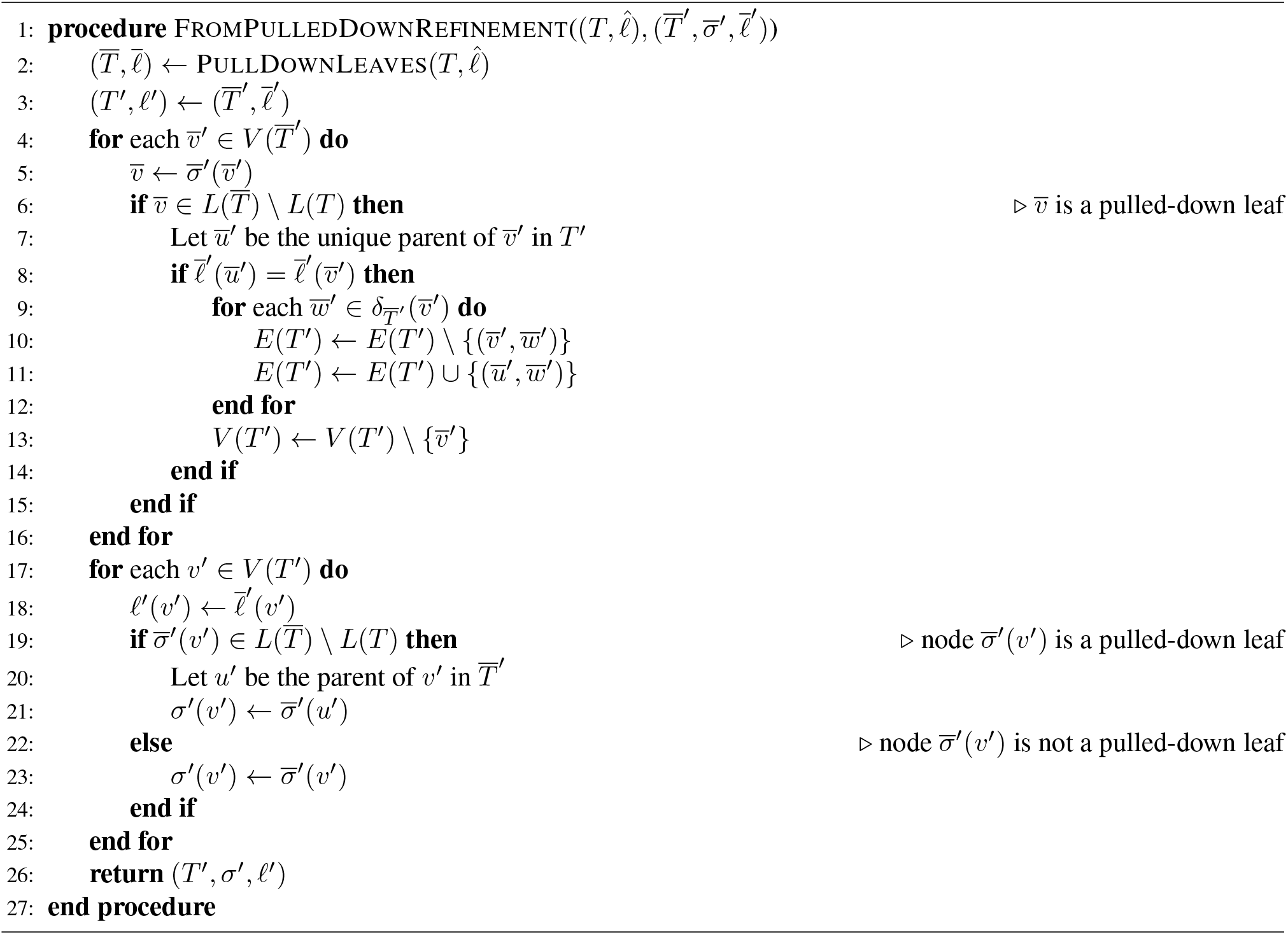

We now prove our second lemma, stating that for ℱ ∈ {MC, MC} the function FromPulledDownRefine-ment with domain PMH-TR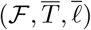 is a surjective function for the range of feasible solutions PMH-TR^*^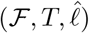with the same number of migrations and comigrations as solutions in PMH-TR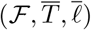.

#### Lemma 12.

*For* ℱ ∈ {MC, MC}, *the function* FromPulledDownRefinement *with domain* PMH-TR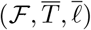 *is a surjective function for the range of feasible solutions* PMH-TR^*^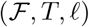*with the same number of migrations and comigrations as solutions in* PMH-TR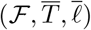.

*Proof*. For any refinement 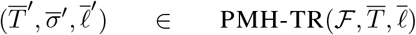, the function call 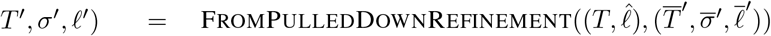 attaches children of *v*^*′*^ to the parent *u*^*′*^ of *v*^*′*^ after deleting *v*, effectively merging the edge (*u*^*′*^, *v*^*′*^) in tree 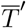 to obtain *T*^′^, *T*^′^ is a tree. Moreover, clearly *ℓ*^*′*^ is valid as it maps every node in *T*^′^ to a location from Σ. It remains to show that *σ*^*′*^ maps each node in *T*^′^ to a node in *T* . Assume for contradiction that node *v*^*′*^ in *T*^′^ is mapped to pulled-down leaf 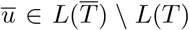. From FromPulledDownRefinement, this is only possible if the parent *u*^*′*^ of *v*^*′*^ is mapped to 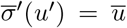. Again, since the nodes with original labels 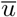 induce a subtree 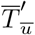 in 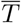 (condition (i) of Definition 2), and 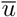 is a leaf in 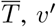 must be labeled with 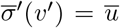 too in 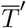. On the other hand, the incoming edge (*w*^*′*^, *u*^*′*^) in 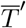 is a migration, otherwise *u*^*′*^ would be deleted. Assume 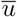 is observed in the unique location *s* in 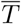 by PullDownLeaves. Now, if *ℓ*^*′*^(*u*^*′*^) = *s*, then deleting migration (*u*^*′*^, *v*^*′*^) decreases the number of migrations in 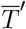 by one. Again, if *ℓ*^*′*^(*v*^*′*^) = *s*, then deleting node *u*^*′*^ and attaching *v*^*′*^ as a child of *w*^*′*^ also decreases the number of migrations in 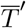 by one. Since both the cases violate the premise that 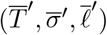 is optimal, *v*^*′*^ in *T*^′^ cannot have pulled-down leaf 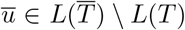 as the original node. Thus, *σ*^*′*^ is valid.

Since FromPulledDownRefinement only deletes non-migrations from 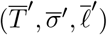 to obtain (*T*^′^, *σ*^*′*^, *ℓ*^*′*^), they have identical sets of migrations and comigrations, i. e. 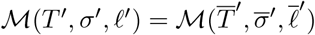 and 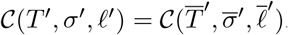. By Problem 1, (*T*^′^, *σ*^*′*^, *ℓ*^*′*^) is a feasible refinement to PMH-TR instance 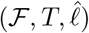 if (*T*^′^, *σ*^*′*^, *ℓ*^*′*^) is a refinement of 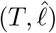. To that end, we show that (*T*^′^, *σ*^*′*^, *ℓ*^*′*^) meets all the four conditions of refinement from Definition 2.

1. For condition (i), we prove for any original node *v* in *T*, the nodes *v*^*′*^ in *T*^′^ where *σ*^*′*^(*v*^*′*^) induce a connected subtree 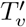 in *T*^′^. From FromPulledDownRefinement, 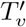 consists of all the nodes from 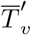, and the nodes *v*^*′*^ with incoming edge (*u*^*′*^, *v*^*′*^) such that *v*^*′*^ has a pulled-down leaf as original node in 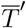, i. e. 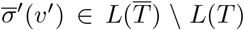, (*u*′ *v*′) is a migration, and *u*^*′*^ is in 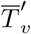. Thus, 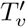 is a connected subtree in *T*^′^.
2. Condition (ii) states that for each edge (*u, v*) in *T*, there is a refined edge (*u*^*′*^, *v*^*′*^) in *T*^′^ such that *σ*^*′*^(*u*^*′*^) = *u* and *σ*^*′*^(*v*^*′*^) = *v*. Now by PullDownLeaves, each edge (*u, v*) of *T* is present in 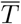, which ensures exactly one edge 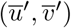 in 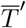 where 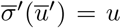 and 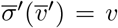. Since 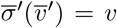 is in *T*, FromPulledDownRefinement keeps the edge 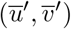 in *T*^′^ without changing the corresponding original nodes, i. e. 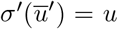 and 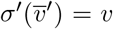. Thus, condition (ii) is satisfied.
3. Condition (iii) states that for any two nodes *v*^*′*^, *v*^″^ with the same original node *σ*^*′*^(*v*^*′*^) = *σ*^*′*^(*v*^″^) = *v* have different location labels, i. e. *ℓ*^*′*^(*v*^*′*^) *ℓ*^*′*^(*v*^″^). Suppose for contradiction that 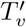 has two nodes *v*^*′*^, *v*^″^ with the same label *ℓ*^*′*^(*v*^*′*^) = *ℓ*^*′*^(*v*^″^) = *s*. Since 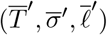 is a refinement of 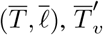 cannot have two nodes with the same label. Thus, both *v*^*′*^ and *v*^″^ cannot be in 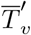. Without loss of generality, assume *v*^*′*^ with incoming edge (*u*^*′*^, *v*^*′*^) is in 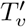 but not in 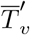. Then by FromPulledDownRefinement, *v*^*′*^ has a pulled-down leaf as original node in 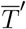, i. e. 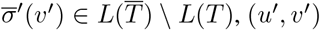 is a migration, and *u*^*′*^ is in 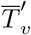. Now deleting the edge (*u*^*′*^, *v*^*′*^) and adding (*v*^″^, *v*^*′*^) decreases the number of migrations by one, which contradicts with the premise that 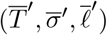 is an optimal refinemtnt to PMH-TR instance 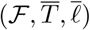. So condition (iii) is satisfied.
4. Condition (iv) states that for each node-location pair (*v, s*) where 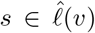, there is a node *v*^*′*^ in 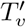 such that *ℓ*^*′*^(*v*^*′*^) = *s*. Assume for any node-location pair (*v, s*) where 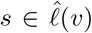 that 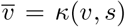 is a leaf in 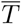 with label 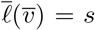. Thus there is a node 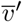 in 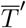 such that 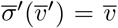 and 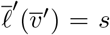. By FromPulledDownRefine-ment, 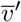 is deleted if the incoming edge 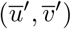 to 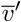 is a non-migration, which ensures there is a node 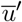 labeled by 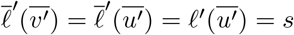 in 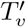. On the other hand, if 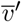 is not deleted, then 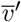 labeled by 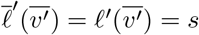 is a part of 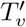. Thus, condition (iv) is satisfied.

Therefore, (*T*^′^, *σ*^*′*^, *ℓ*^*′*^) is a feasible solution to PMH-TR instance 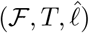.

Last, we prove that FromPulledDownRefinement(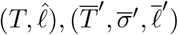,) is a surjective function i. e. for any feasible optimal solution (*T*^′^, *σ*^*′*^, *ℓ*^*′*^) to PMH-TR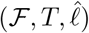, there is a refinement 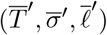 such that 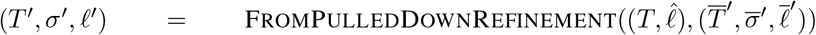. We assume for contradiction that there is no refinement 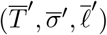 for 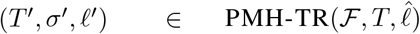 such that 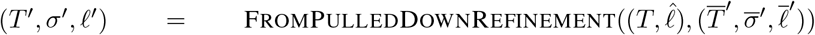. Let 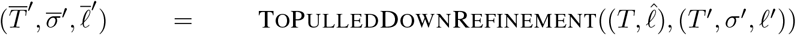, and 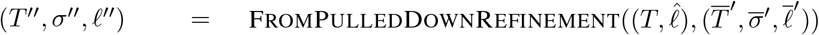, (*T ′, σ*^*′*^, **ℓ*′*)). To prove that 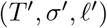 and (*T* ^″^, *σ*^″^, *ℓ*^″^) are the same, it suffices to show that for each edge (*u*^*′*^, *v*^*′*^) in *T*^′^, (i) (*u*^*′*^, *v*^*′*^) is in *T* ^″^, (ii) *σ*^*′*^(*u*^*′*^) = *σ*^″^(*u*^*′*^) and *σ*^*′*^(*v*^*′*^) = *σ*^″^(*v*^*′*^), and (iii) *ℓ*^*′*^(*u*^*′*^) = *ℓ*^″^(*u*^*′*^) and *ℓ*^*′*^(*v*^*′*^) = *ℓ*^″^(*v*^*′*^).

We begin by showing that for each edge (*u*^*′*^, *v*^*′*^) in *T*^′^, (i) (*u*^*′*^, *v*^*′*^) is in *T* ^″^. By ToPulledDownRefinement, (*u*^*′*^, *v*^*′*^) is in 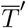. Assume for a contradiction that (*u*^*′*^, *v*^*′*^) is deleted by FromPulledDownRefinement in *T* ^″^. This is possible only if 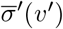 is a pulled-down leaf, and (*u*^*′*^, *v*^*′*^) is a non-migration. As 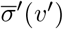 is a pulled-down leaf, 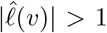 or *v* is an internal node in *T*, where *v* = *σ*^*′*^(*v*^*′*^). But if *v* is an internal node, *v*^*′*^ cannot be labeled with a pulled down leaf. On the other hand, if 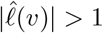, then *u*^*′*^ must also be in 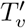, and thus (*u*^*′*^, *v*^*′*^) is a migration, leading to a contradiction. Thus for each edge (*u*^*′*^, *v*^*′*^) in *T*^′^, (*u*^*′*^, *v*^*′*^) is in *T* ^″^.

Next we show that for any node *u*^*′*^ in *T*^′^, it holds that *σ*^*′*^(*u*^*′*^) = *σ*^″^(*u*^*′*^). By ToPulledDownRefinement, either 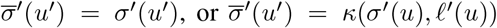. In both cases, FromPulledDownRefinement assigns *σ*^″^(*u*^*′*^) = *σ*^*′*^(*u*^*′*^).

Last, we show that *ℓ*^*′*^(*v*^*′*^) = *ℓ*^″^(*v*^*′*^) for any node *v*^*′*^ in *T*^′^. This is true as both ToPulledDownRefinement and FromPulledDownRefinement retain the labels of the clones present in both input and output. Therefore, FromPulledDownRefinement is a surjective function.

**(Main text) Theorem 2**. *There exists a bijection between refinements* (*T*^′^, *σ*^*′*^, *ℓ*^*′*^) *and pairs* (*R, e*) *of edge-labeled refinement graphs for a given tree T with observed labeling* 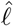.

First, we show that given a pair (*R, e*) of refinement graph *R* with edge labeling *e*, there exists a unique refinement (*T*^′^, *σ*^*′*^, *ℓ*^*′*^). We present a construction of (*T*^′^, *σ*^*′*^, *ℓ*^*′*^) from (*R, e*) given 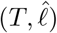 in Algorithm S4.

#### Algorithm S4

Construction of (*T*^′^, *σ*^*′*^, *ℓ*^*′*^) from (*R, e*)

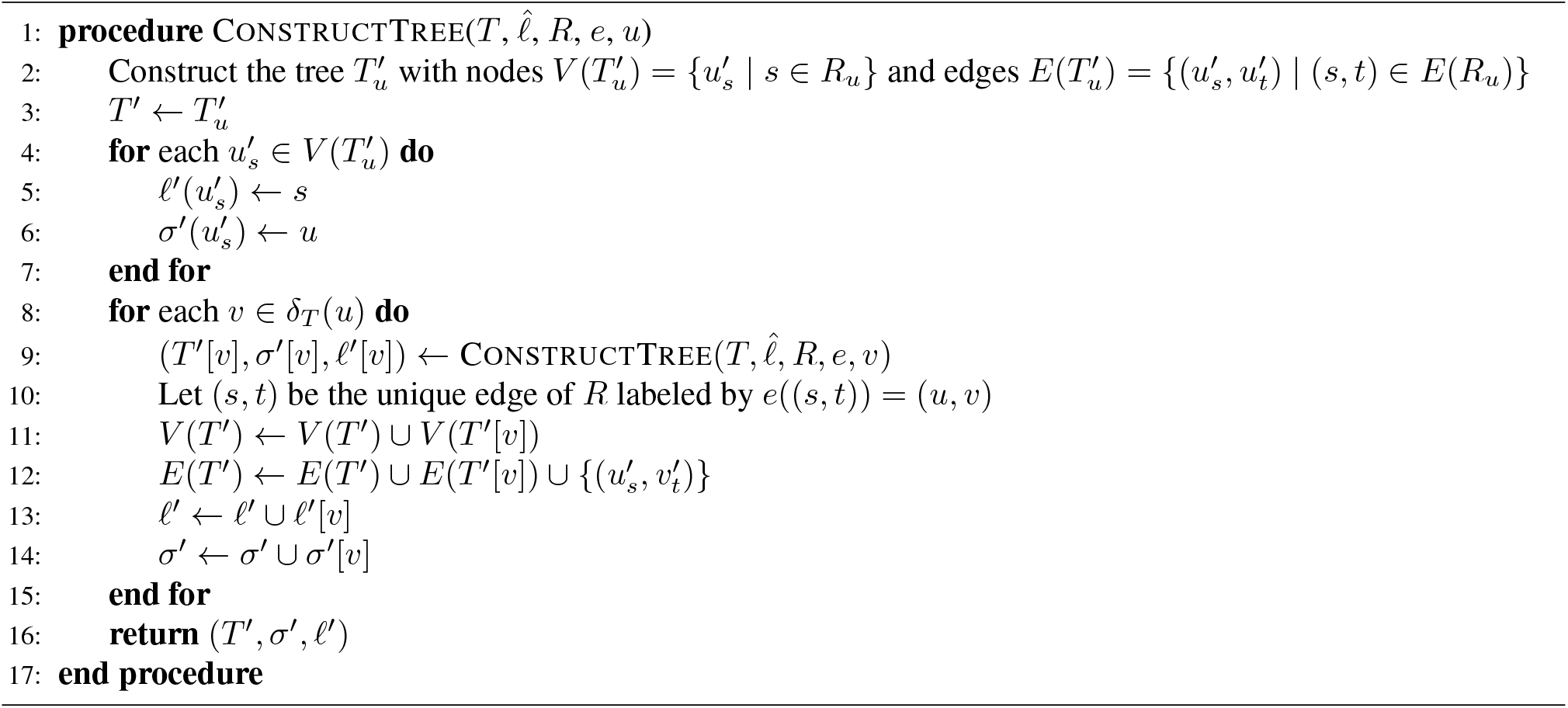

Given an edge-labeled refinement graph (*R, e*), we now prove that ConstructTree 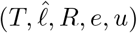returns a refinement (*T*^′^, *σ*^*′*^, *ℓ*^*′*^) for the subtree *T* [*u*] of *T* rooted at *u*. As such, ConstructTree 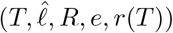 will yield a refinement for 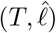. Recall that 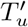 is the subtree of *T*^′^ induced by the nodes

#### Lemma 13.

*For any node u* ∈ *V* (*T*), *the tuple* (*T*^′^, *σ*^*′*^, *ℓ*^*′*^) *returned by* ConstructTree 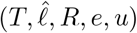 *is a refinement for the subtree T* [*u*] *of T rooted at node u. Moreover, the refined tree T*^′^ *is rooted at the node* 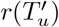.

*Proof*. We present a proof by structural induction on *T* .

*(Base case)* For the base case, *u* is a leaf in *T* labeled by 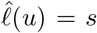. We prove the base case by showing that (i) *T*^′^ is a tree rooted at 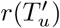, and (ii) *T*^′^ is a refinement of the single node tree *T* [*u*]. For a leaf *u* without any children, *T*^′^ equals 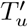 (see Line 2-3 and Line 8 of Algorithm S4), which is clearly isomorphic to *R*_*u*_ (Line 2 of Algorithm S4). Since *R*_*u*_ is a tree by Definition 9, *T*^′^ is a tree rooted at 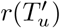.

To prove *T*^′^ to be a refinement of *T* [*u*], we show that *T*^′^ maintains all four conditions of refinement from Definition 2. First, condition (i) states that 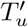 induces a tree in *T*^′^, which is true as 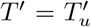. Second, condition (ii) states that for each original edge (*u, v*) in *T*, there is exactly one edge (*u*^*′*^, *v*^*′*^) in *T*^′^ such that *σ*^*′*^(*u*^*′*^) = *u* and *σ*^*′*^(*v*^*′*^) = *v*. This is trivially satisfied, as *T* [*u*] does not have any edge. Third, condition (iii) states that 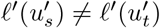 for any two nodes 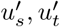 in 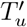, which is satisfied as the construction adds a unique node 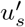 with label 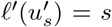 for each node *s* ∈ *V* (*R*_*u*_) (Line 2 and Line 5 of Algorithm S4). Fourth, condition (iv) states that there is a node 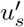 in 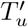 with label 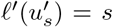 for each 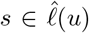. From Definition 9, 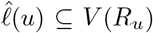, which means for each location 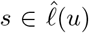, there is a node *s* in *R*_*u*_, which adds the node 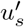 in 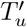 with label 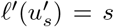 (Line 2 and Line 5 of Algorithm S4). Hence, (*T*^′^, *σ*^*′*^, *ℓ*^*′*^) is a refinement for the subtree *T* [*u*] of *T* and *T*^′^ is rooted at 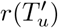.

*(Induction step)* In this step, we have that *u* is an internal node of *T* . The proof proceeds in four steps: showing that (i) *T*^′^ is a tree rooted at 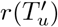, and (ii) (*T*^′^, *σ*^*′*^) is a refinement of *T* [*u*].

First, to prove *T*^′^ is a tree, notice that *T*^′^ is initialized in Line 3 by 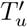. Thus we begin by showing that 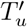 constructed in Line 2 of ConstructTree is a rooted tree. From Line 2, it is clear that 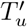 is isomorphic to *R*_*u*_. Since *R*_*u*_ is a directed tree in *R* by Definition 9, 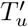 is a tree. In Line 4-15, ConstructTree iterates over each child *v* of *u* in *T* and adds *T*^′^[*v*] and the edge 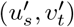 to *T*^′^, where (*s, t*) is the unique edge in *R* labeled by *e*((*s, t*)) = (*u, v*). We show that *T*^′^ remains a tree rooted at 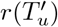 after each iteration processing node *v*∈ *δ*_*T*_ (*u*) (Line 8). By the induction hypothesis, *T*^′^[*v*] is a tree rooted at 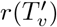. The edge 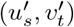 added in Line 12 of ConstructTree connects a node 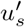 from 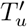 with the root 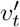 of *T*^′^[*v*]. By Definition 9, if *e*((*s, t*)) = (*u, v*), then *s ∈ V* (*R*_*u*_) (since *s* has an outgoing edge labeled by (*u, v*)), which implies that 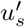 is a node in 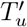. On the other hand, *e*((*s, t*)) = (*u, v*) implies *t* = *r*(*R*_*v*_), and so 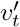 is the root of 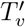 since 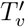 and *R*_*v*_ are isomorphic, and by extension, the root of *T*^′^[*v*]. Thus, *T*^′^ remains a tree rooted at 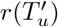 after each iteration.

Second, to show that (*T*^′^, *σ*^*′*^, *ℓ*^*′*^) is a refinement of 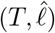, we prove that the four conditions of refinement from Definition 2 are met.

1. Condition (i) states that 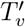 induced by nodes *v*^*′*^ where *σ*^*′*^(*v*^*′*^) = *v* for any internal node *v* in *T* [*u*] is a tree. Clearly, 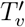 generated in Line 2 of ConstructTree for any *v*∈ *T* [*u*] comprises all nodes *v*^*′*^ such that *σ*^*′*^(*v*^*′*^) = *v*. Given that 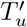 is a tree and by induction hypothesis, it follows that 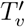 for any internal node *v* in *T* [*u*] is a tree.
2. Condition (ii) states that for each edge 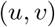 in *T*, there is exactly one edge 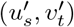 such that 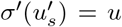 and 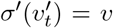. Note that any edge of *T*^′^ is either 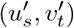 where *e*((*s, t*)) = (*u, v*), or in 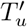 or *T*^′^[*v*] for any child *v* of *u* in *T* . From ConstructTree, clearly *σ*^*′*^(*u*^*′*^) = *u* for every node *u*^*′*^ in 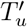 and there can not be any node *w*^*′*^ in *T*^′^[*v*] where *σ*(*w*^*′*^) = *u*. Thus there cannot be any edge 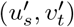 in 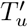 or *T*^′^[*v*] such that 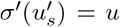 and 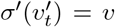. By Definition 9, there is exactly one edge (*s, t*) in *R* labeled by *e*((*s, t*)) = (*u, v*) for each (*u, v*), which causes exactly one edge 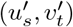 to be added in *T*^′^ by ConstructTree, where 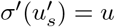 and 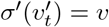.
3. Condition (iii) states that for any two nodes 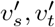 in 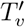 for any original node *v* ∈*V* (*T*), it holds that *ℓ*^*′*^(*u*^*′*^) ≠ *ℓ*^*′*^(*v*^*′*^). Now, ConstructTree adds a unique node 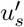 with label 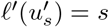 for each node *s* ∈*V* (*R*_*u*_) (Line 2 and Line 5), which renders each node of 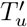 to have unique label. Thus, given the induction hypothesis, condition (iii) is met for any original node *v* ∈ *V* (*T*).
4. Condition (iv) states that for each node-location pair (*v, s*) such that 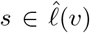, there is a refined node *v*^*′*^ where the corresponding original node is *σ*^*′*^(*v*^*′*^) = *v* and the location label is *ℓ*^*′*^(*v*^*′*^) = *s*. Now, by condition (iii) of the definition of a refinement graph (Definition 9), *s* ∈ *V* (*R*_*v*_) if 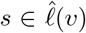. Consequently, ConstructTree adds a unique node 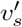 with label 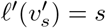 node-location pair (*v, s*) such that 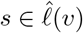 (Line 2 and Line 5).

Therefore, (*T*^′^, *σ*^*′*^, *ℓ*^*′*^) is a refinement. □

We next show that given a refinement (*T*^′^, *σ*^*′*^, *ℓ*^*′*^) comprised of a refinement (*T*^′^, *σ*^*′*^) and location labeling *ℓ*^*′*^, there exists a unique edge-labeled refinement graph (*R, e*). We present a construction of (*R, e*) from *T*^′^, *σ*^*′*^, and *ℓ*^*′*^ given 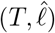 in Algorithm S5. We prove in the following lemma that (*R, e*) is an edge-labeled refinement graph.

#### Lemma 14.

ConstructGraph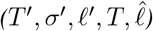 *returns an edge-labeled refinement graph* (*R, e*) *for* 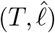.

*Proof*. From ConstructGraph, clearly *V* (*R*) = Σ, and the edge labeling *e* labels the edges of *R* either with a clonal tree node from *V* (*T*), or a clonal tree edge from *E*(*T*). Thus to show that (*R, e*) is an edge-labeled refinement graph, it suffices to prove that (*R, e*) adheres to all five conditions from Definition 9.

1. By Definition 2, for each edge (*u, v*) in *T*, there exists exactly one edge 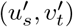 in *T*^′^ such that 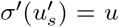 and 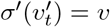, and thus ConstructGraph adds exactly one edge (*s, t*) in *R* where 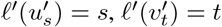, labeled by *e*((*s, t*)) = (*u, v*), satisfying Condition (i) of Definition 9.
2. Following from the previous point, if *v* is a leaf in *T*, then 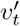 is a leaf in *T*^′^ by Definition 2, which implies 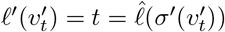 according to Problem 1. Thus condition (ii) of Definition 9 is fulfilled.
3. *R*_*u*_ is a tree as it is clearly isomorphic to 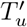 by Line 4 of ConstructGraph, and the edges (*s, t*) of *R*_*u*_ are labeled with *e*((*s, t*)) = *u* for any *u*∈ *I*(*T*). From ConstructGraph, clearly if node 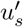 is in 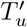, then 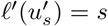 is in *R*_*u*_. Now since 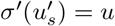 is internal in 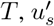 cannot be a leaf in *T*^′^ by Definition 2. As a result, there exists at least one outgoing edge 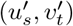 in *T*^′^. Now if 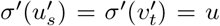, then edge 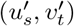 is in 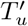, and so *s* has an outgoing edge 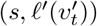 in isomorphic *R*_*u*_. On the other hand, if 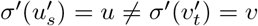, then from Definition 2, it follows that (*u, v*) is an edge in *T*, and edge 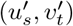 is the only edge of *T*^′^ where 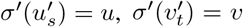. Consequently, ConstructGraph adds edge 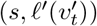 labeled by 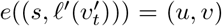 in *R*. So each node *s* of *R*_*u*_ has at least one outgoing edge in *R* labeled either by node *u* ∈*V* (*T*) or edge (*u, v*) ∈*E*(*T*). Thus condition (iii) of Definition 9 is satisfied.
4. Since (*T*^′^, *σ*^*′*^, *ℓ*^*′*^) is a refinement, for any two nodes 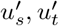 in *T*^′^ such that 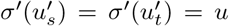, both 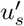 and 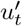 have at least two outgoing edges in *T*^′^. Now 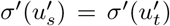 implies nodes 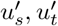 are in 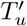. In this case, ConstructGraph adds nodes 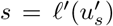 and 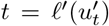 to *R*_*u*_. Now if 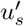 has an outgoing edge 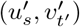, then ConstructGraph adds edge 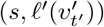 and labels it with *u* if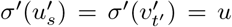, or with (*u, v*) if 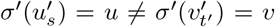. Thus for two outgoing edges of 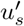 and 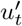, ConstructGraph adds two outgoing edges from *s* and *t* labeled either by *u* or (*u, v*). Thus condition (iv) of Definition 9 is satisfied.
5. From Definition 2, for any edge (*u, v*) in *T*, there exists exactly one edge 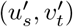 in *T*^′^ where 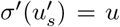 and 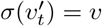. Since 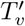 is a subtree of tree 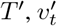 must be the root of 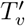 Since 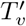 and *R*_*v*_ are isomorphic, 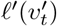 is the root of directed tree *R*_*v*_ and ConstructGraph adds edge 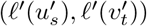 labeled by 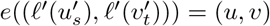 in *R*. Thus, condition (v) of Definition 9 is satisfied. Therefore, (*R, e*) is an edge-labeled refinement graph. □

#### Algorithm S5

Construction of (*R, e*) from (*T*^′^, *σ*^*′*^, *ℓ*^*′*^)

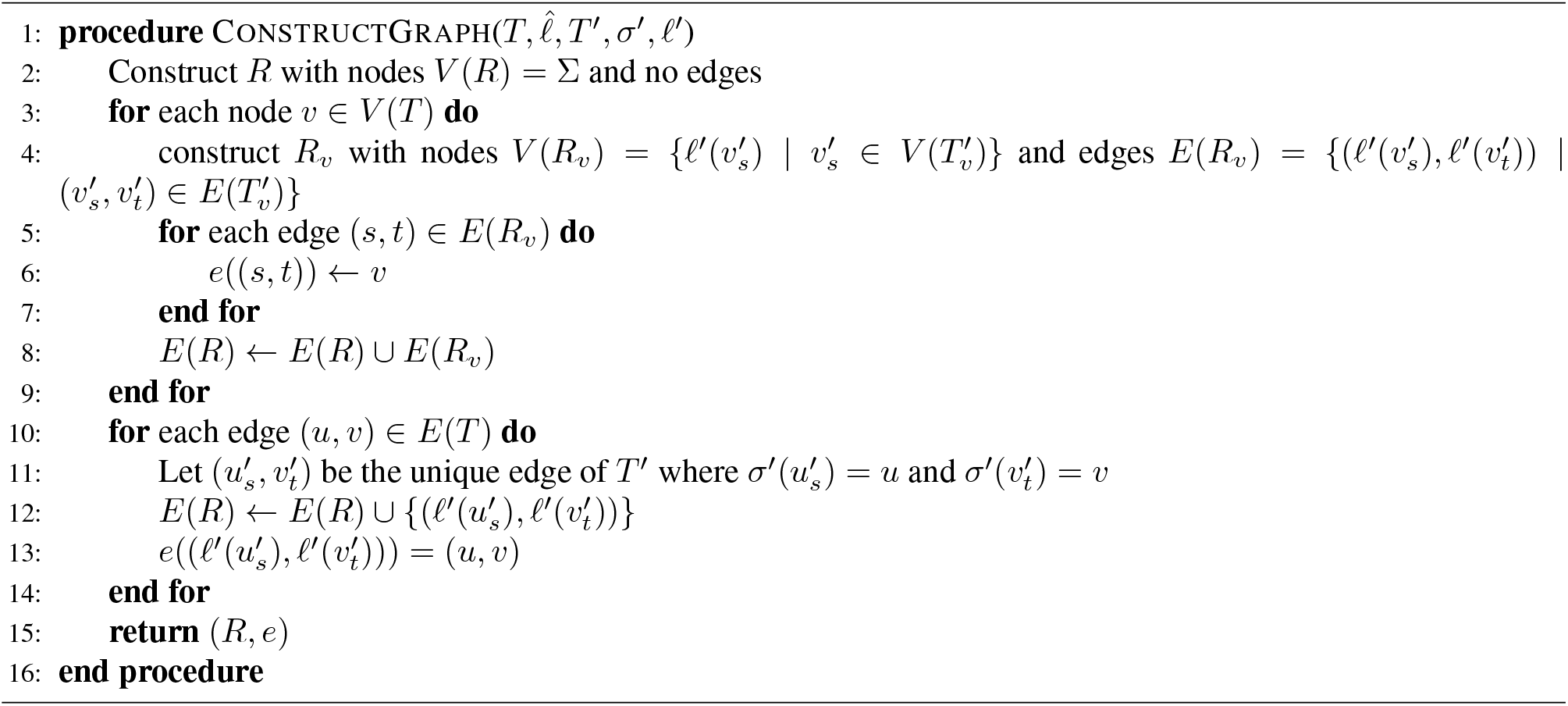

#### Proposition 15.

*Let* (*R, e*) *be a refinement graph and* (*T* ′, *σ*′, *ℓ*′) *be a refinement for a given a tree T with leaf labeling* 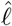. *Then*, ConstructTree*(T*, 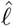, *R, e, r*(*T*)*) returns tuple* (*T*′, *σ*′, *ℓ*′) *if and only if* ConstructGraph*(T*, 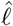, *T* ′, *σ*′, *ℓ*′*) returns* (*R, e*).

*Proof*. (⇒) Let (*T*′, *σ*′, *ℓ*′) be the tuple returned by ConstructTree(*T*, 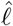, *R, e, r*(*T*)). Assume for a contradiction that ConstructGraph(*T*, 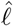, *T* ′, *σ*′, *ℓ*′) returns a pair (*R*′, *e*′) distinct from (*R, e*). By Definition 9, each edge (*s, t*) of *R* is labeled either by node *v* ∈ *V* (*T*) or edge (*u, v*) ∈ *E*(*T*). Now, if any edge (*s, t*) of *R* is labeled by *e*((*s, t*)) = *u* for any node *u* ∈ *V* (*T*), then (*s, t*) ∈ *R*_*u*_ by Definition 9. Thus, ConstructTree(*T*, 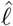, *R, e, u*) adds edge 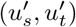 to *T* ′ and sets 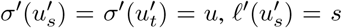, and 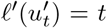. But then ConstructGraph(*T*, 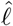, *T* ′, *σ*′, *ℓ*′) adds 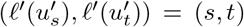 labeled by *e*′((*s, t*)) = *u* to *R*′. On the other hand, if any edge (*s, t*) of *R* is labeled by *e*((*s, t*)) = (*u, v*) for any edge (*u, v*) ∈ *E*(*T*), then ConstructTree(*T*, 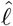, *R, e, u*) adds edge 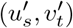 to *T* ′ and sets 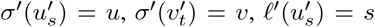, and 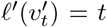. But then 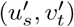 is the unique edge of *T* ′ where 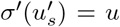 and 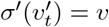 by Definition 2, and thus ConstructGraph(*T*, 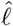, *T*′, *σ*′, *ℓ*′) adds 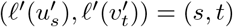 labeled by *e*′((*s, t*)) = (*u, v*) to *R*′. Therefore, *R* and *R*′ are isomorphic and *e* = *e*′, which leads to a contradiction.

(⇐) Let (*R, e*) be the pair returned by ConstructGraph(*T*, 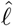, *T* ′, *σ*′, *ℓ*′), and assume for a contradiction that ConstructTree(*T*, 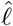, *R, e, r*(*T*)) returns (*T* ″, *σ*″, *ℓ*″) that is different from (*T* ′, *σ*′, *ℓ*′). Now for any edge 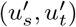 in *T* ′ where 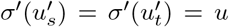 for any *u* ∈ *V* (*T*) and 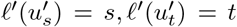, it holds that 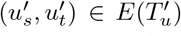 by Definition 2. Thus, ConstructGraph(*T*, 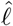, *T* ′, *σ*′, *ℓ*′) adds edge (*s, t*) labeled by *e*((*s, t*)) = *u* to *R*. But then (*s, t*) ∈ *E*(*R*_*u*_) by Definition 9, and so ConstructTree(*T*, 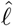, *R, e, u*) adds edge 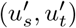 to *T* ″ and sets 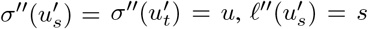, and 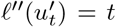. On the other hand, for any edge 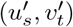 in *T* ′ such that 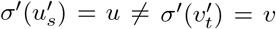 and 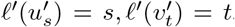, ConstructGraph(*T*, 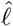, *T* ′, *σ*′, *ℓ*′) adds edge (*s, t*) labeled by *e*((*s, t*)) = (*u, v*) to *R*. But then ConstructTree(*T*, 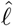, *R, e, u*) adds edge 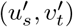 to *T* ″ and sets 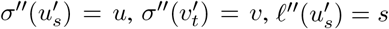, and 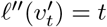. Therefore, *T* ′ and *T* ″ are isomorphic, and *σ*″ and *ℓ*″ are the same as *σ*′ and *ℓ*′ respectively, leading to a contradiction. □

**(Main text) Proposition 7**. *It holds that* 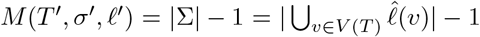 *if and only if (i) ℓ*′(*v*′) ∈ Σ *for each node v*′ *of T*′ *and (ii) for any two nodes w*′, *w*″ *of T*′ *where ℓ*′(*w*′) = *ℓ*′(*w*″), *each node in the undirected path from w*′ *to w*″ *is labeled with ℓ*′(*w*′) = *ℓ*′(*w*″).

*Proof*. (⇒) We prove the lemma in three steps — showing that (i) 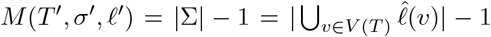 implies there is a bijection between the migrations (*u*′, *v*′) in *T* ′ and the locations *s* from Σ where *s* ≠ *ℓ*′(*r*(*T* ′)), (ii) *ℓ*′(*v*′) ∈ Σ for each node *v*′ of *T* ′, and (iii) for any two nodes *w*′, *w*″ of *T* ′ where *ℓ*′(*w*′) = *ℓ*′(*w*″), each node in the undirected path from *w*′ to *w*″ is labeled with *ℓ*′(*w*′) = *ℓ*′(*w*″).

First, we prove that 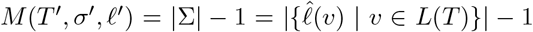 implies there is a bijection between the migrations (*u*′, *v*′) in *T* ′ and the locations *s* from Σ where *s* ≠ *ℓ*′(*r*(*T* ′)}. To be more specific, we show that there is exactly one unique migration (*u*′, *v*′) for each of |Σ| − 1 locations *s* ∈ Σ\*ℓ*′(*r*(*T* ′)} such that *ℓ*′(*v*′) = *s*, and for each migration (*u*′, *v*′) in *T*′, *ℓ*′(*v*′) is in Σ and not the label of the root *r*(*T* ′). Consider any location *s* ∈ Σ where *s* is not the label of the root *r*(*T* ′) and leaf *w*′ in *T* ′ is labeled with *ℓ*′(*w*′) = *s*. Then there is at least one migration (*u*′, *v*′) in the path from root *r*(*T* ′) to leaf *w*′ such that *ℓ*′(*v*′) = *s*. Given *M* (*T* ′, *σ*′, *ℓ*′) = |Σ| − 1, each of the |Σ| − 1 locations *s* ∈ Σ *ℓ*′(*r*(*T* ′)) can have exactly one migration (*u*′, *v*′) with *ℓ*′(*v*′) = *s*, and for each migration (*u*′, *v*′) in *T* ′, *ℓ*′(*v*′) is in Σ \*ℓ*′(*r*(*T* ′)} .

Second, we show that *ℓ*′(*v*′) ∈ Σ for each node *v*′ of *T* ′. Assume for contradiction that *ℓ*′(*v*′) ∉ Σ for a node *v*′ of *T* ′. Now two cases can happen.

- *v*′ = *r*(*T*′) : Since each location *s* from Σ\{*ℓ*′(*r*(*T* ′))} = Σ\{*ℓ*′(*u*′)} = Σ has at least one migration (*u*′, *v*′) with *ℓ*′(*v*′) = *s*, the number *M* (*T* ′, *σ*′, *ℓ*′) of migrations is at least |Σ|, contradicting the premise that *M* (*T* ′, *σ*′, *ℓ*′) = |Σ| − 1.
- *v*′≠ *r*(*T* ′) : In this case, *ℓ*′(*r*(*T* ′)) ∈ Σ, and there exists at least one migration (*u*″, *v*″) in the path from root *r*(*T* ′) to *v*′ such that *ℓ*′(*v*″) = *ℓ*′(*v*′). Since each of |Σ| − 1 locations *s* ∈ Σ\{*ℓ*′(*r*(*T* ′))} has at least one migration (*u*′, *v*′) with *ℓ*′(*v*′) = *s*, the total number *M* (*T* ′, *σ*′, *ℓ*′) of migrations stands at least |Σ| after including the migration (*u*″, *v*″), contradicting the premise that *M* (*T* ′, *σ*′, *ℓ*′) = |Σ| − 1.

Third, we prove that for any two nodes *w*′, *w*″ in *T* ′ where *ℓ*′(*w*′) = *ℓ*′(*w*″), each node in the undirected path from *w*′ to *w*″ is labeled with *ℓ*′(*w*′) = *ℓ*′(*w*″). Let *u*′ be the most recent common ancestor of any two nodes *w*′, *w*″ in *T* ′ where *ℓ*′(*w*′) = *ℓ*′(*w*″). Thus, the path from *w*′ to *w*″ comprises the nodes along the path from *u*′ to *w*′ and from *u*′ to *w*″, and no edge can be both in the path from *u*′ to *w*′ and from *u*′ to *w*″. We begin by showing that *ℓ*′(*u*′) = *ℓ*′(*w*′) = *ℓ*′(*w*″), and then we show the nodes in the path from *u*′ to *w*′ and from *u*′ to *w*″ are labeled with *ℓ*′(*w*′) = *ℓ*′(*w*″). Assume for a contradiction that *ℓ*′(*u*′) *ℓ*′(*w*′) ≠ *ℓ*′(*w*″). Then clearly *u*′ ≠ *w*′ and *u*′ ≠ *w*″, and therefore, there is at least one migration (*u*″, *v*″) in the path from *u*′ to *w*′ and at least one migration (*u*‴, *v*‴) different from (*u*″, *v*″) in the path from *u*′ to *w*″ where *ℓ*′(*v*″) = *ℓ*′(*w*′) = *ℓ*′(*v*‴) = *ℓ*′(*w*″). But then there are two migrations (*u*″, *v*″) and (*u*‴, *v*‴) where *ℓ*′(*v*″) = *ℓ*′(*v*‴), contradicting that there is a bijection between the migrations in *T* ′ and the locations *s* from Σ where *s* ≠ *ℓ*′(*r*(*T* ′)). Thus, *ℓ*′(*u*′) = *ℓ*′(*w*′) = *ℓ*′(*w*″).

(⇐) To prove the reverse direction, we assume the conditions in the lemma are true, and prove that *M* (*T* ′, *σ*′, *ℓ*′) = |Σ| − 1. From Proposition 6, *M* (*T*′, *σ*′, *ℓ*′) ≥ |Σ| − 1. For contradiction, assume *M* (*T* ′, *σ*′, *ℓ*′) *>* |Σ| − 1. Then by the pigeonhole principle, there is at least one location *s* ∈ Σ such that either (i) there are at least two migrations (*u*′, *v*′) and (*u*″, *v*″) such that *ℓ*′(*v*′) = *ℓ*′(*v*″) = *s* ≠ *ℓ*′(*u*′), or (ii) the root *r*(*T* ′) is labeled with *ℓ*′(*r*(*T* ′)) = *s* and additionally, there is at least one migrations (*u*′, *v*′) such that *ℓ*′(*v*′) = *s*/= *ℓ*′(*u*′). In both cases, node *u*′ is in the path from two nodes labeled by *s* in tree *T* ′, which contradicts with condition (ii) of the premise. Thus, *M* (*T* ′, *σ*′, *ℓ*′) = |Σ| − 1. □

**(Main text) Proposition 8**. *It holds that C*(*T* ′, *σ*′, *ℓ*′) ≤ |*E*(*R*)| ≤ |*E*(*T*)| + |*V* (*T*)|(|Σ| − 1).

*Proof*. Clearly *C*(*T* ′, *σ*′, *ℓ*′) ≤ *M* (*T* ′, *σ*′, *ℓ*′) ≤ |*E*(*R*)|, since for each clonal tree edge (*u, v*) ∈ *E*(*T*), there exists a refinement graph edge (*s, t*) labeled by *e*((*s, t*)) = (*u, v*) by Definition 9. To show that |*E*(*R*)| ≤ |*E*(*T*)| + (|*V* (*T*)| × |Σ|), note that for any clonal tree node *v* ∈ *V* (*T*), *R*_*v*_ is a tree without any multi-edges by Definition 9. Since *V* (*G*) = Σ, *R*_*v*_ can at most have |Σ| nodes, and so there can be at most |Σ| − 1 edges labeled with *v* for any *v* ∈ *V* (*T*). So in *R*, there are |*E*(*T*)| refinement tree edges labeled with a clonal tree edge and at most |*V* (*T*)|(|Σ|−1) edges labeled with a clonal tree node. Therefore, |*E*(*R*)| ≤ |*E*(*T*)| + |*V* (*T*)|(|Σ| − 1).

**(Main text) Proposition 9**. *If* M *is the first criterion in* ℱ, *then for any node u* ∈ *V* (*T*) *without polytomies, i*. *e*. |*δ*_*T*_ (*u*)| *<* 2, *it holds that* 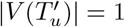.

*Proof*. Assume for contradiction that 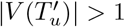 for a node *u* observed in less than two locations. Therefore by the pigeonhole principle, at least one node *u*′ in 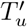 labeled by *ℓ*′(*u*′) = *s* is is an unobserved clone, i. e 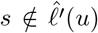. As 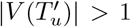 implies there is at least one edge in connected subtree 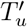, without loss of generality, assume (*u*′, *u*″) is an edge in 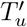 . We construct a new refinement (*T* ″, *σ*″, *ℓ*″) by (i) deleting *u*″, (ii) attaching the children of *u*^″^ as the children of *u*′, and (iii) assigning the location of *u*″ to be the location of *u*′, i. e. *ℓ*″(*u*′) = *ℓ*′(*u*″). Clearly, *U* (*T* ″, *σ*″, *ℓ*″) *< U* (*T* ′, *σ*′, *ℓ*′), which contradicts with the fact that (*T* ″, *σ*″, *ℓ*″) is optimal when M is the first criterion in ℱ.

**(Main text) Proposition 10**. *Let* (*T* ′, *σ*′, *ℓ*′) *be a refinement that minimizes the number of unobserved clones. Then there exists an optimal refinement* (*T* ′, *σ*′, *ℓ*′) *such that for any node u* ∈ *V* (*T*) *without polytomies, i*. *e*. |*δ*_*T*_ (*u*)| *<* 2 *and observed in less than two locations, i*. *e*. 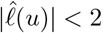, *it holds that* 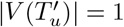.

*Proof*. Assume (*T* ′, *σ*′, *ℓ*′) is an optimal refinement where for a node *u* ∈ *V* (*T*) without polytomies, i. e. |*δ*_*T*_ (*u*)| *<* 2, it holds that 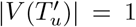. Assume (*u*′, *u*″) is an edge in *T*_*u*_ such that *u*″ is an edge in 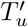. Note that (*u*′, *u*″) is a migration by condition (ii) of Definition 2. Now we construct a new refinement (*T* ″, *σ*″, *ℓ*″) with 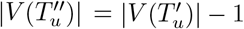 without increasing the number of migrations. We consider three cases.

1. *u*″ does not have any children. In this case, deleting migration (*u*′, *u*″) decreases the number of migrations, contradicting the premise.
2. *u*″ has one child *v*′. In this case, assigning the label of *u*″ to *v*′, i. e. *ℓ*″(*v*′) = *ℓ*′(*u*″) and then collapsing edge (*u*′, *u*″) reduces the number of migration by one (if (*u*″, *v*′) is a migration), or keeps the migration number the same (if (*u*″, *v*′) is not a migration).
3. *u*″ has two children *v*′ and *w*′. Since *u* has at most two children, *u*′ cannot have any outgoing edge outside of 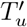 . Thus deleting migration (*u*′, *u*″) decreases the number of migrations.

Thus we can get a refinement (*T* ″, *σ*″, *ℓ*″) that aligns with the premise by repeatedly applying the abovementioned steps.

### B. 4 Complexity

In this section, we study the complexity of PMH-TR for different criteria ordering ℱ. To that end, we consider the natural decision and counting versions of the PMH-TR problem, which itself was posed as an optimization problem. Fig. S2c summarizes all the prior and new complexity results. We start with the decision problem.

#### Problem 2

(Decision Parsimonious Migration History with Tree Refinement (Dec-PMH-TR)). *Given a clonal tree T*, *set* Σ *of locations, observed labeling* 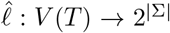, *criteria ordering* ℱ = {*f*_1_, …, *f*_*k*_} *and cost vector* (*c*_1_, …, *c*_*k*_) ∈ ℕ, *does there exist a refinement* (*T* ′, *σ*′, *ℓ*′) *of* (*T*, 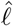) *with cost* (*f*_1_(*T* ′, *σ*′, *ℓ*′), …, *f*_*k*_(*T* ′, *σ*′, *ℓ*′)) *at most* (*c*_1_, …, *c*_*k*_).

**Figure S12:**
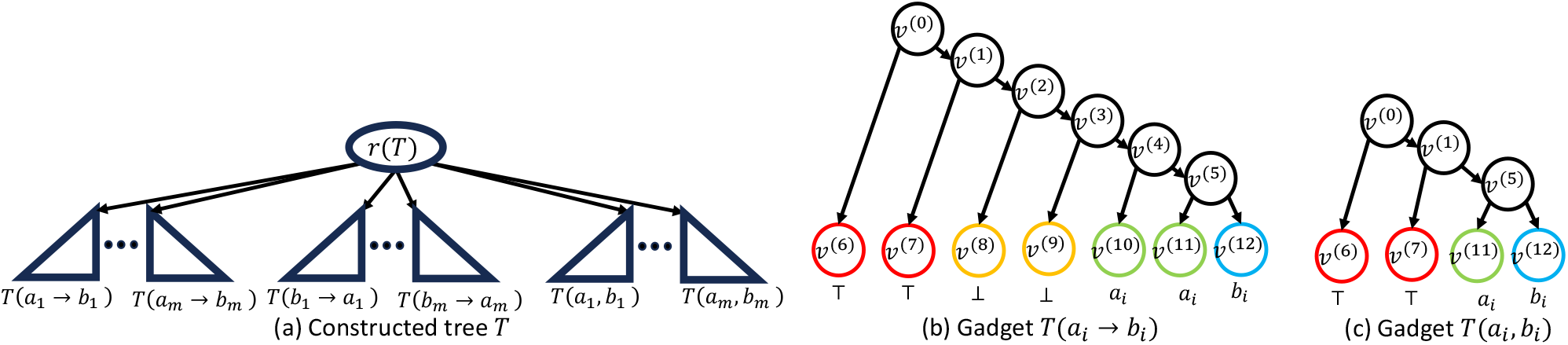
Reduction from VERTEX COVER. (a) For each undirected edge (*ai, bi*) in graph 𝒢, the constructed clonal tree *T* contains three gadgets *T* (*ai* → *bi*), *T* (*bi* → *ai*), and *T* (*ai, bi*). (b) and (c) illustrates the gadgets *T* (*ai* → *bi*) and *T* (*ai, bi*) with observed labeling 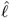, respectively. Each leaf *v* is observed in exactly one location, i. e. 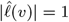. Internal nodes *v* are not observed anywhere, i. e. 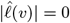.

We also consider the counting problem defined as follows.

#### Problem 3

(Counting Parsimonious Migration History with Tree Refinement (#PMH-TR)). *Given a clonal tree T*, *set* Σ *of locations, observed labeling* 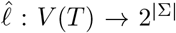 *and criteria ordering* F, *count the number of refinements* (*T* ′, *σ*′, *ℓ*′) *of* (*T*, 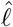) *that are lexicographically optimal with respect to* ℱ.

#### B. 4. 1 Hardness of the PMH-TR Decision Problem

We prove that Dec-PMH-TR is NP-complete when ℱ= MC, UMC, MUC, MCU by reduction from Vertex Cover, an NP-complete problem [46]. In an undirected graph 𝒢 with node set *V* (𝒢) and edge set *E*(𝒢), a *vertex cover K* is a subset of nodes *V* (𝒢) such that for each edge (*a*_*i*_, *b*_*i*_) *E*(), at least one of *a*_*i*_ or *b*_*i*_ is in *K*. Given 𝒢 and an integer *k*, the Vertex Cover problem asks if there is vertex cover *K*^*^ of size *k*.

##### Theorem 16.

Dec-PMH-TR *is NP-complete for* ℱ ∈ {MC, UMC, MUC, MCU}.

We do this by giving a polynomial-time reduction from Vertex Cover to Dec-PMH-TR. More specifically, given an undirected graph 𝒢 with |*V* (𝒢)| = *n* nodes and |*E*(𝒢)| = *m* edges, we construct a Dec-PMH-TR instance consisting of a clonal tree *T* with locations Σ = *V* (𝒢) ∪ {⊥}, and observed labeling 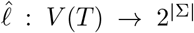. The construction ensures that (i) the optimal refinement (*T* ^*^, *σ*^*^, *ℓ*^*^) admits *U* (*T* ^*^, *σ*^*^, *ℓ*^*^) = 15*m* unobserved clones, *M* (*T* ^*^, *σ*^*^, *ℓ*^*^) = 8*m* migrations and *C*(*T* ^*^, *σ*^*^, *ℓ*^*^) = 2*m* + *n* + 1 + |*K*^*^| comigrations, where *K*^*^ is the cardinality of a minimum vertex cover *K*^*^ for 𝒢, and (ii) there is a comigration from to *a* if node *a ∈ V* (𝒢) is in *K*^*^. We satisfy these requirements by using two types of gadgets that form subtrees of *T*, as illustrated in Fig. S12a.

i. Gadget *T* (*a*_*i*_ → *b*_*i*_). For each undirected edge (*a*_*i*_, *b*_*i*_) in G, we attach two gadgets *T* (*a*_*i*_ → *b*_*i*_) and *T* (*b*_*i*_ → *a*_*i*_) with {⊤, ⊥, *a*_*i*_, *b*_*i*_} distinct labels to *r*(*T*) in *T* (Fig. S12b). There are 2*m* such gadgets.
ii. Gadget *T* (*a*_*i*_, *b*_*i*_). For each undirected edge (*a*_*i*_, *b*_*i*_) in 𝒢, we attach one gadget *T* (⊤, ⊥, *a*_*i*_, *b*_*i*_) with,, *a*_*i*_, *b*_*i*_ distinct labels to *r*(*T*) in *T* (Fig. S12c). There are *m* such gadgets.

Note that in constructed tree *T*, all the nodes *v*^(*k*)^ are observed in either zero or one location, i. e. 0 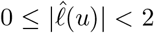. Therefore, for ℱ= UMC, it holds that 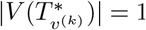 for each node *v*^(*k*)^ by Proposition 9. On the other hand, only the root *r*(*T*) has more than two children. Thus, by Proposition 10, there exists an optimal refinement (*T* ′, *σ*′, *ℓ*′) such that for each non-root node *v*^(*k*)^, 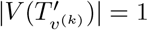 when ℱ ∈ {MC, MCU, MUC}, and in this case, we assume (*T* ^*^, *σ*^*^, *ℓ*^*^) to be such a refinement. Let *v*^*′*(*k*)^ be the unique node mapping to the original non-root node *σ*^*^(*v*^*′*(*k*)^) = *v*^(*k*)^. By applying Lemma 7 to the gadgets of *T* ′, we prove that (*T* ^*^, *σ*^*^, *ℓ*^*^) has 8*m* migrations and determine the location label of all the nodes in gadgets *T* (*a*_*i*_ → *b*_*i*_) and the nodes *v*^*′*(0)^ and *v*^*′*(1)^ of gadgets *T* (*a*_*i*_, *b*_*i*_).

##### Lemma 17.

*For any optimal refinement* (*T*^*^, *σ*^*^, *ℓ*^*^) *for criteria ordering* ℱ ∈ {MC, UMC, MUC, MCU}, *it holds that*

i. *U* (*T* ^*^, *σ*^*^, *ℓ*^*^) = 15*m*,
ii. *M* (*T* ^*^, *σ*^*^, *ℓ*^*^) = 8*m*,
iii. *for each node v*′ *in a gadget, there is a leaf w*′ *in the same gadget such as ℓ*^*^(*v*′) = *ℓ*′ (*w*′),
iv. *ℓ*^*^(*v*′ ^(0)^) = *ℓ*^*^(*v*′ ^(1)^) = ⊤ *for every gadget*,
v. *ℓ*^*^(*v*′ ^(2)^) = *ℓ*^*^(*v*′ ^(23)^) = ⊥ *for gadgets T* (*a*_*i*_ → *b*_*i*_),
vi. 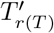 *contains exactly one node r*(*T* ′) *and ℓ*^*^(*r*(*T* ′)) = ⊤.

*Proof*. To prove (i)-(ii), first we prove that *U* (*T*^*^, *σ*^*^, *ℓ*^*^) ≥ 15*m* and *M* (*T* ^*^, *σ*^*^, *ℓ*^*^) ≥ 8*m*. Note that *U* (*T*^*^, *σ*^*^, *ℓ*^*^) ≥ 15*m* directly follows from applying Lemma 5 to (*T* ^*^, *σ*^*^, *ℓ*^*^). To prove *M* (*T* ^*^, *σ*^*^, *ℓ*^*^) ≥ 8*m*, we apply Lemma 6 to each gadget and find the minimum number of migrations in *T* ^*^.

- Each gadget *T* ^*^(*a*_*i*_ → *b*_*i*_) has 4 distinct leaf labels {⊤, ⊥, *a*_*i*_, *b*_*i*_}, and so has a minimum of 4−1 = 3 migrations. Thus, 2*m* gadgets *T* ^*^(*a*_*i*_ → *b*_*i*_) contribute at least 3 · 2*m* = 6*m* migrations in *T* ^*^.
- Each gadget *T* ^*^(*a*_*i*_, *b*_*i*_) has 3 distinct leaf labels, {⊤, *a*_*i*_, *b*_*i*_ }, and so has a minimum of 3 − 1 = 2 migrations. Thus, *m* gadgets *T* ^*^(*a*_*i*_, *b*_*i*_) contribute at least 2*m* migrations in *T* ^*^.

Totalling, *M* (*T* ^*^, *σ*^*^, *ℓ*^*^) ≥ 8*m*.

Now to prove that the optimal refinement (*T* ^*^, *σ*^*^, *ℓ*^*^) for ℱ ∈ {MC, UMC, MUC, MCU} has exactly 15*m* unobserved clones and 8*m* migrations, it suffices to show that there exists at least one refinement (*T* ′, *σ*′, *ℓ*′) that simultaneously reaches the lower limit for unobserved clones *U* (*T* ^*^, *σ*^*^, *ℓ*^*^) = 15*m* and the number of migrations *M* (*T* ′, *σ*′, *ℓ*′) = 8*m*. We construct such a (*T* ′, *σ*′, *ℓ*′) as follows.

- set *ℓ*′ (*v*′ ^(0)^) = *ℓ*′ (*v*′ ^(1)^) = ⊤ for every gadget,
- set *ℓ*′ (*v*′ ^(2)^) = *ℓ*′ (*v*′ ^(3)^) = ⊥ and *ℓ*′ (*v*′ ^(4)^) = *ℓ*′ (*v*′ ^(5)^) = *a*_*i*_ for gadgets *T* (*a*_*i*_ → *b*_*i*_).
- set *ℓ*′ (*v*′^(5)^) = *a*_*i*_ for gadgets *T* (*a*_*i*_, *b*_*i*_).

Clearly (*T* ′, *σ*′, *ℓ*′) has 15*m* unobserved clones and 8*m* migrations. Therefore, *U* (*T* ^*^, *σ*^*^, *ℓ*^*^) = 15*m* and *M* (*T* ^*^, *σ*^*^, *ℓ*^*^) = 8*m*.

To prove (iii)-(v), notice that (*T* ^*^, *σ*^*^, *ℓ*^*^) achieves the minimum number 8*m* of migrations if each gadget has the minimum number of migrations. As such, by Lemma 7, for each node *v*′ in a gadget, there is a leaf *w*′ in the same gadget such as *ℓ*^*^(*v*′) = *ℓ*′(*w*′). Moreover, if two leaves *w*′ and *w*″ in a gadget in *T* ′ have the same location label, i. e. *ℓ*^*^(*w*′) = *ℓ*^*^(*w*″), then each node *v*′ in the (undirected) path from *w*′ to *w*″ has the same label *ℓ*^*^(*v*′) = *ℓ*^*^(*w*′). Now in every gadget, *ℓ*^*^(*v*′^(6)^) = *ℓ*^*^(*v*′^(7)^) = ⊤, and the path between them contains *v*′^(0)^ and *v*′^(1)^. Thus, *ℓ*′(*v*′^(0)^) = *ℓ*′(*v*′^(1)^) = ⊤. Similarly, *ℓ*′(*v*′^(2)^) = *ℓ*′(*v*′^(3)^) = ⊥ and *ℓ*′(*v*′^(4)^) = *ℓ*′(*v*′^(5)^) = *a*_*i*_ for gadgets *T* (*a*_*i*_ → *b*_*i*_), since *v*′^(2)^ and *v*′^(3)^ are in the path between *v*′^(8)^ and *v*′^(9)^ where *ℓ*′(*v*′^(8)^) = *ℓ*′(*v*′^(9)^) = ⊥, and *v*′^(4)^ and *v*′^(5)^ are in the path between *v*′^(10)^ and *v*′^(11)^ where *ℓ*′(*v*′^(10)^) = *ℓ*′(*v*′^(11)^) = *a*_*i*_.

We finish proving that the lemma is true for ℱ ∈ {MC, MCU, MUC, UMC} by showing that 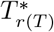 contains exactly one node *r*(*T* ^*^) and *ℓ*^*^(*r*(*T* ^*^)) = ⊤. As 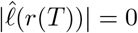, by Proposition 9, 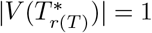 for ℱ = UMC. For ℱ ∈ {MC, MCU, MUC}, notice that *M* (*T* ^*^, *σ*^*^, *ℓ*^*^) = 8*m* implies every migration is inside a gadget, every gadget has the minimum number of migrations, and the root *v*′^(0)^ of each gadget is labeled by ⊤. Now if there is a node *v*′ such that *σ*^*^(*v*′) = *r*(*T*) and *ℓ*^*^(*v*′) ≠ ⊤, then there is at least one migration in the path from *v*′ to the root *v*′^(0)^ of any gadget in *T* ^*^. But that puts migration outside the gadgets, which increases the number of migrations, thereby reaching a contradiction. So there is exactly one node *r*(*T* ′) such that *σ*^*^(*r*(*T* ′)) = *r*(*T*), and *ℓ*^*^(*r*(*T* ′)) = ⊤. □

Based on Lemma 17, we can get a list of comigrations that are present in (*T* ^*^, *σ*^*^, *ℓ*^*^).

##### Corollary 18.

*For any optimal refinement* (*T* ^*^, *σ*^*^, *ℓ*^*^) *for criteria ordering* ℱ ∈ {MC, UMC, MUC, MCU }, *there are comigrations from*

- ⊤ *to* ⊥,
- ⊥ *to a*_*i*_ *for each a*_*i*_ ∈ *V* (𝒢),
- *a*_*i*_ *to b*_*i*_ *and b*_*i*_ *to a*_*i*_ *for each* (*a*_*i*_, *b*_*i*_) ∈ *E*(𝒢).

Finally, we prove that the size of the minimum vertex cover for graph 𝒢 can be determined from the number *C*(*T* ^*^, *σ*^*^, *ℓ*^*^) of comigrations.

##### Proposition 19.

*There exists a minimum vertex cover K*^*^ *of size* |*K*^*^| = *k if and only if there exists an optimal refinement* (*T* ^*^, *σ*^*^, *ℓ*^*^) *for criteria ordering* ℱ ∈ {MC, UMC, MUC, MCU} *with U* (*T* ^*^, *σ*^*^, *ℓ*^*^) = 15*m unobserved clones, M* (*T* ^*^, *σ*^*^, *ℓ*^*^) = 8*m migrations and C*(*T* ^*^, *σ*^*^, *ℓ*^*^) = *n* + 2*m* + 1 + *k comigrations*.

*Proof*. (⇒) Assume there is a refinement (*T* ^*^, *σ*^*^, *ℓ*^*^) with *U* (*T* ^*^, *σ*^*^, *ℓ*^*^) = 15*m* unobserved clones, *M* (*T* ^*^, *σ*^*^, *ℓ*^*^) = 8*m* migrations and *C*(*T* ^*^, *σ*^*^, *ℓ*^*^) = *n* + 2*m* + 1 + *k* comigrations. Let us consider the gadget *T* ^*^(*a*_*i*_, *b*_*i*_). *T* ^*^(*a*_*i*_, *b*_*i*_) has distinct leaf labels, {⊤, *a*_*i*_, *b*_*i*_ }, and node *v*′^(1)^ in *T* ^*^(*a*_*i*_, *b*_*i*_) is labeled with *ℓ*^*^(*v*′^(1)^) = ⊤. Now by condition (iii) of Lemma 17, *v*′^(5)^ is labeled with one of ⊤, *a*_*i*_, or *b*_*i*_. If *v*′^(5)^ is labeled with ⊤, then there exists comigrations from ⊤ to both *a*_*i*_ and *b*_*i*_, which is not listed in Corollary 18. Again, if *v*′^(5)^ is labeled with *a*_*i*_ or *b*_*i*_, then there exists comigrations from ⊤ to either *a*_*i*_ or *b*_*i*_ not listed in Corollary 18. Let *K* be the set of nodes from 𝒢 such that *a*_*i*_ is in *K* if there is a migration from ⊤ to *a*_*i*_ in the gadget *T* (*a*_*i*_, *b*_*j*_).

Now for each edge (*a*_*i*_, *b*_*i*_) in 𝒢, *T* ^*^(*a*_*i*_, *b*_*i*_) has at least one comigration from ⊤ to either *a*_*i*_ or *b*_*i*_ or both, which implies that at least one of *a*_*i*_ or *b*_*i*_ is in *K*. Thus *K* is a vertex cover for 𝒢. Again, Corollary 18 lists *n* + 2*m* + 1 comigrations present in (*T* ^*^, *σ*^*^, *ℓ*^*^). Thus, (*T* ^*^, *σ*^*^, *ℓ*^*^) has (*n* + 2*m* + 1 + *k*) − (*n* + 2*m* + 1) = *k* comigrations not listed in Corollary 18. Since all the comigrations in (*T* ^*^, *σ*^*^, *ℓ*^*^) are either listed in Corollary 18 or from to an element from *K*, |*K*| = *k*.

(⇐) Assume that *K*^*^ is a minimum vertex cover for the undirected graph 𝒢. We show that there exists an optimal refinement with *U* (*T* ^*^, *σ*^*^, *ℓ*^*^) = 15*m* unobserved clones, *M* (*T* ^*^, *σ*^*^, *ℓ*^*^) = 8*m* migrations and *C*(*T* ^*^, *σ*^*^, *ℓ*^*^) = *n* + 2*m* + 1 + *k* comigrations. We construct one such (*T* ^*^, *σ*^*^, *ℓ*^*^) as follows.

1. Let *T* ^*^ have the same topology as *T* .
2. Set the mapping function *σ*^*^ such that *σ*^*^ induces a bijection between the nodes of *T* and *T* ^*^.
3. Following Lemma 17, set *ℓ*^*^(*r*(*T* ′)) = *ℓ*^*^(*v*′^(0)^) = *ℓ*^*^(*v*′^(1)^) = ⊤ for every gadget, and set *ℓ*^*^(*v*′^(2)^) = *ℓ*^*^(*v*′^(3)^) = ⊥ and *ℓ*^*^(*v*′^(4)^) = *ℓ*^*^(*v*′^(5)^) = *a*_*i*_ for gadgets *T* ^*^(*a*_*i*_ → *b*_*i*_).
4. For every edge (*a*_*i*_, *b*_*i*_) from 𝒢 where *a*_*i*_ is in vertex cover *K*^*^, set *ℓ*^*^(*v*′^(5)^) = *a*_*i*_ in gadget *T* ^*^(*a*_*i*_, *b*_*i*_).

Clearly, (*T* ^*^, *σ*^*^, *ℓ*^*^) is a refinement with *U* (*T* ^*^, *σ*^*^, *ℓ*^*^) = 15*m* unobserved clones, *M* (*T* ^*^, *σ*^*^, *ℓ*^*^) = 8*m* migrations and *C*(*T* ^*^, *σ*^*^, *ℓ*^*^) = *n* + 2*m* + 1 + *k* comigrations. □

Finally, we prove our second theorem about NP-completeness for criteria orderings {CM, UCM, CMU}.

##### Theorem 20.

Dec-PMH-TR *is NP-complete for* ℱ ∈ {CM, CMU, UCM}.

*Proof*. We show the hardness of PMH-TR separately for each criteria ordering {CM, CMU, UCM}.

1. *F* = CM. PMH-TR is proven to be NP-complete for *F* = CM in El-Kebir et al. [41].
2. *F* = CMU. Notice that PMH-TR solution space for *F* = CMU is a subset of the solution space for *F* = CM. Therefore, PMH-TR for *F* = CM can be trivially reduced to PMH-TR for *F* = CMU, as if there is a refinement with at most *c* comigrations and *m* migrations, then there is a refinement with at most *c* comigrations, *m* migrations, and 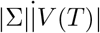 unobserved clones.
3. *F* = UCM. This problem can be reduced from PMH for *F* = CM, proven to be NP-complete in El-Kebir et al. [41]. In PMH for *F* = CM, given a clonal tree *T* with leaf labeling 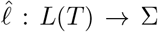, and two integers *c* and *m*, one asks if there is a location labeling *ℓ*′ that minimizes the number of comigrations first, and then migrations. In other words, given 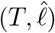 and two integers *c* and *m*, PMH seeks a refinement (*T* ′, *σ*′, *ℓ*′) such that 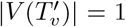 for each node *v, C*(*T* ′, *σ*′, *ℓ*′) = *c*, and *M* (*T* ′, *σ*′, *ℓ*′) = *m*. As 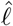 labels clones with less than two locations, the constraint 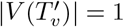 can be enforced by requiring minimizing the number of unobserved clones first, as suggested by Proposition 5. Therefore, the solution space for PMH instance 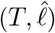 for criteria ordering *F* = CM and the solution space for PMH-TR instance 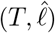 for criteria ordering *F* = UCM are identical. Thus, PMH-TR is NP-complete for *F* = UCM.

**Figure S13:**
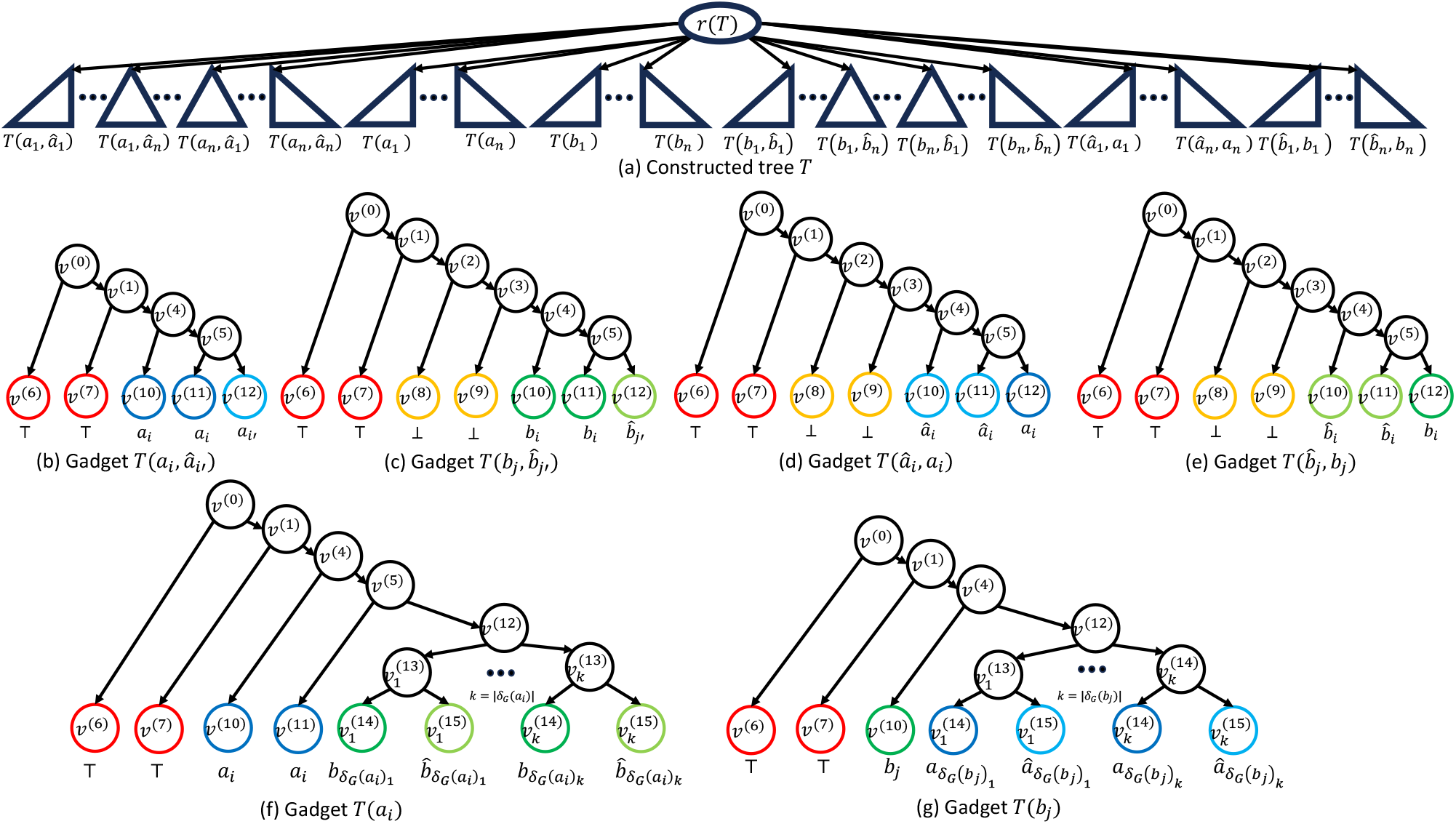
Construction of the clonal tree *T* given a bipartite graph 𝒢 with *V* (𝒢) = *A* ∪ *B* nodes. (a) The constructed tree *T* containing six types of gadget is shown. (b-g) illustrates six types of gadgets.

#### B. 4. 2 Hardness of the #PMH-TR Counting Problems

We prove that PMH-TR is #P-complete when ℱ ∈ {UMC, MUC, MCU} by providing a strong parsimony reduction from the problem of counting the number of perfect matchings in a bipartite graph, which itself is #P-complete [47]. A graph 𝒢 with nodes *V* (𝒢) and edges *E*(𝒢) is bipartite if its nodes *V* (𝒢) can be partitioned into two disjoint sets *A* and *B* such that each edge (*a*_*i*_, *b*_*j*_) *E* ∈ (𝒢) connects a node *a*_*i*_ from *A* to a node *b*_*j*_ from *B*. A perfect matching *M* in a bipartite graph 𝒢 is a subset of edges *E*(𝒢) such that each node *a*_*i*_ or *b*_*j*_ in *V* (𝒢) = *A ∪ B* is incident to exactly one edge (*a*_*i*_, *b*_*j*_) in *M* . Though determining if there is a perfect matching in a given bipartite graph is polynomial-time solvable, counting the number of perfect matchings is #P-complete [47].

##### Theorem 21.

#PMH-TR *is #P-complete for* ℱ ∈ {UMC, MUC, MCU}.

Clearly, #PMH-TR is in #P as the underlying decision problem (where one additionally specifies the set of comigrations as a partition of migrations) is in NP, i. e. one can verify correctness of the solution in polynomial time. To prove #P-hardness, we show a polynomial-time parsimonious reduction from perfect matching in a bipartite graph to PMH-TR for criteria ordering ℱ ∈ {UMC, MUC, MCU}. To that end, consider a bipartite graph 𝒢 with *V* (𝒢) = *A* ∪ *B* nodes and *E*(𝒢) edges, where |*A*| = |*B*| = *n*, |*E*(𝒢)| = *m, A* = {*a*_1_, …, *a*_*n*_} and *B* = {*b*_1_, …, *b*_*n*_}. Let the function *δ*_𝒢_ : *V* (𝒢) → 2^|*V* (𝒢)|^ indicate node adjacency in 𝒢 such that *δ*_𝒢_ (*a*_*i*_) = *b*_*j*_ and *δ*_𝒢_ (*b*_*j*_) = *a*_*i*_ if (*a*_*i*_, *b*_*j*_) is in 𝒢. In addition, we declare new symbols 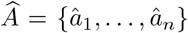 and 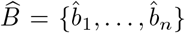. Given the bipartite graph 𝒢, we construct a clonal tree *T* with leaf labeling 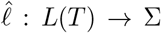 that labels the leaves of *T* with locations 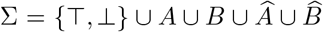 such that the comigrations induced by each optimal refinement (*T* ^*^, *σ*^*^, *ℓ*^*^) encodes a perfect matching *K* of 𝒢. Specifically, we want to ensure that (i) for each optimal refinement (*T* ^*^, *σ*^*^, *ℓ*^*^) there are exactly 10*n*^2^ + 21*n* + 2*m* unobserved clones, 5*n*^2^ + 8*n* + 4*m* migrations and at least 2*n*^2^ + 6*n* + 1 comigrations, and (ii) there is a perfect matching *K* if (*T* ^*^, *σ*^*^, *ℓ*^*^) has exactly 2*n*^2^ + 6*n* + 1 comigrations and edge (*a*_*i*_, *b*_*j*_) is in *K* if there is a comigration from *a*_*i*_ to *b*_*j*_. We accomplish this by using six different types of gadget that for the subtrees of *T*, as shown in Fig. S13.

i. Gadget *T* (*a*_*i*_, *â*_*i*_*′*). For each 1 ≤ *i, i*^*′*^ ≤ *n*, we attach one gadget *T* (*a*_*i*_, *â*_*i*_*′*) with {⊤, *a*_*i*_, *â*_*i*_*′*} distinct labels to the root *r*(*T*) (Fig. S13b). There are *n*^2^ such gadgets.
ii. Gadget 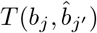. For each 1 ≤ *j, j*^′^≤ *n*, we attach one gadget 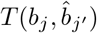 with 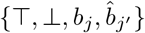 distinct labels to the root *r*(*T*) (Fig. S13c). There are *n*^2^ such gadgets.
iii. Gadget *T* (*â*_*i*_, *a*_*i*_). For each 1 ≤ *i* ≤ *n*, we attach one gadget 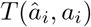 with 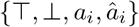 distinct labels to the root *r*(*T*) (Fig. S13d). There are *n* such gadgets.
iv. Gadget 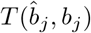. For each 1 ≤ *j* ≤ *n*, we attach one gadget 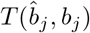 with 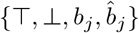 distinct labels to the root *r*(*T*) (Fig. S13e). There are *n* such gadgets.
v. Gadget *T* (*a*_*i*_). For each 1 ≤ *i* ≤ *n*, we attach one gadget *T* (*a*_*i*_) with 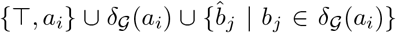 distinct labels to the root *r*(*T*) (Fig. S13f). There are *n* such gadgets.
vi. Gadget *T* (*b*_*j*_). For each 1 ≤ *j* ≤ *n*, we attach one gadget *T* (*b*_*j*_) with 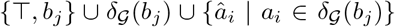 distinct labels to the root *r*(*T*) (Fig. S13g). There are *n* such gadgets.

Note that all the nodes *v*^(*k*)^ of *T* are observed in either zero or one location, i. e. 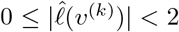. Therefore, for ℱ = UMC, it holds that 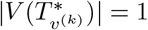 for each node *v*^(*k*)^ (Proposition 9). Moreover, in constructed tree *T*, only the root *r*(*T*) and *v*^(12)^ in gadgets *T* (*a*_*i*_) and *T* (*b*_*j*_) have more than two children. Thus, by Proposition 10, there exists an optimal refinement (*T*^′^, *σ*^*′*^, *ℓ*^*′*^) such that for each node *v*^(*k*)^ where *v*^(*k*)^ *r*(*T*) and 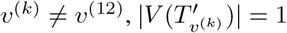 when ℱ ∈ {MCU, MUC}, and in this case, we assume (*T* ^*∗*^, *σ*^*∗*^, *ℓ*^*∗*^) to be such a refinement. Additionally, for each node *v*^(*k*)^ where *v*^(*k*)^ ≠ *r*(*T*) and *v*^(*k*)^ *v*^(12)^, we let *v*^*′*(*k*)^ be the unique node mapping to the original node *σ*^*∗*^(*v*^*′*(*k*)^) = *v*^(*k*)^. By applying Lemma 5 and Lemma 7 to the gadgets of *T*^′^, we prove that (*T* ^*∗*^, *σ*^*∗*^, *ℓ*^*∗*^) has 10*n*^2^ + 21*n* + 2*m* unobserved clones and 5*n*^2^ +8*n*+4*m* migrations, and determine the location label of all the nodes in gadgets 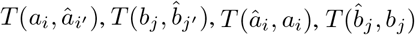 the nodes *v*^*′*(0)^, *v*^*′*(1)^, *v*^*′*(2)^, *v*^*′*(3)^ in gadgets *T* (*a*_*i*_), and the nodes *v*^*′*(0)^, *v*^*′*(1)^ in gadgets *T* (*b*_*j*_).

##### Lemma 22.

*For any optimal refinement* (*T* ^*∗*^, *σ*^*∗*^, *ℓ*^*∗*^) *for criteria ordering* ℱ ∈ {UMC, MUC, MCU}, *it holds that*

i. *U* (*T* ^*∗*^, *σ*^*∗*^, *ℓ*^*∗*^) = 10*n*^2^ + 21*n* + 2*m*,
ii. *M* (*T* ^*∗*^, *σ*^*∗*^, *ℓ*^*∗*^) = 5*n*^2^ + 8*n* + 4*m*,
iii. *for each node u*^*′*^ *in a gadget, if there exists a leaf w*^*′*^ *in the same gadget where *ℓ**^*∗*^(*u*^*′*^) = *ℓ*^*∗*^(*w*^*′*^), *then the nodes v*^*′*^ *in the path from u*^*′*^ *to w*^*′*^ *are labeled with *ℓ**^*∗*^(*v*^*′*^) = *ℓ*^*∗*^(*u*^*′*^),
iv. *ℓ*^*∗*^(*v*^*′*(0)^) = *ℓ*^*∗*^(*v*^*′*(1)^) = ⊤ *and *ℓ**^*∗*^(*v*^*′*(2)^) = *ℓ*^*∗*^(*v*^*′*(3)^) = ⊥ *for every gadget*,
v. *both *ℓ**^*∗*^(*v*^*′*(4)^) *and *ℓ**^*∗*^(*v*^*′*(5)^) *equals* 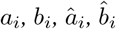 *and a*_*i*_ *for gadgets* 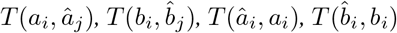 *and T* (*a*_*i*_), *respectively*,
vi. 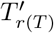 *contains exactly one node r*(*T*^′^) *and *ℓ**^*∗*^(*r*(*T*^′^)) = ⊤.

*Proof*. First we prove that *U* (*T* ^*∗*^, *σ*^*∗*^, *ℓ*^*∗*^) ≥ 10*n*^2^ + 21*n* + 2*m* and *M* (*T* ^*∗*^, *σ*^*∗*^, *ℓ*^*∗*^) ≥ 5*n*^2^ + 8*n* + 4*m*. To that end, we apply Lemma 5 and Lemma 7 to each gadget and find the minimum number of migrations in *T* ^*∗*^.

- Each gadget *T* ^*∗*^(*a*_*i*_, *â*_*i*_*′*) has 4 clones not observed anywhere, and so has a minimum of 4 unobserved clones. Moreover, *T* ^*∗*^(*a*_*i*_, *â*_*i*_*′*) has 3 distinct leaf labels {⊤, *a*_*i*_, *â*_*i*_*′*, and so has a minimum of 3 − 1 = 2 migrations. *n*^2^ gadgets *T* ^*∗*^(*a*_*i*_, *â*_*i*_*′*) contribute 4*n*^2^ unobserved clones and 2*n*^2^ migrations in *T* ^*∗*^.
- Each gadget 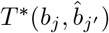 has 6 clones not observed anywhere, and so has a minimum of 6 unobserved clones. Moreover, 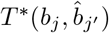 has 4 distinct leaf labels 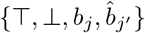, and so has a minimum of 4 − 1 = 3 migrations. *n*^2^ gadgets 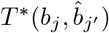 contribute 6*n*^2^ unobserved clones and 3*n*^2^ migrations in *T* ^*∗*^.
- Each gadget *T* ^*∗*^(*â*_*i*_, *a*_*i*_) has 6 clones not observed anywhere, and so has a minimum of 6 unobserved clones. Moreover, *T* ^*∗*^(*â*_*i*_, *a*_*i*_) has 4 distinct leaf labels {⊤, ⊥, *a*_*i*_, *â*_*i*_, and so has a minimum of 4 − 1 = 3 migrations. *n* gadgets *T* ^*∗*^(*â*_*i*_, *a*_*i*_) contribute 6*n* unobserved clones and 3*n* migrations in *T* ^*∗*^.
- Each gadget 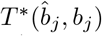 has 6 clones not observed anywhere, and so has a minimum of 6 unobserved clones. Moreover, 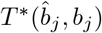 has 4 distinct leaf labels 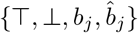 and so has a minimum of 4 − 1 = 3 migrations. *n* gadgets 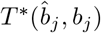 contribute 6*n* unobserved clones and 3*n* migrations in *T* ^*∗*^.
- Each gadget *T* ^*∗*^(*a*_*i*_) has 5 + |*δ*_*G*_(*a*_*i*_)| clones not observed anywhere, and so has a minimum of 5 + |*δ*_*G*_(*a*_*i*_)| unobserved clones. Moreover, *T* ^*∗*^(*a*_*i*_) has 2+2|*δ*_*G*_(*a*_*i*_)| distinct leaf labels 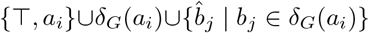 and so has a minimum of 2 + 2|*δ*_*G*_(*a*_*i*_)| − 1 = 2|*δ*_*G*_(*a*_*i*_:)| + 1 migrations. Thus, gadgets *T* ^*∗*^(*a*_*i*_) contribute 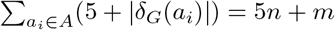 unobserved clones and 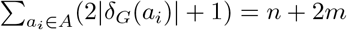 migrations.
- Each gadget *T* ^*∗*^(*b*_*j*_) has 4 + |*δ*_*G*_(*b*_*j*_)| clones not observed anywhere, and so has a minimum of 4 + |*δ*_*G*_(*b*_*j*_)| unobserved clones. Moreover, *T* ^*∗*^(*b*_*j*_) has 2+2|*δ*_*G*_(*b*_*j*_)| distinct leaf labels {⊤, *b*_*j*_}∪*δ*_*G*_(*b*_*j*_)∪{*â*_*i*_ | *a*_*i*_ ∈ *δ*_*G*_(*b*_*j*_)}, and so has a minimum of 2 + 2|*δ*_*G*_(*b*_*j*_)| − 1 = 2|*δ*_*G*_(*b*_*j*_I:)| + 1 migrations. Thus, gadgets *T* ^*∗*^(*b*_*j*_) contribute 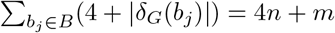 unobserved clones and 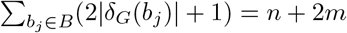 migrations.

Totalling, *U* (*T* ^*∗*^, *σ*^*∗*^, *ℓ*^*∗*^) ≥ 10*n*^2^ + 21*n* + 2*m* and *M* (*T* ^*∗*^, *σ*^*∗*^, *ℓ*^*∗*^) ≥ 5*n*^2^ + 8*n* + 4*m*.

Now to prove that the optimal refinement (*T* ^*∗*^, *σ*^*∗*^, *ℓ*^*∗*^) has exactly 10*n*^2^ + 21*n* + 2*m* unobserved clones and 5*n*^2^ + 8*n* + 4*m* migrations, it suffices to show that there exists at least one PMH-TR refinement (*T*^′^, *σ*^*′*^, *ℓ*^*′*^) such that *U* (*T*^′^, *σ*^*′*^, *ℓ*^*′*^) = 10*n*^2^ + 21*n* + 2*m* and *M* (*T*^′^, *σ*^*′*^, *ℓ*^*′*^) = 5*n*^2^ + 8*n* + 4*m*. We construct such a (*T*^′^, *σ*^*′*^, *ℓ*^*′*^).

- set *ℓ*^*′*^(*v*^*′*(0)^) = *ℓ*^*′*^(*v*^*′*(1)^) = ⊤ and *ℓ*^*′*^(*v*^*′*(2)^) = *ℓ*^*′*^(*v*^*′*(3)^) = ⊥ for every gadget,
- set both *ℓ*^*′*^(*v*^*′*(4)^) and *ℓ*^*′*^(*v*^*′*(5)^) to be 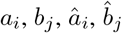 and *a*_*i*_ for gadgets 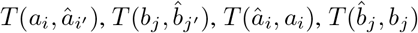 and *T* (*a*_*i*_), respectively,
- let 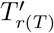 contain exactly one node *r*(*T*^′^) and set *ℓ*^*′*^(*r*(*T*^′^)) = ⊤,
- in gadgets *T*^′^(*a*_*i*_), set *ℓ*^*′*^(*v*^(12)^) and 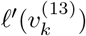 for any 1 ≤ *k* ≤ |*δ*_*G*_(*a*_*i*_)| to be *a*_*i*_, and in gadgets *T*^′^(*b*_*j*_), set 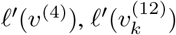 for any 1 ≤ *k* ≤ |*δ*_*G*_(*b*_*j*_)|, and *ℓ*^*′*^(*v*^(13)^) to be *b*_*j*_.

Clearly *T*^′^ has 10*n*^2^ + 21*n* + 2*m* unobserved clones and 5*n*^2^ + 8*n* + 4*m* migrations. Therefore, *U* (*T* ^*∗*^, *σ*^*∗*^, *ℓ*^*∗*^) = 10*n*^2^ + 21*n* + 2*m* and *M* (*T* ^*∗*^, *σ*^*∗*^, *ℓ*^*∗*^) = 5*n*^2^ + 8*n* + 4*m*.

To prove (iii)-(v), notice that (*T* ^*∗*^, *σ*^*∗*^, *ℓ*^*∗*^) achieves the minimum number 5*n*^2^ + 8*n* + 4*m* of migrations if each gadget has the minimum number of migrations. As such, by Lemma 7, for each node *v*^*′*^ in a gadget, there is a leaf *w*^*′*^ in the same gadget such as *ℓ*^*∗*^(*v*^*′*^) = *ℓ*^*′*^(*w*^*′*^). Moreover, if two leaves *w*^*′*^ and *w*^″^ in a gadget in *T*^′^ have the same location label, i. e. *ℓ*^*∗*^(*w*^*′*^) = *ℓ*^*∗*^(*w*^″^), then each node *v*^*′*^ in the (undirected) path from *w*^*′*^ to *w*^″^ has the same label *ℓ*^*∗*^(*v*^*′*^) = *ℓ*^*∗*^(*w*^*′*^). Now in every gadget, *ℓ*^*∗*^(*v*^*′*(6)^ = *ℓ*^*∗*^(*v*^*′*(7)^ = ⊤, and the path between them contains *v*^*′*(0)^ and *v*^*′*(1)^. Thus, *ℓ*^*′*^(*v*^*′*(0)^) = *ℓ*^*′*^(*v*^*′*(1)^) = ⊤. Similarly we can prove that *ℓ*^*′*^(*v*^*′*(2)^) = *ℓ*^*′*^(*v*^*′*(3)^) = ⊥ for every gadget, and both *ℓ*^*∗*^(*v*^*′*(4)^) and *ℓ*^*∗*^(*v*^*′*(5)^) equals 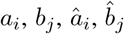 and *a*_*i*_ for gadgets 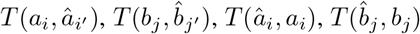 and *T* (*a*_*i*_), respectively.

Last, we prove that 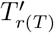 contains exactly one node *r*(*T*^′^) and *ℓ*^∗^(*r*(*T*′)) = ⊤. 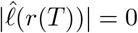 by Proposition 9, 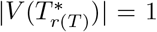 for ℱ = UMC. For ℱ ∈ {MC, MCU, MUC}, notice that |*M* (*T* ^*∗*^, *ℓ*^*∗*^)| = 5*n*^2^ + 8*n* + 4*m* implies every migration is inside a gadget, every gadget has the minimum number of migrations, and the root *v*^*′*(0)^ of each gadget is labeled by . Now if there is a node *v*^*′*^ such that *σ*^*∗*^(*v*^*′*^) = *r*(*T*) and *ℓ*^*∗*^(*v*^*′*^) =, then there is at least one migration in the path from *v*^*′*^ to the root *v*^*′*(0)^ of any gadget in *T* ^*∗*^. But that puts migration outside the gadgets, which increases the number of migrations, thereby reaching a contradiction. So there is exactly one node *r*(*T*^′^) such that *σ*^*∗*^(*r*(*T*^′^)) = *r*(*T*), and *ℓ*^*∗*^(*r*(*T*^′^)) = ⊤.

Based on Lemma 22, we can get a list of comigrations that are present in (*T* ^*∗*^, *σ*^*∗*^, *ℓ*^*∗*^).

##### Corollary 23.

*For any optimal refinement* (*T* ^*∗*^, *σ*^*∗*^, *ℓ*^*∗*^) *for criteria ordering* ℱ ∈{UMC, MUC, MCU}, *there are comigrations from*

- ⊤ *to* ⊥,
- ⊤ *to a*_*i*_ *for any* 1 ≤ *i* ≤ *n*,
- ⊥ *to â*_*i*_ *for any* 1 ≤ *i* ≤ *n*,
- *a*_*i*_ *to â*_*i*_*′ for any* 1 ≤ *i, i*^*′*^ ≤ *n*,
- *â*_*i*_ *to a*_*i*_ *for any* 1 ≤ *i* ≤ *n*,
- ⊥ *to b*_*j*_ *for any* 1 ≤ *j* ≤ *n*,
- ⊥ *to* 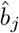 *for any* 1 ≤ *j* ≤ *n*,
- *b*_*j*_ *to* 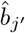 *for any* 1 ≤ *j, j*^*′*^ ≤ *n*,
- 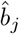 *to b*_*j*_ *for any* 1 ≤ *j* ≤ *n*.

Last, we prove that there is a bijection between perfect matchings *K* and optimal refinements (*T* ^*^, *σ*^*^, *ℓ*^*^) with *U* (*T* ^*^, *σ*^*^, *ℓ*^*^) = 10*n*^2^ + 21*n* + 2*m* unobserved clones, *M* (*T* ^*^, *σ*^*^, *ℓ*^*^) = 5*n*^2^ + 8*n* + 4*m* migrations, and *C*(*T* ^*^, *σ*^*^, *ℓ*^*^) = 2*n*^2^ + 6*n* + 1 comigrations.

##### Proposition 24.

*There is a bijection between the set of perfect matchings K of G and the set of optimal refinements* (*T* ^*^, *σ*^*^, *ℓ*^*^) *with U* (*T* ^*^, *σ*^*^, *ℓ*^*^) = 10*n*^2^ + 21*n* + 2*m unobserved clones, M* (*T* ^*^, *σ*^*^, *ℓ*^*^) = 5*n*^2^ + 8*n* + 4*m migrations, and C*(*T* ^*^, *σ*^*^, *ℓ*^*^) = 2*n*^2^ + 6*n* + 1 *comigrations*.

*Proof*. We prove the lemma in four steps: (i) we show that *C*(*T* ^*^, *σ*^*^, *ℓ*^*^) ≥ 2*n*^2^ + 6*n* + 1 for any optimal refinement (*T* ^*^, *σ*^*^, *ℓ*^*^), which implies that any refinement (*T* ^*^, *σ*^*^, *ℓ*^*^) with *U* (*T* ^*^, *σ*^*^, *ℓ*^*^) = 10*n*^2^ + 21*n* + 2*m* unobserved clones, *M* (*T* ^*^, *σ*^*^, *ℓ*^*^) = 5*n*^2^ + 8*n* + 4*m* migrations, and *C*(*T* ^*^, *σ*^*^, *ℓ*^*^) = 2*n*^2^ + 6*n* + 1 comigrations is optimal, (ii) we prove that for any optimal refinement (*T* ^*^, *σ*^*^, *ℓ*^*^) with *U* (*T* ^*^, *σ*^*^, *ℓ*^*^) = 10*n*^2^ + 21*n* + 2*m* unobserved clones, *M* (*T* ^*^, *σ*^*^, *ℓ*^*^) = 5*n*^2^ +8*n*+4*m* migrations, and *C*(*T* ^*^, *σ*^*^, *ℓ*^*^) = 2*n*^2^ +6*n*+1 comigrations, there is a unique perfect matching *K* such that (*T* ^*^, *σ*^*^, *ℓ*^*^) has a comigration from *a*_*i*_ to *b*_*j*_ for each edge (*a*_*i*_, *b*_*j*_) in *K*, (iii) given a perfect matching *K*, we construct an optimal refinement (*T* ^*^, *σ*^*^, *ℓ*^*^) with *U* (*T* ^*^, *σ*^*^, *ℓ*^*^) = 10*n*^2^ + 21*n* + 2*m* unobserved clones, *M* (*T* ^*^, *σ*^*^, *ℓ*^*^) = 5*n*^2^ + 8*n* + 4*m* migrations, and *C*(*T* ^*^, *σ*^*^, *ℓ*^*^) = 2*n*^2^ + 6*n* + 1 comigrations that admits a comigration from *a*_*i*_ to *b*_*j*_ for each edge (*a*_*i*_, *b*_*j*_) in *K*, and (iv) we prove that the constructed refinement (*T* ^*^, *σ*^*^, *ℓ*^*^) as the only refinement admitting a comigration from *a*_*i*_ to *b*_*j*_ for each edge (*a*_*i*_, *b*_*j*_) in *K*.

First, we prove *C*(*T* ^*^, *σ*^*^, *ℓ*^*^) ≥ 2*n*^2^ +6*n*+1. Note that there are at least 2*n*^2^ +5*n*+1 comigrations in (*T* ^*^, *σ*^*^, *ℓ*^*^) listed in Corollary 23. Let us consider the gadget *T* ^*^(*a*_*i*_). *T* ^*^(*a*_*i*_) has distinct leaf labels 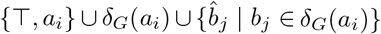 and node *v*^*′*(5)^ in *T* ^*^(*a*_*i*_) is labeled with *ℓ*^*^(*v*^*′*(5)^) = *a*_*i*_. Since *v*^*′*(5)^ has descendant leaves *v*^*′*(14)^ and *v*^*′*(15)^ such that *ℓ*^*^(*v*^*′*(14)^) ≠ *a*_*i*_ and *ℓ*^*^(*v*^*′*(15)^) ≠ *a*_*i*_, there is at least one migration (*v*^*′*^, *v*^″^) in the path from *v*^*′*(5)^ to *v*^*′*(14)^ or *v*^*′*(15)^ such that *ℓ*^*^(*v*^*′*^) = *a*_*i*_. Now by condition (iii) of Lemma 22, 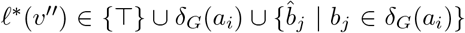. Thus, each of *n* gadgets *T* (*a*_*i*_) has a unique comigration from *a*_*i*_ to one of T, *b*_*j*_ or 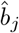 where *b*_*j*_ ∈ *δ*_*G*_(*a*_*i*_), which is not listed in Corollary 23. Let the set of these *n* distinct comigrations be 𝒜 Adding these comigrations from 𝒜 with the comigrations listed in Corollary 23, we get *C*(*T* ^*^, *σ*^*^, *ℓ*^*^) ≥ 2*n*^2^ + 6*n* + 1.

Second, we prove that for any optimal refinement (*T* ^*^, *σ*^*^, *ℓ*^*^) with *U* (*T* ^*^, *σ*^*^, *ℓ*^*^) = 10*n*^2^ +21*n* +2*m* unobserved clones, *M* (*T* ^*^, *σ*^*^, *ℓ*^*^) = 5*n*^2^ +8*n*+4*m* migrations, and *C*(*T* ^*^, *σ*^*^, *ℓ*^*^) = 2*n*^2^ +6*n*+1 comigrations, there is a unique perfect matching *K* such that (*T* ^*^, *σ*^*^, *ℓ*^*^) has a comigration from *a*_*i*_ to *b*_*j*_ for each edge (*a*_*i*_, *b*_*j*_) in *K*. To that end, let us consider the comigrations of the gadgets *T* ^*^(*b*_*j*_) with distinct leaf labels, {T *b*_*j*_ } ∪ *δ*_*G*_(*b*_*j*_) ∪ {*â*_*i*_ | *a*_*i*_ ∈ *δ*_*G*_(*b*_*j*_)} . Since node *v*^*′*(3)^ is labeled with *ℓ*^*^(*v*^*′*(3)^) = T and leaf *v*^*′*(10)^ is labeled with *ℓ*^*^(*v*^*′*(10^)) = *b*_*j*_, there is at least one migration (*v*^*′*^, *v*^″^) in the path from *v*^*′*(3)^ to *v*^*′*(10)^ such that *ℓ*^*^(*v*^″^) = *b*_*j*_. By condition (iii) of Lemma 22, *ℓ*^*^(*v*^*′*^) ∈ {T} ∪ *δ*_*G*_(*b*_*j*_) ∪ {*â*_*i*_ | *a*_*i*_ ∈ *δ*_*G*_(*b*_*j*_) } . Thus, each of *n* gadgets *T* ^*^(*b*_*j*_) has a comigration from one of T, *a*_*i*_ or *â*_*i*_ to *b*_*j*_, where *a*_*i*_ ∈ *δ*_*G*_(*b*_*j*_). Let the set of these *n* distinct comigrations be ℬ

Now (*T* ^*^, *σ*^*^, *ℓ*^*^) has (2*n*^2^ +6*n* +1) − (2*n*^2^ +5*n* +1) = *n* comigrations not listed in Corollary 23, which includes comigrations from 𝒜 and ℬ. Since each of 𝒜 and ℬ has *n* distinct comigrations, 𝒜 must be equal to ℬ But 𝒜 = ℬ is possible only if both includes comigrations only from *a*_*i*_ to *b*_*j*_ such that (*a*_*i*_, *b*_*j*_) ∈ *E*(𝒢). Let *K* be a subset of *E*(𝒢) such that (*a*_*i*_, *b*_*j*_) ∈ *K* if there is a comigrations from *a*_*i*_ to *b*_*j*_. Now by construction, 𝒜 = ℬ has exactly one comigration from each node *a*_*i*_ in *A*, and exactly one comigration to each node *b*_*j*_ in *B*, which implies each node *a*_*i*_ or *b*_*j*_ from *A* ∪ *B* is incident to exactly one edge in *K*. Therefore, *K* is a perfect matching for bipartite graph . 𝒢

Third, given a perfect matching *K*, we construct an optimal refinement (*T* ^*^, *σ*^*^, *ℓ*^*^) with *U* (*T* ^*^, *σ*^*^, *ℓ*^*^) = 10*n*^2^ + 21*n* + 2*m* unobserved clones, *M* (*T* ^*^, *σ*^*^, *ℓ*^*^) = 5*n*^2^ + 8*n* + 4*m* migrations, and *C*(*T* ^*^, *σ*^*^, *ℓ*^*^) = 2*n*^2^ + 6*n* + 1 comigrations that admits a comigration from *a*_*i*_ to *b*_*j*_ for each edge (*a*_*i*_, *b*_*j*_) in *K*. The steps are as follows.

1. Set the refinement (*T* ^*^, *σ*^*^, *ℓ*^*^) such that *T* and *T* ^*^ have the same topology and *σ*^*^ induces a bijection between the nodes of *T* and *T* ^*^. Let *v*^*′*(12)^ be the unique node such that *σ*^*^(*v*^*′*(12)^) = *v*^(12)^.
2. Following Lemma 22, set *ℓ*^*∗*^(*r*(*T*^′^)) = *ℓ*^*∗*^(*v*^*′*(0)^) = *ℓ*^*∗*^(*v*^*′*(1)^) = ⊤ and *ℓ*^*∗*^(*v*^*′*(2)^) = *ℓ*^*∗*^(*v*^*′*(3)^) = ⊥ for every gadget, and set both *ℓ*^*∗*^(*v*^*′*(4)^) and *ℓ*^*∗*^(*v*^*′*(5)^) to be 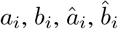, and *a*_*i*_ for gadgets *T* (*a*_*i*_, *â*_*j*_), *T* (*b*_*i*_, 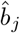), *T* (*â*_*i*_, *a*_*i*_), *T* (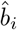, *b*_*i*_), and *T* (*a*_*i*_), respectively.
3. If edge (*a*_*i*_, *b*_*j*_) is in *K*, set *ℓ*^*∗*^(*v*^*′*(12)^) = *b*_*j*_ in gadget *T* (*a*_*i*_). Moreover, for any 1 ≤ *k* ≤ |*δ*_*G*_(*a*_*i*_)|, set 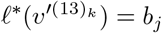 if 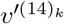 is labeled with *b*_*j*_, or set 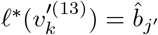 if 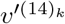 is labeled with *b*_*j*_^*′*^ *≠b*_*j*_. Similarly, if edge (*a*_*i*_, *b*_*j*_) is in *K*, set *ℓ*^*∗*^(*v*^*′*(4)^) = *ℓ*^*∗*^(*v*^*′*(12)^) = *a*_*i*_ in gadget *T* (*b*_*j*_). Moreover, for any 1 ≤ *k* ≤ |*δ*_*G*_(*b*_*j*_)|, set 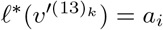 if 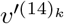 is labeled with *a*_*i*_, or set 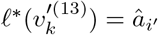 if 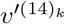 is labeled with *a*_*i*_*′* ≠ *a*_*i*_.

Clearly, (*T* ^*∗*^, *σ*^*∗*^, *ℓ*^*∗*^) is an optimal refinement with *U* (*T* ^*∗*^, *σ*^*∗*^, *ℓ*^*∗*^) = 10*n*^2^ + 21*n* + 2*m* unobserved clones, *M* (*T* ^*∗*^, *σ*^*∗*^, *ℓ*^*∗*^) = 5*n*^2^ + 8*n* + 4*m* migrations, and *C*(*T* ^*∗*^, *σ*^*∗*^, *ℓ*^*∗*^) = 2*n*^2^ + 6*n* + 1 comigrations and admits comigration from *a*_*i*_ to *b*_*j*_ for each edge (*a*_*i*_, *b*_*j*_) in *M* .

Finally, we prove bijection by showing the constructed refinement (*T* ^*∗*^, *σ*^*∗*^, *ℓ*^*∗*^) as the only refinement admitting a comigration from *a*_*i*_ to *b*_*j*_ for each edge (*a*_*i*_, *b*_*j*_) in *K*. Assume for contradiction that the optimal refinement (*T*^′^, *σ*^*′*^, *ℓ*^*′*^) with *U* (*T* ^*∗*^, *σ*^*∗*^, *ℓ*^*∗*^) = 10*n*^2^ + 21*n* + 2*m* unobserved clones, *M* (*T* ^*∗*^, *σ*^*∗*^, *ℓ*^*∗*^) = 5*n*^2^ + 8*n* + 4*m* migrations, and *C*(*T* ^*∗*^, *σ*^*∗*^, *ℓ*^*∗*^) = 2*n*^2^ + 6*n* + 1 comigrations, is different from (*T* ^*∗*^, *σ*^*∗*^, *ℓ*^*∗*^) and admits a comigration from *a*_*i*_ to *b*_*j*_ for each edge (*a*_*i*_, *b*_*j*_) in *K*. Since both (*T* ^*∗*^, *σ*^*∗*^, *ℓ*^*∗*^) and (*T*^′^, *σ*^*′*^, *ℓ*^*′*^) are optimal, they both follow the conditions mentioned in Lemma 22. As such, 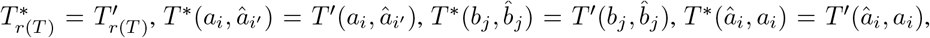, 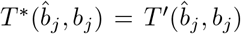 for any 1 ≤ *i, i*^*′*^, *j, j*^*′*^ *≤n*. Moreover, *ℓ*^*∗*^(*v*^*′*(0)^) = *ℓ*^*∗*^(*v*^*′*(1)^) = *ℓ*^*′*^(*v*^*′*(0)^) = *ℓ*^*′*^(*v*^*′*(1)^) = ⊤ and *ℓ*^*∗*^(*v*^*′*(2)^) = *ℓ*^*∗*^(*v*^*′*(3)^) = *ℓ*^*′*^(*v*^*′*(2)^) = *ℓ*^*′*^(*v*^*′*(3)^) = ⊥ in the refined gadgets of *T* (*a*_*i*_), and *ℓ*^*∗*^(*v*^*′*(0)^) = *ℓ*^*∗*^(*v*^*′*(1)^) = *ℓ*^*′*^(*v*^*′*(0)^) = *ℓ*^*′*^(*v*^*′*(1)^) = ⊤ in the refined gadget of *T* (*b*_*j*_) for any 1 ≤ *i, j* ≤ *n*. Thus, to show that (*T* ^*∗*^, *σ*^*∗*^, *ℓ*^*∗*^) and (*T*^′^, *σ*^*′*^, *ℓ*^*′*^) cannot be different and thus lead to contradiction, the following statements must be true.

i. For any node *a*_*i*_ in *A* where (*a*_*i*_, *b*_*j*_) ∈ *K* and for any 1 ≤ *k* ≤ |*δ*_*G*_(*a*_*i*_)|, it holds that 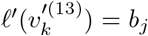 if 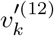 is labeled with *b*_*j*_, or 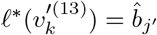, if 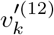 is labeled with *b*_*j*_*′ ≠b*_*j*_ in gadget *T*^′^(*a*_*i*_). Similarly, for any node *b*_*j*_ in *B* where (*a*_*i*_, *b*_*j*_) ∈ *K* and for any 1 ≤ *k* ≤ |*δ*_*G*_(*b*_*j*_)|, 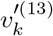 is labeled with *a*_*i*_ if 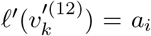, or 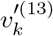 is labeled with *â*_*i*_*′* if 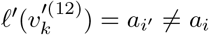 in gadget *T*^′^(*b*_*j*_).
ii. For any node *a*_*i*_ in *A*, 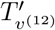 has only one node *v*^*′*(12)^ and is labeled with *a*_*i*_ in gadget *T*^′^(*a*_*i*_).
iii. For any node *b*_*j*_ in *B* where (*a*_*i*_, *b*_*j*_) ∈ *M*, *l*^*∗*^(*v*^*′*(4)^) = *ℓ*^*∗*^(*v*^*′*(12)^) = *a*_*i*_ in gadget *T*^′^(*b*_*j*_).

To prove (i), we consider the node *v*^*′*(13)^ from the gadget *T*^′^(*a*_*i*_) without loss of generality. Note that *v*^*′*(13)^ has two leaves with labels *b*_*j*_ and 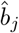, where *b*_*j*_ is the *k*-th element in *δ*_*G*_(*a*_*i*_).

So *v*^*′*(13)^ is labeled either with *b*_*j*_ or 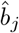. Now two cases can happen.

- (*a*_*i*_, *b*_*j*_) ∈ *K*: In this case, there must be a comigration (*u*^*′*^, *v*^*′*^) from *a*_*i*_ to *b*_*j*_ in the path from *v*^*′*(5)^ to *v*^*′*(14)^ in *T*^′^(*a*_*i*_) such that the nodes in the path from *v*^*′*^ to *v*^*′*(14)^ are labeled with *b*_*j*_. Since labeling *v*^*′*(13)^ with *a*_*i*_ admits new comigration from *a*_*i*_ to 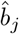 and thus makes (*T*^′^, *σ*^*′*^, *ℓ*^*′*^) suboptimal, *ℓ*^*′*^(*v*^*′*(13)^) = *b*_*j*_.
- (*a*_*i*_, *b*_*j*_) *∉ K*: In this case, there can only be comigration to *b*_*j*_ from 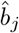 in *T*^′^(*a*_*i*_). Thus if 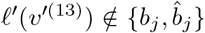, then (*T*^′^, *σ*^*′*^, *ℓ*^*′*^) has a new comigration to *b*_*j*_ and is suboptimal. On the other hand, if *v*^*′*(13)^ is labeled with *b*_*j*_, then there is a a migration (*u*^*′*^, *v*^*′*^) such that 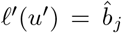 and *ℓ*^*′*^(*v*^*′*^) = *b*_*j*_ in the path from *v*^*′*(5)^ to *v*^*′*(14)^ in *T*^′^(*a*_*i*_). But that implies the two nodes *v*^*′*^ and *v*^*′*(14)^ labeled with *b*_*j*_ are disconnected, which means (*T*^′^, *σ*^*′*^, *ℓ*^*′*^) is suboptimal. Thus, 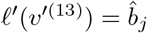.

Similarly, we can show that for any node *b*_*j*_ in *B* where (*a*_*i*_, *b*_*j*_) ∈ *K* and for any 1 ≤ *k* ≤ |*δ*_*G*_(*b*_*j*_)|, 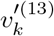 is labeled with *a*_*i*_ if 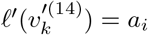, or 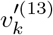 is labeled with *â*_*i*_*′* if 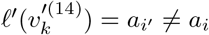 in gadget *T*^′^(*b*_*j*_). Therefore, (i) is true.

Next, we prove that for any node *a*_*i*_ in *A*, 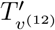 has only one node *v*^*′*(12)^ and is labeled with *b*_*j*_ in gadget *T*^′^(*a*_*i*_) where (*a*_*i*_, *b*_*j*_) ∈ *K*. To that end, we first prove 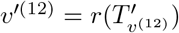 is labeled with *b*_*j*_. Note that *v*^*′*(5)^, the parent of the root of 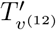, is labeled with *a*_*i*_. Since there is a comigration from *a*_*i*_ to *b*_*j*_, 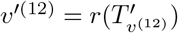 is labeled either with *a*_*i*_ or with *b*_*j*_. Assume *ℓ*^*′*^(*v*^*′*(12)^) = *a*_*i*_. Now two cases can happen.

- 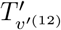 has only one node *v*^*′*(12)^: In this case, there are comigrations from *a*_*i*_ to 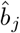, where (*a*_*i*_, *b*_*j*_) ∉ *M*, which are not in the 2*n*^*2*^ + 6*n* + 1 comigrations mentioned above. Thus (*T*, *σ*, *ℓ*) increases the number of comigrations and is not optimal.
- 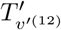 has at least two nodes: Since 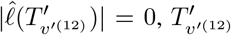 having two nodes increases the number of unobserved clones, making (*T*^′^, *σ*^*′*^, *ℓ*^*′*^) suboptimal.

Therefore, *ℓ*^*′*^(*v*^*′*(12)^) ≠ *a*_*i*_, and so *ℓ*^*′*^(*v*^*′*(12)^) = *b*_*j*_.

Next, we prove that for any node *b*_*j*_ in *B* where (*a*_*i*_, *b*_*j*_) ∈*K*, *ℓ*^*∗*^(*v*^*′*(4)^) = *ℓ*^*∗*^(*v*^*′*(12)^) = *a*_*i*_ in gadget *T*^′^(*b*_*j*_). Note that there is a comigration from *a*_*i*_ to *b*_*j*_ in *T*^′^(*b*_*j*_) if (*a*_*i*_, *b*_*j*_) ∈*M* . Since *ℓ*^*′*^(*v*^*′*(1)^) = ⊤, *ℓ*^*′*^(*v*^*′*(10)^) = *b*_*j*_, and *v*^*′*(4)^) is the only node between *v*^*′*(1)^) and *v*^*′*(10)^), *v*^*′*(4)^ must be labeled with *a*_*i*_ to admit the comigration from *a*_*i*_ to *b*_*j*_. Let 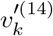 be the unique leaf in *T*^′^(*b*_*j*_) labeled with *a*_*i*_. Since 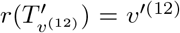 is in the path from *v*^*′*(4)^) to 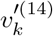 where both are labeled with *a*_*i*_, *v*^*′*(12)^ must be labeled by *a*_*i*_ to keep the number of migrations minimum. □

Therefore, (*T* ^*∗*^, *σ*^*∗*^, *ℓ*^*∗*^) = (*T*^′^, *σ*^*′*^, *ℓ*^*′*^).

## C MACH2 Integer Linear Program

We start by modeling (*R, e*) with binary variables **x** ∈ {0, 1}^|Σ|*×*|Σ|*×*(|*V* (*T*)|+|*E*(*T*))|^. Specifically, *x*_*s*, *t*, (*u*, *v*)_ = 1 if a refinement graph edge (*s, t*) is labeled by clonal tree edge *e*((*s, t*)) = (*u, v*) and *x*_*s*, *t*, (*u*, *v*)_ = 0 otherwise. Similarly, *x*_*s*, *t*, *v*_ = 1 if a refinement graph edge (*s, t*) is labeled by clonal tree node *e*((*s, t*)) = *v* and *x*_*s*, *t*, *v*_ = 0 otherwise. To ensure **x** encodes an edge-labeled refinement graph (*R, e*) for a pair 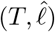 of clonal tree *T* and observed labeling 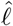, we impose constraints that model all four conditions of a refinement graph (Definition 9). Condition (i) of Definition 9 states that the subgraph *R*_*u*_ consisting of nodes *V* (*R*_*u*_) = {*s*|*e*((*s, t*)) = *u* or *e*((*s, t*)) = (*u, v*) or *e*((*s*^*′*^, *s*)) = *u* or *e*((*s*^*′*^, *s*)) = (*u*^*′*^, *u*) and edges *E*(*R*_*u*_) = (*s, t*) |*e*((*s, t*)) = *u* induces a directed tree without multi-edges in *R*. As this condition does not directly translate to ILP constraints, we resort to the following lemma.

### Lemma 25.

*For any node u* ∈ *V* (*T*), *the subgraph R*_*u*_ *induced in R is a directed tree without multi-edges if there is a root node s such that (i) s does not have any incoming edge* (*s*^*′*^, *s*) *in R*_*u*_, *i*. *e. e*((*s*^*′*^, *s*)) ≠ *u for any* (*s*^*′*^, *s*) ∈ *R; (ii) for any location t* ∈ *V* (*R*_*u*_) *where t*≠*s, there is exactly one incoming edge* (*t*^*′*^, *t*) *in R*_*u*_ *(i*. *e. labeled by e*((*t*^*′*^, *t*)) = *u); and (iii) there is a path from s to any location t* ∈ *V* (*R*_*u*_) *where t*≠*s in R*_*u*_.

*Proof*. (⇒) From the premise, *R*_*u*_ is a tree rooted at *s*. Then every node *t* of *R*_*u*_ that is not the root i. e. *t* ≠ *s*, has exactly one incoming edge (*s*^*′*^, *t*) in *R*_*u*_ labeled by *e*((*s*^*′*^, *t*)) = *u*, since *R*_*u*_ only contains edges labeled by *u*. For the same reason, the root *s* does not have any incoming edge labeled with *u*. Last, since *R*_*u*_ is a tree, there is a directed path from the root *s* to other non-root nodes *t* in *R*_*u*_.

(⇐) Assume that the conditions in the lemma are true. To prove that *R*_*u*_ is a tree rooted at *s*, we show that there is exactly one directed path from root *s* to each other node *t* in *R*_*u*_. Assume for a contradiction that there exists one node *t* in *R*_*u*_ such that there are at least two directed paths from *s* to *t* in *R*_*u*_. But then there exists at least one common node *s*^*′*^ in both the paths from *s* to *t*, which implies *s*^*′*^ has two incoming edges in *R*_*u*_, which contradicts with the premise. Thus, there is exactly one directed path from *s* to each other node *t* in *R*_*u*_, and so *R*_*u*_ is a tree rooted at *s*. □

Condition (ii) of Lemma 25 characterizes the non-root nodes *t* of *R*_*u*_ to be the ones with exactly one incoming edge in *R*_*u*_. More specifically, *t* is a non-root node in *R*_*u*_ if 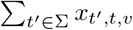 is 1. Moreover, condition (iv) of Definition 9 states that the root of *R*_*u*_ is *s* if *u* is not the root of *T* and there is an incoming edge (*s*^*′*^, *s*) in *R* labeled by *e*((*s*^*′*^, *s*)) = (*π*_*T*_ (*u*), *u*). In other words, *s* is the root of *R*_*u*_ if 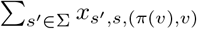 is 1. Note that Definition 9 does not characterize the root *R*_*r*(*T*)_ of *R*_*u*_ if *u* is the root of *T* . To that end, we introduce a new binary variable **w** ∈ {0, 1} ^|Σ|^ that models the root of *R*_*r*(*T*)_ such that *w*_*s*_ = 1 if the root of *R*_*r*(*T*)_ is *s* ∈ Σ, and *w*_*s*_ = 0 otherwise. The following constraint ensures that the root of *R*_*r*(*T*)_ is unique.

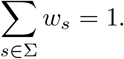

We enforce conditions (i) and (ii) from Lemma 25 requiring the root node *s* in *R*_*u*_ to have no incoming edges and each non-root node *t* to have exactly one incoming edge in *R*_*u*_ as follows.

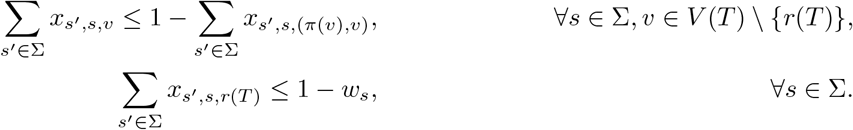

To enforce condition (iii) of Lemma 25, we introduce two new binary variables **y** = 0, 1 ^|Σ|×|Σ|×|*V*(*T*)|^ and **z** = {0, 1} ^|Σ|×|Σ|×|V(T)|^ to model the relations between any two nodes in *R*_*u*_ for any original node *u* ∈ *V* (*T*). More specifically, we require *y*_*s*, *t*, *u*_ to be 1 if there is a directed path from location *s* to *t* in *R*_*u*_, and *y*_*s*, *t*, *u*_ = 0 otherwise. The following constraints enforce *y*_*s*, *t*, *u*_ = 1 under this condition but do not explicitly enforce *y*_*s*, *t*, *u*_ = 0 otherwise.

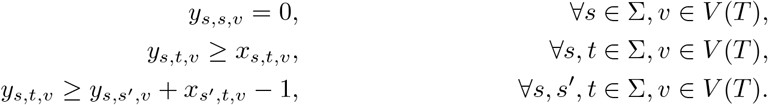

We require *z*_*s*, *t*, *u*_ to be 1 if *s* and *t* are in *R*_*u*_, but there is no directed path from *s* to *t* or from *t* to *s* in *R*_*u*_, and *z*_*s*, *t*, *u*_ = 0 otherwise. The following constraints set *z*_*s*, *t*, *u*_ = 1 under these conditions but do not explicitly enforce *z*_*s*, *t*, *u*_ = 0 otherwise.

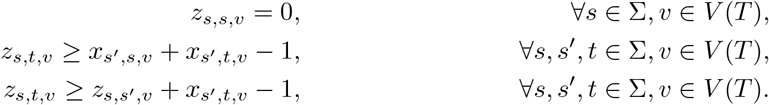

The following constraints enforce the entries in **y** and **z** to be 0 when applicable by requiring that two locations *s* and *t* are in *R*_*u*_ if and only if either there is a path from *s* to *t*, or from *t* to *s*, or there are no paths between *s* and *t* in *R*_*u*_.

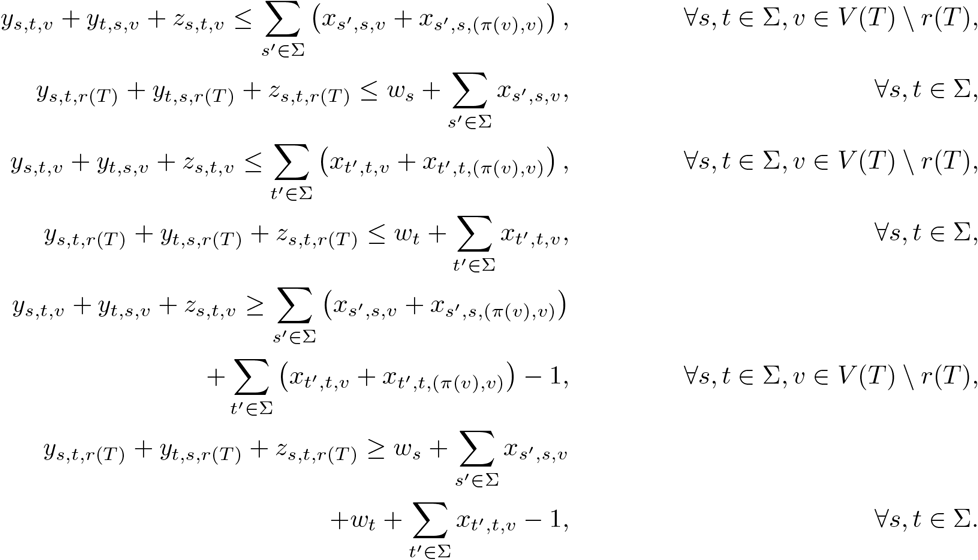

We enforce condition (iii) of Lemma 25 with the following constraints.

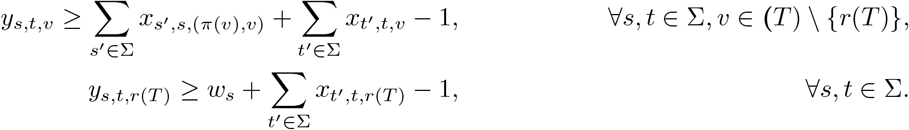

To conclude enforcing condition (i) of Definition 9, we require *s*^*′*^ to be in *R*_*u*_ for each edge (*s*^*′*^, *t*^*′*^) labeled by *e*((*s*^*′*^, *t*^*′*^)) = (*u, v*) where *v* is a child of *u*.

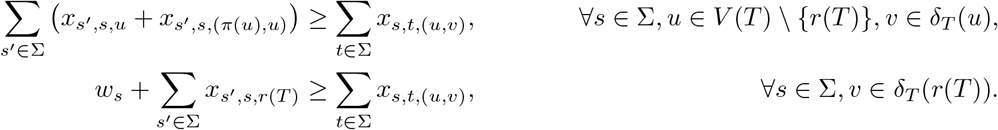

The following constraint ensures condition (ii) of Definition 9 by allowing exactly one edge (*s, t*) of *R* to be labeled by *e*((*s, t*)) = (*u, v*) for each edge (*u, v*) in *T* .

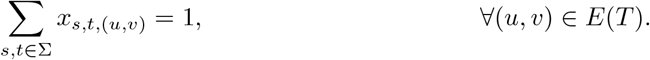

Last, we add the following constraints to model condition (iii) of a refinement graph *R* with edge labeling *e* from Definition 9, ensuring that for each pair (*v, s*) of node *v* ∈ *L*(*T*) and location *s* where 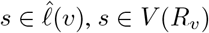.

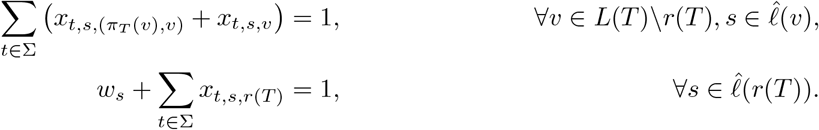

Note that condition (iv) of Definition 9 is already enforced.

For the objective function, we count the number of unobserved clones as follows.

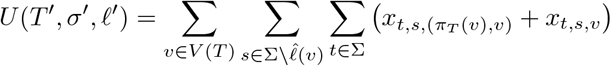

Next, we first compute the number of migrations. From Corollary 4, it follows that ℳ(*T*^′^, *σ*^*′*^, *ℓ*^*′*^) = {(*s, t*) ∈ *E*(*R*) | *s* ≠ *t*}. Thus, in ILP, we compute the number of migrations as follows.

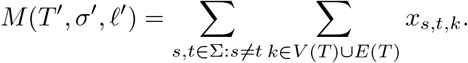

We do not compute the number of comigrations in the ILP, rather we use Parsimonious Consistent Comi-grations (PCC) algorithm [34] to compute the number of comigrations and eliminate solutions from the solution space that exceed the minimum. Based on the discussion above, we propose the following minimization functions for ILP.

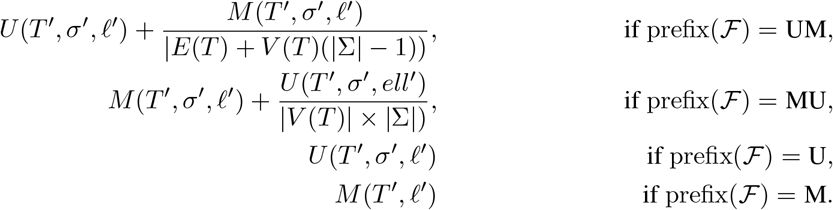

The factors |*E*(*T*) + *V* (*T*)(|Σ| − 1)) and 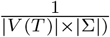 ensure the ILP performs lexicographical minimization following the specified criteria ordering ℱ. We state the full ILP in Section C.

### C. 1 Complete ILP

We present the ILP for ℱ = UM here.

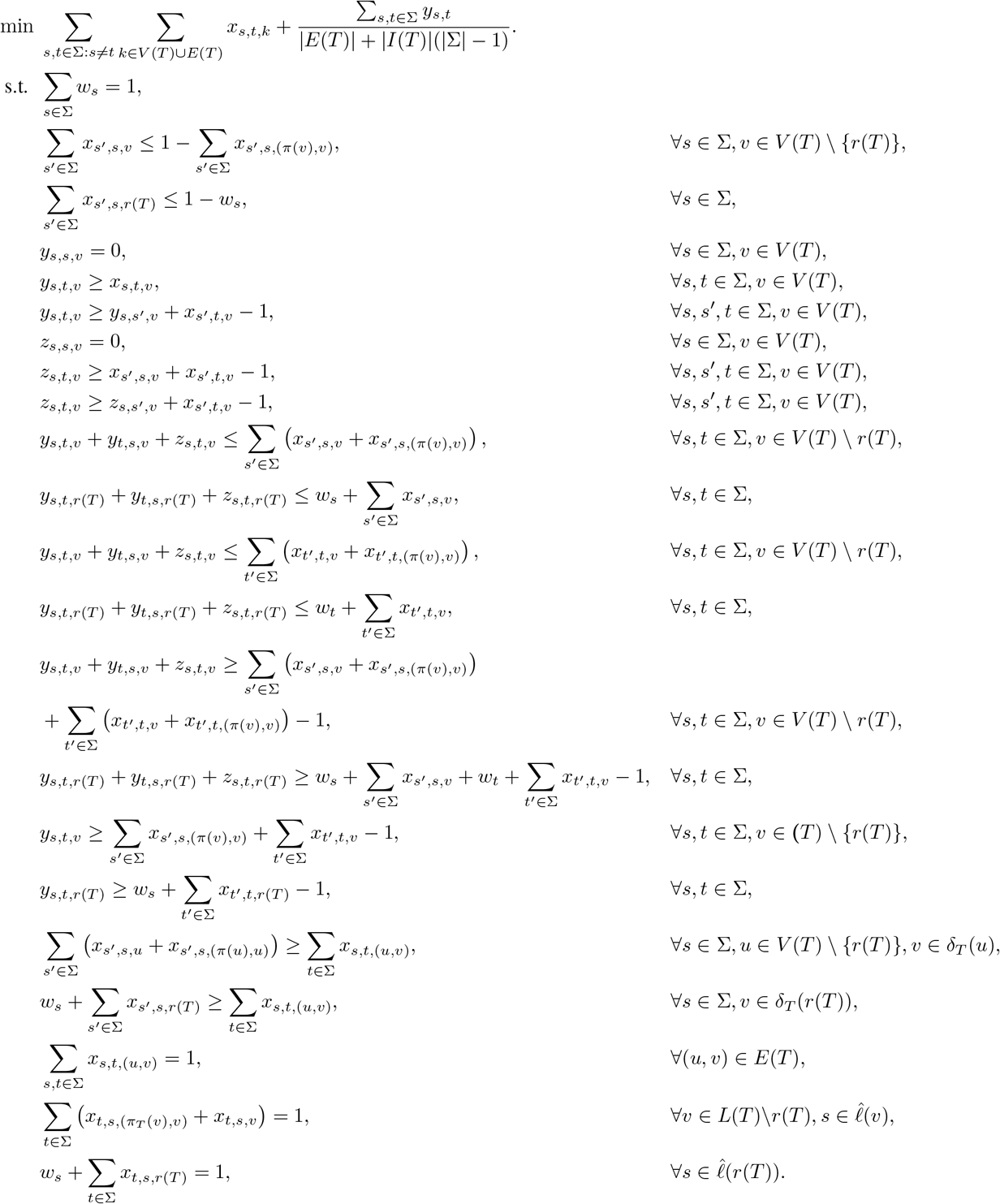

## D MACH2-Viz

MACH2-Viz contains the following features:

1. Visualization of corresponding nodes and edges between clonal phylogenies and migration graphs.
2. Visualization of the correspondence between input clonal trees and their refined polytomies.
3. Users can view observed locations of input clonal tree nodes, and unobserved clones in the output clonal tree.
4. Users can also compare and examine corresponding components between solutions.

The summary view (see Fig. S3) illustrates the summary graph. This view in the visualizer contains the following features:

1. Users can enforce priors such as known migrations and known absences of migrations to constrain the solution space.
2. If the solution space contains multiple possible primary tumors, the user can constrained to a known primary tumor, or rule out a primary tumor.

We also provide documentation^1^ to guide further open-source contributions to the visualizer. Specifically, we offer guidelines for integrating `mach2` results from your research into the `mach2-viz` landing page’s table of datasets. This systematic approach ensures a seamless and standardized process for the inclusion of new data.

## E Evaluation Metrics

We define three additional evaluation metrics that we use to compare MACH2 with MACHINA on simulated and real data.

### E. 1 Migration Graph *F*_1_ Score

To compare the solutions inferred by MACH2 and MACHINA with the simulated ground truth instances, we compute the *migration graph F*_1_ *score*, the harmonic mean of the precision and recall of migration edges. More precisely, for an inferred migration graph *G*^*′*^ and the corresponding simulated ground truth migration graph *G*^*^, we define recall and precision as the fraction of common edges *E*(*G*^*′*^) ∩ *E*(*G*^*^) in *G*^*^ and *G*^*′*^, respectively.

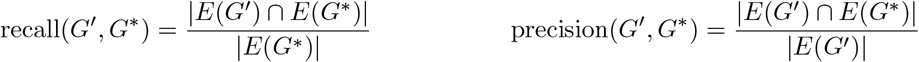

Note that since migration graphs are multi-edge graphs, the sets *E*(*G*^*′*^) and *E*(*G*^*^) of edges are also multisets, and thus consider multiplicity while computing intersection and cardinality. From precision and recall, we define the migration graph *F*_1_ score as follows.

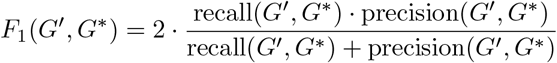

Migration graph *F*_1_ score ranges between 0 and 1, while *F*_1_(*G*^*′*^, *G*^*^) = 1 implying the migration graphs *G*^*′*^ and *G*^*^ to be exactly the same. Since MACH2 returns a set of solutions unlike MACHINA, we considered the median and minimum migration graph *F*_1_ scores among all the solutions returned by MACH2 (Section 5. 1)

### E. 2 Migrating Clone *F*_1_ Score

We define a *migrating clone* to be a refined node *u* in a solution (*T*^′^, *σ*^*′*^, *ℓ*^*′*^) such that there is a migration (*u, v*) ∈ *M* (*T*^′^, *σ*^*′*^, *ℓ*^*′*^). For an inferred set of migrating clones *R*^*′*^ and the corresponding simulated ground truth set of migrating clones *R*^*^, we define recall and precision as the fraction of common migrating clones *R*^*′*^ ∩ *R*^*^, respectively.

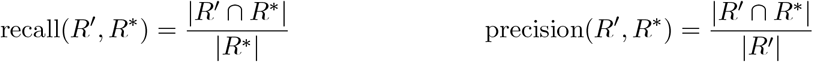

From precision and recall, we define the migrating clone *F*_1_ score as follows.

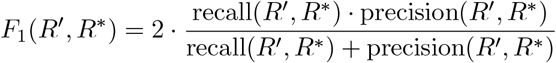

Migrating clone *F*_1_ score ranges between 0 and 1, while *F*_1_(*R*^*′*^, *R*^*^) = 1 implying the set of migrating clones *R*^*′*^ and *R*^*^ to be exactly the same. Since MACH2 returns a set of solutions unlike MACHINA, we considered the median and minimum migrating clone *F*_1_ scores among all the solutions returned by MACH2 (Section 5. 1)

### E. 3 Migration Pattern Totality Ratio (MPTR)

Methods like MACH2 and Metient often return multiple solutions, introducing uncertainty about whether a migration occurred. To quantify this uncertainty, we note that each solution (*T*^′^, *σ*^*′*^, *ℓ*^*′*^) induces a set *X*(*T*^′^, *ℓ*^*′*^) of ordered location pairs that correspond to migrations, i. e. *X*(*T*^′^, *ℓ*^*′*^) = {(*ℓ*^*′*^(*u*^*′*^), *ℓ*(*v*^*′*^)) | (*u*^*′*^, *v*^*′*^) ∈ *M* (*T*^′^, *ℓ*^*′*^)}. Given an enumerated space 𝒮 of solutions 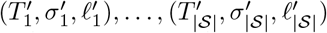, we define the *migration pattern to-tality ratio* (MPTR) as the fraction of inferred location pairs that are present in each enumerated solution, i. e. 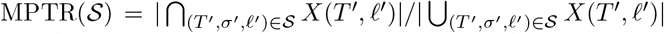. Thus, an MPTR value of 1 indicates that all inferred solutions agree on the migration pattern (modulo multiplicity), whereas a value of 0 means there is no agreement in the solution space on any specific migration between two locations.

^1^https://elkebir-group.github.io/mach2-docs/

## Notes

https://github.com/elkebir-group/MACH2/tree/main

### Competing Interest Statement

The authors have declared no competing interest.

### Summary of Updates

In the revised article we have added several changes and new analyses. First, we streamlined the text to emphasize that MACH2-UMC is the default and preferred ordering of parsimony objectives, this included simplifying Fig. 2 and the simulation section. Additionally, we provided more intuition and justification for several key concepts of the PMH-TR problem, including the new unobserved clones criterion, polytomy resolution and the summary graph. Finally, we included two new analyses showing the importance of enforcing temporal consistency as well as the use of the seeding location criterion as a post-processing step.

